# Modeling patient tissues at molecular resolution with Eva

**DOI:** 10.64898/2025.12.10.693553

**Authors:** Yufan Liu, Rishabh Sharma, Matthew Bieniosek, Amy Kang, Eric Wu, Peter Chou, Irene Li, Maha Rahim, Erica Bauer, Ran Ji, Wei Duan, Li Qian, Ruibang Luo, Padmanee Sharma, Renu Dhanasekaran, Christian M. Schürch, Gregory Charville, Aaron T. Mayer, James Zou, Alexandro E. Trevino, Zhenqin Wu

## Abstract

Tissue structure is essential to function and homeostasis in all organs, and disruptions to structure usually indicate disease. Modeling relationships between structural, molecular, and clinical aspects of tissues could advance new diagnostics and treatment strategies. Although profiling techniques like spatial proteomics can capture these relationships, the data remain challenging to extract insight from. Here, we present Eva, a foundation model for tissue imaging data that learns multi-scale spatial representations of tissues at the molecular, cellular, and sample level. Eva uses a novel vision transformer architecture and is pre-trained on masked reconstruction of over 40 million matched spatial proteomics and histopathology images. We show that Eva excels at a variety of tasks, including cross-modal inference from H&E to proteomics stains, quality control, data annotation, zero-shot retrieval, survival modeling, and patient stratification. Extensive evaluations on held-out validation data demonstrate the versatility and generalizability of the learned embeddings. We anticipate that Eva will accelerate translational science by bridging basic research and clinical practice.

## Introduction

Foundation models are deep learning models trained on massive, heterogeneous datasets with the goal of creating general-purpose representations that can be applied to diverse tasks. In biology, foundation model training frameworks have been established for several different types of data, including protein [1,2], DNA [3,4] and RNA sequences [5], single-cell transcriptomes [6], digital pathology [7–9], and microscopy images [10]. These models attempt to capture universal biological patterns or relationships. Here, we extend this paradigm to establish a visual model for multiplexed tissue imaging modalities. Our goal is to build a unified representation of tissue morphology and molecular phenotypes, enabling the exploration of relationships between these characteristics and real-world patient information. To achieve this, we generated and assembled a large repository of aligned spatial proteomics and histology (hematoxylin & eosin [H&E]-staining) imaging data. We then created and trained a model that can capture the complex covariances between diverse molecular biomarkers and spatial locations. Finally, we curated evaluations that showcase the relevance of this model to varied analytical and clinical tasks.

Prior work has extensively explored applications of self-supervised vision transformers (ViT) and masked autoencoders (MAE) [11–13] for routine histology images [8,14,15]. Extending these frameworks to multiplexed tissue imaging modalities, such as multiplexed immunofluorescence images, poses two additional challenges. First, each channel of the image directly encodes the location and intensity of a specific biomolecule, while the overall image further encodes morphology and the spatial interactions between these biomolecules. Second, the design of multiplexed imaging experiments varies widely in terms of the use of molecular probes and imaging platforms. Current efforts to model visual-biological datasets have begun to take these complications into account, but typically still accept a fixed number of channels as input [16], flatten channels and tokens during encoding [17,18], and/or focus on single technical modalities [19].

To advance a comprehensive visual model of tissue biology, we developed Eva, a self-supervised foundation model for tissue imaging data. Eva (“**E**ncoder of **v**isual **a**tlas”) introduces a novel two-stage hierarchical transformer architecture that learns relationships across channels and spatial domains. We pre-trained the model on multiplexed imaging data from over 4,000 tissue regions, 64 million cells, and around 200 protein biomarkers. Subsequently, using an extensive validation cohort of over 8,000 regions and 50 million cells, we showed that Eva outperforms existing models for multiplexed imaging on an array of tasks spanning scales of biology, including imputation of protein expression, image annotation and processing, cell and microenvironment classification, patient stratification, and clinical outcome predictions.

## Results

### Assembly of a pan-tissue multiplexed imaging atlas

To establish the training and validation data for Eva, we curated multiplexed imaging and aligned histology imaging data across tissues, platforms, and sources (**Methods**). The training subset comprised over 4,000 human tissue regions spanning diverse organs and molecular probes (**Figure 1A, Supplementary Table 1**), containing a mixture of tissue microarray cores and whole-slide sections. An additional validation cohort of over 8,000 regions was assembled for downstream evaluations, with 71% of these regions totally unseen during self-supervision training (**Supplementary Figure 1**). Most multiplexed images were derived from the CO-Detection by indEXing assay (CODEX/Phenocycler, 24%) or its commercial successor (PhenoCycler Fusion, 64%). Multiplexed ion beam imaging (MIBI) and Imaging Mass Cytometry (IMC) assays were also represented (1% and 11%, respectively; **Supplementary Figure 2A**). The vast majority of the data originated from human cancers (96%; **Supplementary Figure 2B**), collected from a range of research institutes and hospitals (**Supplementary Figure 2C**). Across all curated images, 66% had spatially co-registered H&E data prepared from either the same tissue slide or an adjacent section (**Supplementary Figure 2D**, **Methods**).

**Figure 1.**
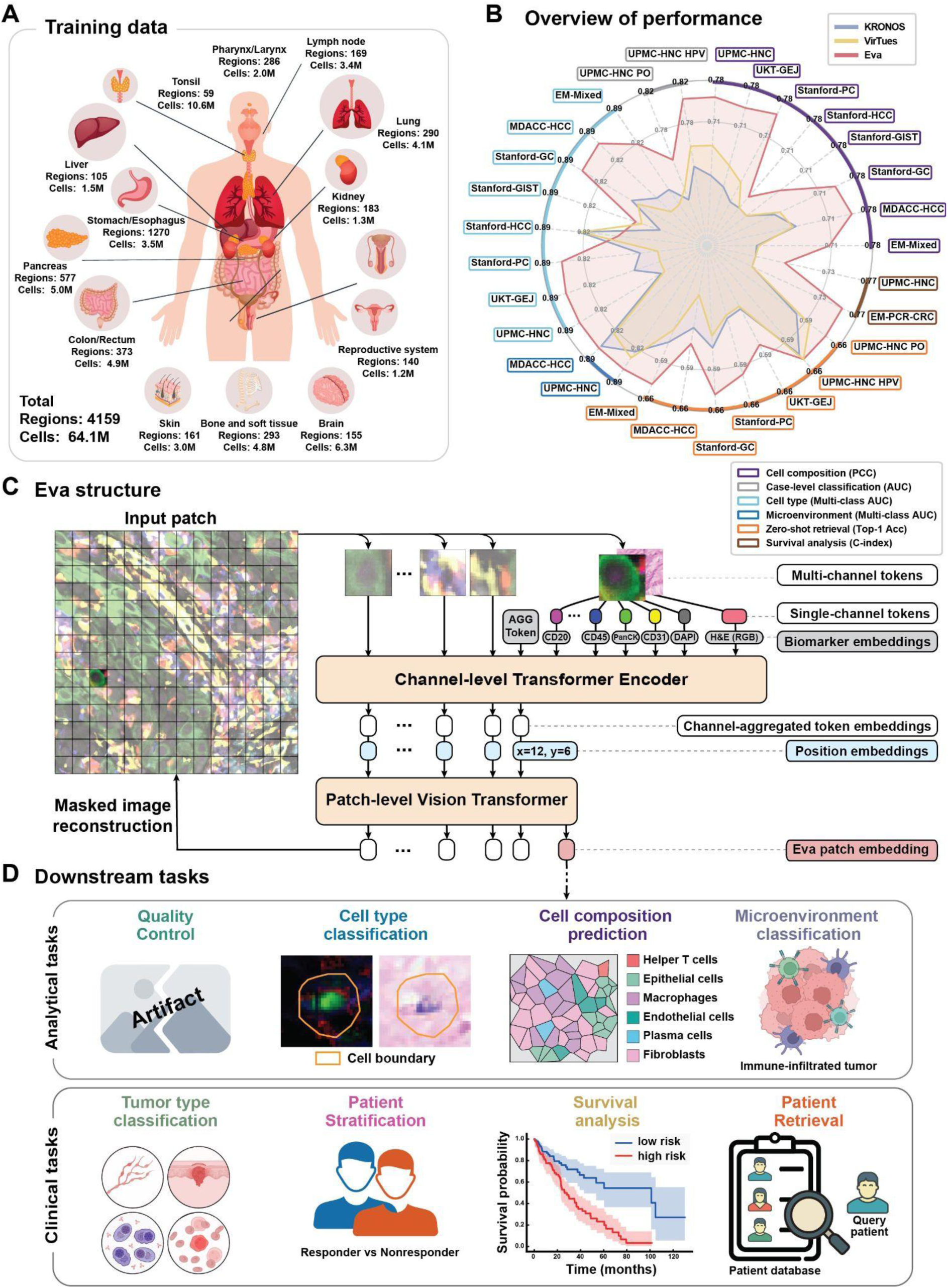
Overview of Eva. **A**, Datasets used for training Eva. **B**, Performance summary of Eva in comparison to related foundation models. The performance of VirTues and KRONOS was evaluated using their released pre-trained models. Axes are scaled according to the metric used. **C**, Schematic of Eva model structure showing the two-stage transformer architecture and self-supervised mask image reconstruction objective. Eva flexibly accepts any multi-channel image as input. **D**, The expressiveness of the Eva embeddings was assessed using many downstream tasks, including cell- and tissue-level tasks (i.e., quality control, cell type classification, cell composition prediction, and microenvironment classification) and clinical-level tasks (i.e., tumor type classification, patient stratification, survival analysis, and patient retrieval).

Our validation pipelines required detailed annotations at the cell, image, and patient levels. We therefore curated cell segmentation masks, biomarker expression tables, cell type annotations, neighborhood or structural annotations, expert reviews of image quality, and clinical label metadata across our validation cohort (**Figure 1B, D** and **Methods**).

### Encoding multi-scale, multi-channel tissue images

To develop a comprehensive visual model of tissue biology, we designed a deep learning architecture that: (1) handles image inputs with arbitrary combinations of channels, to account for the flexible panel design of multiplexed imaging experiments; (2) characterizes prior biological information of the biomolecules or stains; (3) models covariances between biomolecules encoded in the image channels; and (4) models spatial relationships at different scales to capture cell, neighborhood, and tissue region level information.

The resulting model is based on a masked autoencoder architecture. It employs a hierarchical, two-stage encoder-decoder framework to effectively learn both spatial and inter-channel covariances. The two major architecture components of Eva are: a lightweight channel encoder module [20] and a token-level Masked Autoencoder (MAE) Module. These parts operate in sequence to process and reconstruct tissue images (**Figure 1C** and **Methods**).

Briefly, the channel encoder module captures inter-channel covariances of the multiplexed images. This module adopted a channel-agnostic approach to accommodate arbitrary combinations of input channels [17]. In addition, to inject the prior biological knowledge of these biomolecules, we integrated their natural language embeddings [21] into the model (**Supplementary Table 1**). The token-level MAE module employs a vision transformer to learn the spatial relationships between tissue areas (**Figure 1C**).

### Image reconstruction with Eva

Eva was trained via masked image reconstruction. During training, multiplexed images were randomly selected and masked (random masking; **Methods**), and Eva was tasked to reconstruct the missing parts based on the visible contents (**Figure 2A–B** and **Supplementary Figure 3**). This strategy allowed Eva to generalize across arbitrary feature combinations without becoming overly reliant on a particular channel or spatial pattern.

**Figure 2.**
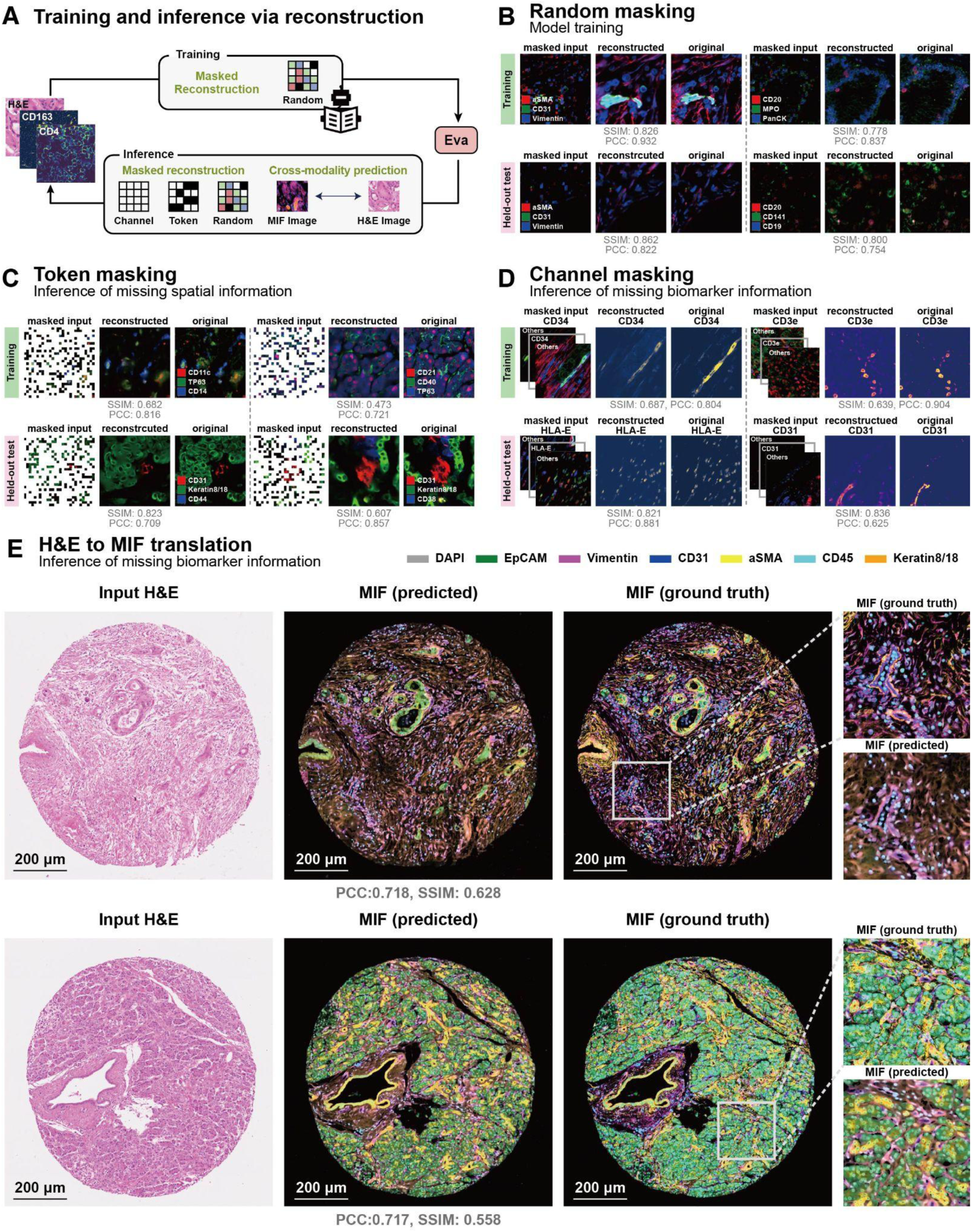
Multiplexed image reconstruction with Eva. **A**, Masking strategies used for training and evaluations of Eva. Random masking with a masking ratio of 0.75 was employed during Eva training. **B-D**, Illustration of reconstruction results and corresponding patch-level metrics (SSIM and PCC) under different masking strategies. **B**, Random masking reconstruction with a masking ratio of 0.75. **C**, Token masking reconstruction with a masking ratio of 0.75, evaluating the reconstruction of missing spatial contents. **D**, Channel masking with masked-channel input and CD34, CD3e, HLA-E, and CD31 reconstructions. This strategy evaluates inference of missing biomarker information. Signal intensity of the single channel is visualized using a pseudocolor scale. In each panel from **B** to **D**, the upper rows show patches sampled from the testing dataset used for Eva pre-training, and the lower rows show patches sampled from held-out validation datasets. **E**, H&E-to-MIF image translation using the fine-tuned Eva model (**Methods**). Two example tissue microarray cores are displayed, along with zoomed in views of selected windows. Seven key markers are shown. SSIM and PCC are calculated across the entire cores.

We also evaluated different masking schemes during inference to explicitly assess the model’s robustness in inferring missing spatial contents or biomolecule expression patterns (**Figure 2A** and **Methods**). “Token masking” masks all channels from randomly sampled spatial locations and “channel masking” masks entire channels. For each scenario, we presented inference examples from both training datasets and held-out validation datasets, illustrating that Eva can credibly reconstruct images across a spectrum of inputs (**Figure 2C–D**, **Supplementary Figures 4–5**).

To quantify this reconstruction performance, we calculated the Pearson correlation coefficients (PCC) and Spearman’s rank correlation between the original image and the reconstructed one on the masked tokens and channels. As an external reference, we employed the public model weights of a published architecture (VirTues [19]**, Methods**). Under both random and token masking strategies, Eva outperformed VirTues on held-out validation data, demonstrating higher PCC and Spearman’s rank correlation alongside lower MSE (**Supplementary Figure 6A**). We also present per-marker breakdowns for random and channel masking reconstruction performances (**Supplementary Figure 6B-C**), where Eva outperformed VirTues consistently across diverse biomarkers.

Tissue imaging encompasses diverse data modalities that often exhibit a trade-off between cost and information depth. A key example is the contrast between inexpensive, clinically ubiquitous H&E histology images and multiplexed immunofluorescence (MIF) data, which are substantially more information-rich but costly. Translating between modalities could unlock the ability to extract complex molecular and cellular features from cost-efficient assays [22–24] or to distill complex signatures into tractable clinical biomarkers. Eva, with its extensive self-supervised pre-training, can be easily finetuned and repurposed for these cross-modal translation tasks (**Methods**).

Here, we demonstrated this by predicting H&E images from MIF (**Supplementary Figure 7)**, and by reconstructing MIF from routine histology images (**Figure 2E** and **Supplementary Figure 8**). For the H&E-to-MIF translation task, we measured Eva’s overall and per-channel prediction performances on held-out validation data, and compared them against dedicated state-of-the-art approaches including ROSIE [22] and GigaTIME [23]. Across all primary evaluation metrics, including Spearman’s rank correlation, PCC, and structural similarity index measure (SSIM [25]). Eva consistently outperformed the other two models at both pixel- and grid-levels (**Supplementary Figure 9A, Methods**). Per-channel breakdown across biomarkers (**Supplementary Figure 9B-C**) demonstrated that Eva maintains a robust edge over GigaTIME and ROSIE while providing broader coverage of biomarkers. Visual comparisons supported the quantitative evidence (**Supplementary Figure 10**).

### Automated image and channel quality assessment

Quality control is an essential pre-processing step in tissue imaging analysis that helps to ensure reliable interpretation of the data. Eva, trained on a large collection of multiplexed tissue images, learned general-purpose representations that capture the normal distribution of imaging data. We investigated whether these learned representations can help us detect diverse imaging and staining anomalies. To do this, we curated three image quality tasks: (1) Naturalness Image Quality Evaluator (NIQE) predictions; (2) artifact detection via region-of-interest (ROI) annotations generated by human experts; (3) biomarker staining quality predictions using expert-generated numeric grades and detailed ROI-level staining reviews (**Methods** and **Figure 3A**).

**Figure 3.**
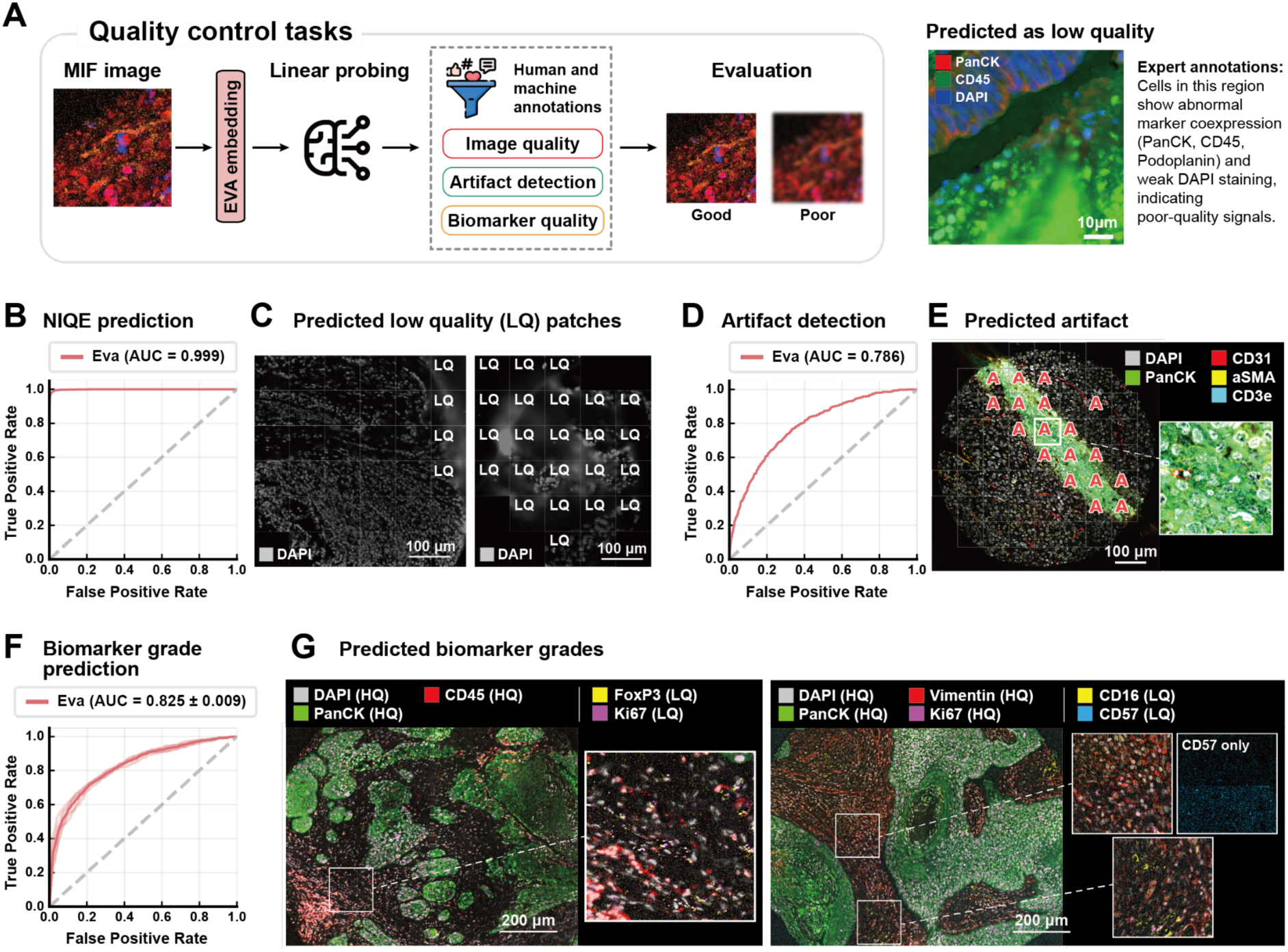
Image and biomarker quality assessment with Eva. **A**, Schematic of the quality control (QC) prediction tasks, including artifact detection, NIQE-based image quality assessment, and biomarker quality evaluation (**Methods**). All QC tasks were assessed by linear probing patch-level Eva embeddings. The right panel shows an example patch predicted as low quality, consistent with expert annotations. **B**, ROC curve and AUC for NIQE prediction by Eva. **C**, Example held-out test regions with predicted patch-level quality. LQ: low quality. **D**, ROC curve and AUC for artifact detection by Eva. **E**, An example held-out test region with a zoomed-in view of a patch with imaging artifacts. Patches labeled “A” in the region have high prediction scores for artifact presence. **F**, ROC curve and AUC (mean ± standard deviation) for biomarker quality assessment by Eva, evaluated using 5-fold cross-validation. **G**, Examples of biomarker quality assessment by Eva, with full-region and zoomed-in patch views. HQ and LQ denote high-quality and low-quality biomarkers predicted by Eva, respectively.

With a linear probing approach, Eva recapitulated NIQE quality scores almost perfectly (AUC = 0.999; **Figure 3B**). Visualizing the predictions showed that the model readily identified out-of-focus tissue areas (**Figure 3C** and **Supplementary Figure 11A**–**B**). Eva also accurately identified and localized a diverse set of human-annotated imaging artifacts caused by poor-quality staining, tissue folding, blurry imaging, or other issues (AUC = 0.786; **Figure 3D**). We illustrated this capability with several representative examples (**Figure 3E** and **Supplementary Figure 12**).

Another common quality control task in multiplexed imaging is to recognize and exclude channels with a poor signal-to-noise ratio or improper biomolecule staining. We showed that Eva can accurately identify these events and that its predictions aligned with human annotations and visual evidence (AUC = 0.825 ± 0.009; **Figure 3F, G**). On a held-out pan-tissue collection of expert-annotated staining grades, Eva also achieved accurate quality classification (AUC = 0.750 ± 0.008; **Supplementary Figure 13A**), again with clear alignment between Eva predictions and human-generated text descriptors (**Supplementary Figure 13B**). Together, these evaluations showed that Eva can be used to assess multiple dimensions of tissue imaging data quality.

### Automated cell labeling and label transfer

Reliable cell segmentation and classification of multiplexed imaging data is a critical yet often rate-limiting analysis step [26–28]. Automating this process could greatly advance our understanding of how cell states and neighborhoods vary across tissues. Identification of cell types is therefore a key task with which to evaluate a foundation model’s ability to learn and generalize key cell and tissue features [29].

To evaluate the performance of Eva for cell classification, we curated multiplexed imaging datasets from different sources, all with expert-generated cell type annotations. Reference models operating on the exact same multiplexed imaging data, including VirTues [19] and KRONOS [16], were used as comparisons.

We isolated single-cell images (**Methods**) and performed linear probing (**Figure 4A–B** and **Methods**) to predict cell types. Briefly, we extracted 32×32 pixel boxes around the centroids of cells, masked non-cell pixels, calculated embeddings, and applied the linear probe predictions. Eva achieved strong cell classification performance across datasets spanning 10 tumor types (**Figure 4C** and **Supplementary Table 2**), with multi-class AUC metrics ranging from 0.743 to 0.867. Eva achieved the best performance in 11 out of 12 datasets, with an average advantage of 11.2%. Notably, these validation datasets included multiplexed imaging data from diverse platforms (e.g., CODEX/Phenocycler, Phenocycler Fusion, IMC, MIBI), wherein Eva’s performance remained stable and top-class. We also reported F1-scores for each cell type class for two representative datasets (UPMC-HNC and Stanford-GC; **Figure 4D**).

**Figure 4.**
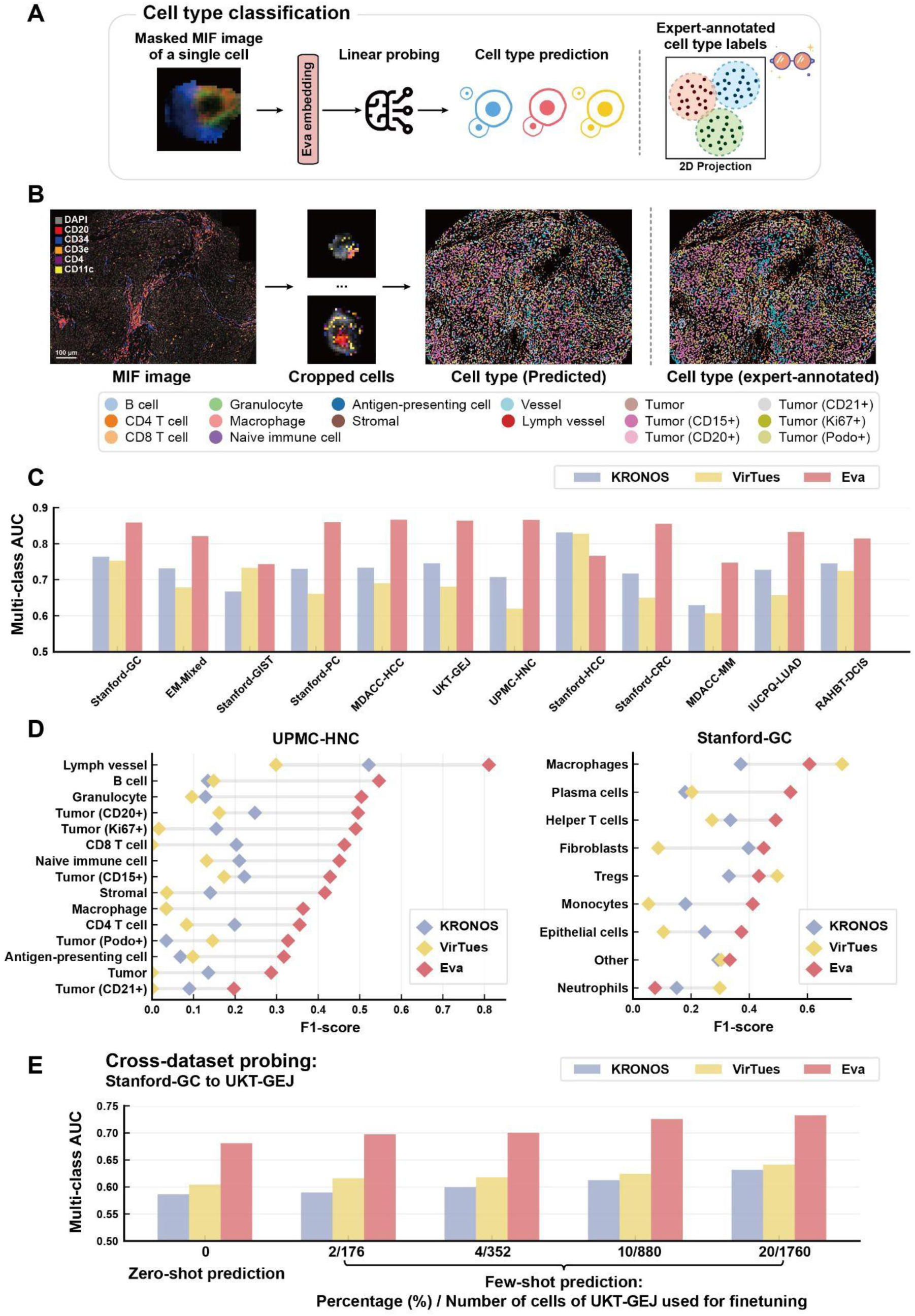
Automated cell type annotation and label transfer with Eva. **A**, Schematic of the cell type classification task. Eva takes a masked cell patch as input, in which only pixels within the segmented cell are visible (**Methods**). The embedding is then used for predictions of cell types via linear probing. **B**, Illustration of cell type annotation with Eva. Cell patches are extracted and annotated using Eva for every segmented cell. The predicted labels are visually compared with the original expert-annotated cell types. **C**, Bar plots comparing multi-class AUC performance of different foundation models on cell type prediction across 12 datasets of diverse tumor types. **D**, Detailed comparison of per cell type F1-scores in the UPMC-HNC and Stanford-GC datasets. **E**, Bar plots comparing multi-class AUC performance of cell type label transfer from the Stanford-GC to the UKT-GEJ datasets, under zero-shot and few-shot learning settings. Zero-shot learning directly applied the linear probe trained on Stanford-GC to UKT-GEJ. Few-shot learning settings included 2%, 4%, 10%, and 20% of cells sampled from the UKT-GEJ dataset for training the linear probe.

Cell states are frequently affected by their surrounding microenvironments. With this in mind, we tested an alternative processing approach that retained the full 32×32 boxes (box masking; **Supplementary Figure 14A**). All models benefited from additional spatial contexts, with AUC improvements between 0.009 and 0.023 (**Supplementary Figure 14B** and **Supplementary Table 2**).

We next evaluated model generalization in a challenging realistic analysis scenario: knowledge transfer from a well-annotated dataset to a new dataset having either no annotations (zero-shot) or minimal annotations (few-shot). We selected two datasets, Stanford-GC and UKT-GEJ, representing different classes of gastric cancer. We mapped cell type labels from both studies to unified classes (**Supplementary Figure 15**), trained a classifier using cell type annotations from Stanford-GC, with or without a minor subset of UKT-GEJ, and predicted on cells from UKT-GEJ. Eva achieved good performance across zero-shot (0%) and few-shot (2%, 4%, 10%, 20%) scenarios, maintaining performance advantage across all settings. All tested methods improved as more target data was used (**Figure 4E**).

### Microenvironment classification and cell composition prediction

The tumor microenvironment (TME) comprises diverse cell types that coordinate to promote, sustain, or regulate tumor development. The cell composition and genetics of the TME can vary widely even within an individual tissue sample [30], and the presence of particular microenvironments (e.g., tertiary lymphoid structures) have emerged as key indicators of prognosis and therapeutic outcomes [31]. In the digital pathology field, prediction of TME classes or quantification of TME characteristics has led to new diagnostic and therapeutic hypotheses [32,33]. The “unit” of a microenvironment, alternatively termed a neighborhood or a niche, has been defined in varying ways, but most current research agrees that multi-cellular neighborhoods provide biologically meaningful signals about the underlying tissue biology or disease state, beyond what can be gleaned from analyzing individual cells [34–37].

In this spirit, we used Eva to perform cell composition prediction and microenvironment classification, with neighborhood-sized image patches as inputs to a linear probing pipeline (**Figure 5A**). Each patch was assigned either a microenvironment class label [36] or the ground truth vector of cell type composition to set up evaluations of classification and regression tasks, respectively (**Methods**).

**Figure 5.**
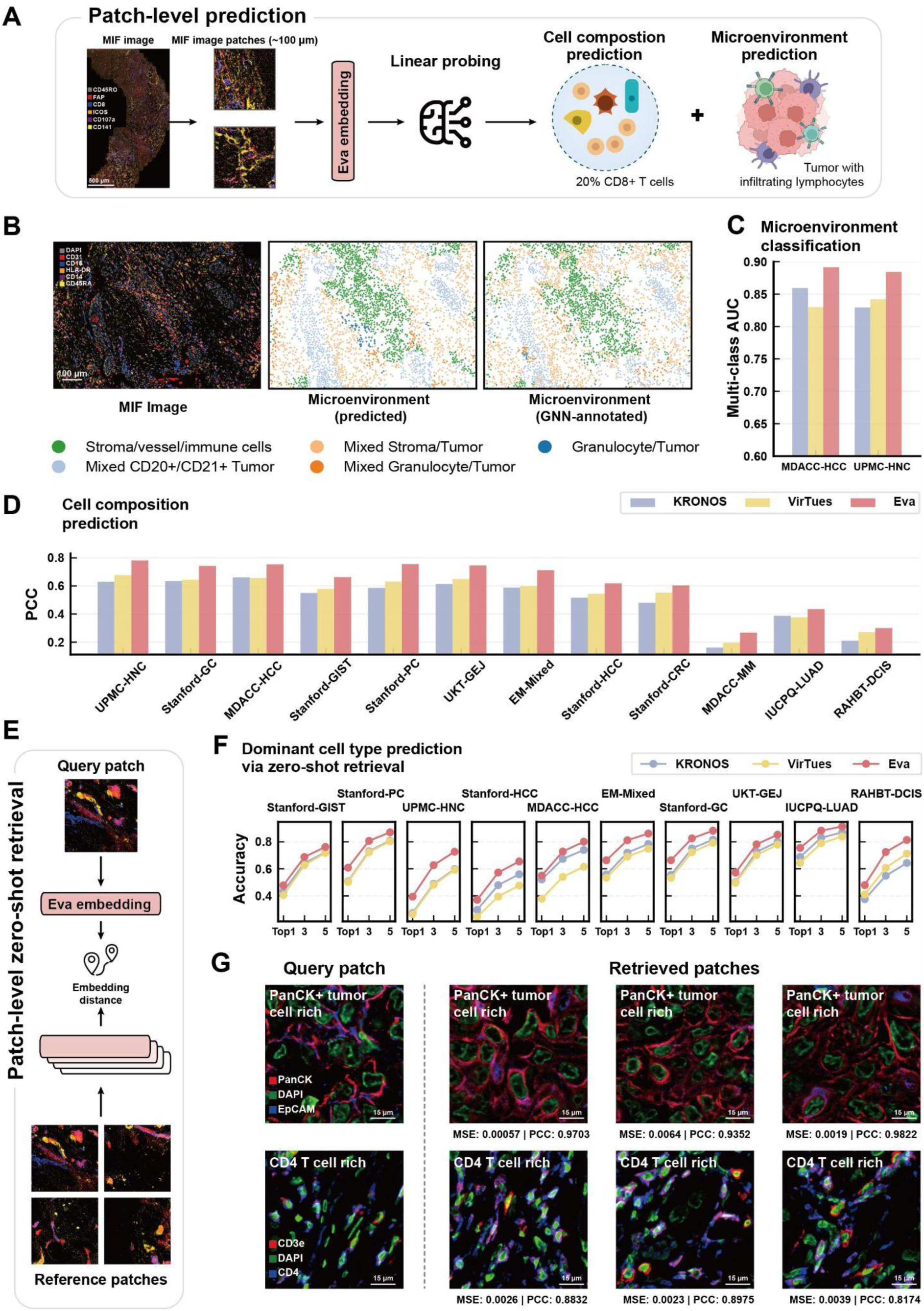
Characterizing cell composition and microenvironment class of neighborhoods with Eva. **A**, Schematic of the patch-level (i.e., neighborhood-level) tasks, where Eva embeddings of multiplexed image patches (∼100 μm) are projected via linear probing to assign a microenvironment label or to predict cell composition. **B**, Visualization of microenvironment prediction over an example region with Eva. Predictions were performed on patches centered on the labeled microenvironment centroid. The MIF image, the predicted microenvironment, and the original microenvironment annotations [36] are shown. **C**, Bar plots comparing multi-class AUC performance of different foundation models on predicting microenvironments from head-and-neck cancer (UPMC-HNC, defined for prognosis modeling) and liver cancer (MDACC-HCC, defined for immunotherapy response modeling). **D**, Bar plots comparing PCC and MSE performance of different foundation models on predicting cell composition. Higher PCC and lower MSE indicate better performance. **E**, Schematic of a zero-shot patch image retrieval framework. Distances between query and reference image patches are computed as cosine similarity between Eva embeddings. **F**, Comparison of different foundation models via the efficacy of image patch retrieval. We assessed whether top-1, top-3, and top-5 similar patches of the query patch share the same dominant cell type and report the accuracy metrics (**Methods**). Note that each dataset has ten or more cell type classes. **G**, Example query patches and their top-3 similar patches based on Eva embeddings. MSE and PCC of cell composition, as well as their dominant cell types are noted.

We present examples of microenvironment annotation maps generated by Eva, where both the classes and the spatial boundaries of different microenvironment archetypes were accurately identified (**Figure 5B** and **Supplementary Figure 16**). Eva annotated microenvironments in head and neck cancer (UPMC-HNC) and liver cancer (MDACC-HCC) tissue regions (**Figure 5C** and **Supplementary Table 3**) with high accuracy: AUC=0.884 and 0.892, respectively. Eva also inferred composition of cell types from neighborhood-sized input image patches. Across all 12 datasets evaluated, Eva achieved the highest correlations (PCC) and the lowest errors (MSE), with relative improvements from 10–36% and error reductions of up to 39% (**Figure 5D** and **Supplementary Table 4–5**).

### Patch-level image retrieval

A powerful application of foundation models is to retrieve similar tissue samples with shared biological and clinical characteristics [38]. Here, we assessed Eva’s embedding quality using a zero-shot, patch-level retrieval pipeline (**Figure 5E** and **Methods**). Eva retrieved patches with both similar biological properties (i.e., cell type compositions) and visual characteristics. Eva exhibited accurate top-1, top-3, and top-5 cell composition matching across validation datasets (**Figure 5F** and **Supplementary Table 6–7**). The retrieved patches were also visually aligned with the query patch in terms of biomarker distributions and cell morphology (**Figure 5G**).

The classification and retrieval of image patches can be affected by image resolution [39]. Therefore, we evaluated this task with upsampled and downsampled patch images, and Eva’s performance remained stable across studies and resolutions (**Supplementary Figure 17**).

### Tissue classification

A major goal in developing Eva was to make clinically relevant predictions, bridging real-world patient outcomes with the molecular, cellular, and morphological properties at play in producing disease. We therefore adopted an attention-based multiple instance learning (MIL) framework to pool image embeddings to the region (i.e., case) level [8,38,40,41] for various downstream tasks (**Figure 6A**).

**Figure 6.**
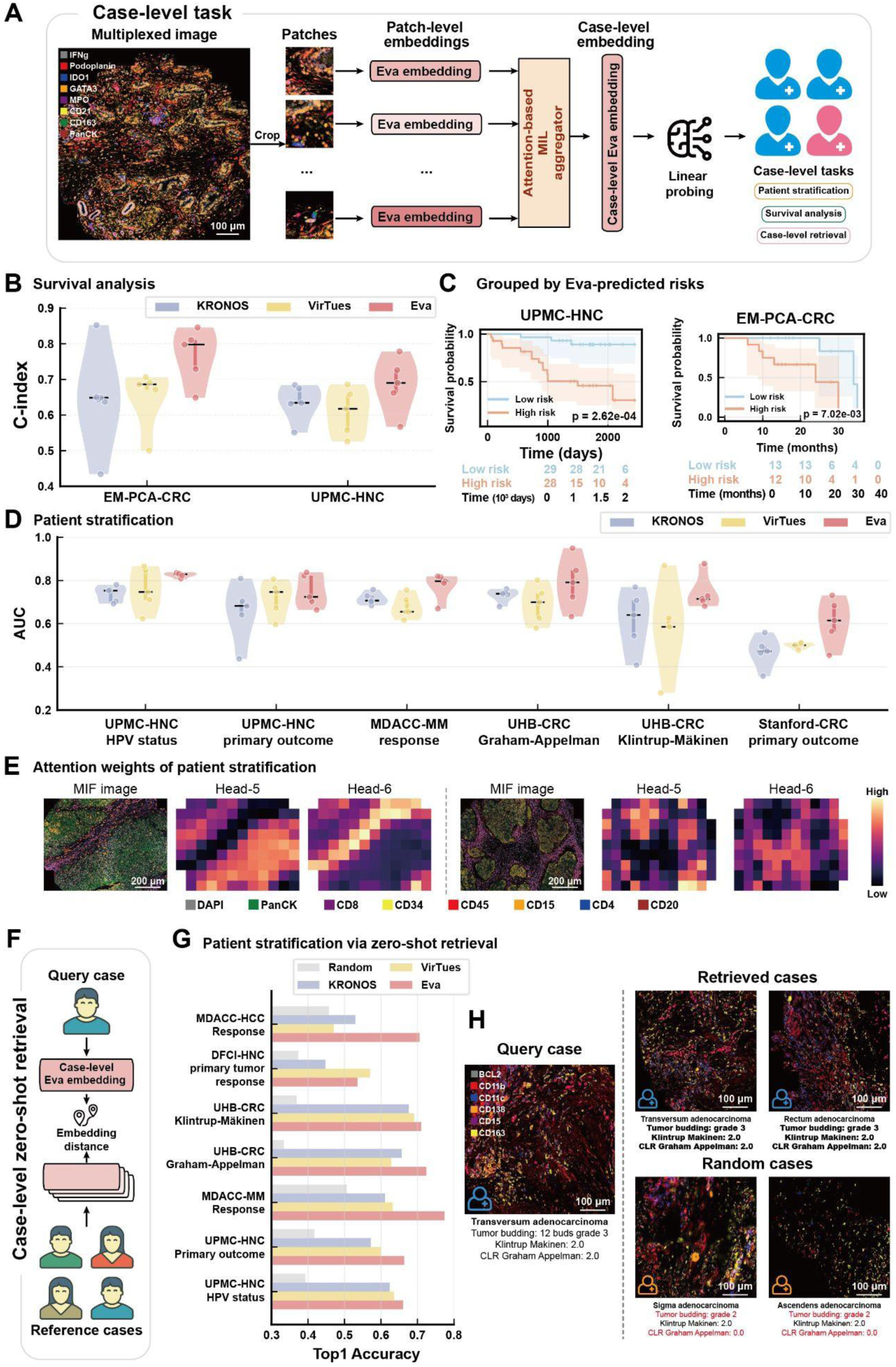
Modeling case-level clinical tasks with Eva. **A**, Illustration of the overall workflow for case-level tasks. Patches are cropped from a large multi-channel region, and their embeddings are aggregated using a multiple instance learning (MIL) approach to form a case-level embedding for each region. The case-level embedding is then projected via linear probing for downstream tasks. **B**, Model comparison of the concordance index (C-index) for survival analysis on the UPMC-HNC and EM-PCA-CRC dataset, evaluated using 5-fold cross-validation. **C**, Kaplan–Meier (KM) curve on the UPMC-HNC and EM-PCR-CRC dataset, with patients grouped by Eva-assigned risk scores. The numbers below the curve indicate, at each time point, the number of patients remaining at risk in the low-risk and high-risk groups. The p-value is calculated using a log-rank test comparing the survival distributions between the two groups. **D**, Model comparison for patient stratification, where each task is a binary classification and model performance is evaluated by AUC using 5-fold cross-validation. Descriptions of tasks are provided in **Methods**. **E,** Attention weights for patient stratification using MIL (**Methods**). MIL employed 8 heads, with the figure showing the 5-th and 6-th heads. **F**, Schematic of the case-level retrieval task, which is based on Eva embeddings aggregated from patch-level embeddings using MIL. Distances between query and reference embeddings are computed using cosine similarity. **G**, Model comparison of case-level retrieval using Top-1 accuracy. “Random” indicates a random baseline, where for each query case, a single case is selected at random from the reference set. **H**, Case studies of case-level retrieval. Each panel shows a multi-channel region with metadata describing the region displayed beneath. The left panel shows the query case. The upper right panels show the top two retrieved cases, and the lower right panels show two randomly sampled reference cases. Metadata different from the query case are shown in red.

We first performed cancer type and subtype classification, a task that relies on recognizing the different morphological characteristics [42–44] and molecular states [36,45] of TMEs. We assembled a dataset containing 2,500 regions from 9 broad cancer types (**Methods**). Each region was assigned a cancer type label, and predictions were derived via linear probing of MIL-aggregated embeddings under 5-fold cross validation. Here, we focused on embeddings of multiplexed imaging inputs computed by different models. All tested models were capable of distinguishing major cancer types, where Eva attained the best performance in 7 out of 9 tumor types and F1-scores ranging from 0.706 to 0.968 (**Supplementary Figure 18A** and **Supplementary Table 8**).

Next, we tested whether we could classify subtypes of lung cancer, a task closely relevant to treatment decisions in oncology settings. In lung cancer tissues, Eva consistently achieved the highest AUC value (0.949 ± 0.019) and F1-scores among MIF foundation models in distinguishing subtypes including adenocarcinoma (ADC), squamous cell carcinoma (SCC), and small cell lung carcinoma (SCLC) (**Supplementary Figure 18B** and **Supplementary Table 9**). Across these tasks, Eva’s performance proved largely comparable to predictions based on paired H&E inputs by pathology foundation models (PFMs) (**Supplementary Table 9**).

### Leveraging Eva for survival analysis

Having demonstrated that Eva learns expressive representations of multiplexed images, including biological contexts across scales and tissue types, we next aimed to use these capabilities for clinically-relevant modeling, including survival analysis, patient stratification, and similar patient retrieval. We curated tasks to systematically evaluate clinical utility in both supervised linear probing and zero-shot retrieval frameworks.

We first assessed if Eva could capture prognostic signatures in the TME. Survival analysis aims to quantitatively model patient risks over time, facilitating the design of follow-up strategies and clinical decision-making [46–49]. Survival outcomes, including event labels and time-to-event data, were curated from the clinical metadata of two validation datasets: UPMC-HNC and EM-PCA (a late-stage colorectal cancer cohort). Following the same approach, we derived MIL-aggregated case embeddings and used them to train a linear model using the Cox proportional hazards survival loss function [50,51]. Eva achieved a concordance index (C-index) of 0.685 ± 0.079 on the UPMC-HNC cohort and 0.766 ± 0.078 on EM-PCA, both representing the best performance among all compared models (**Figure 6B** and **Supplementary Table 10**). Similarly, Kaplan-Meier curves [46,47] stratified by Eva-derived risk scores exhibited the most pronounced separation between high-risk and low-risk groups on both datasets (**Figure 6C** and **Supplementary Figure 19**).

### Patient stratification and case-level retrieval

Next, we conducted patient stratification and clinical outcome prediction using diverse datasets and tasks (**Methods**), including HPV infection status, inflammation levels, binary prognostic outcome labels, and response to immune checkpoint inhibitor treatment. We performed predictive modeling through linear probing of the MIL-aggregated case embeddings, using 5-fold cross-validation and AUC metrics. Eva consistently achieved the highest accuracy and stability (**Figure 6D** and **Supplementary Table 11**).

We further visualized the attention weights from the MIL and highlighted different spatial distribution patterns. For example, head-5 primarily captured information from tumor area, while head-6 was activated on stroma (**Figure 6E** and **Supplementary Figure 20**), demonstrating that Eva learned biologically meaningful features that reflect the spatial organization of TMEs.

Furthermore, we evaluated if zero-shot retrieval based on case-level embeddings can enable query-based patient stratification directly, using the similarity between Eva-generated case embeddings without task-specific training (**Figure 6F** and **Methods**). This framework would provide a flexible and scalable approach for patient search in heterogeneous populations that takes molecular and microenvironment information into account. Eva consistently achieved robust top-1 retrieval accuracy, as measured by whether retrieved cases shared the identical metadata labels (**Figure 6G** and **Supplementary Table 12**). We visualized the top two retrieved cases for an example region, alongside randomly selected cases. Eva not only retrieved images that are visually consistent with the query case, but also samples which were similar at the metadata level (**Figure 6H**). Retrieved cases shared the same tumor budding grades, Graham-Appelman criteria, and Klintrup-Mäkinen score, indicating alignment of tumor inflammatory status.

Building on Eva’s standalone capabilities, we employed a late-fusion strategy to integrate Eva’s MIF embeddings with H&E embeddings generated by leading PFMs, specifically UNI [8] and Prov-GigaPath [52]. We reasoned that the structural and morphological contexts extracted by these very large-scale PFMs could be complemented by the molecularly-informed embeddings generated by Eva, leading to performance gains for patient stratification and survival modeling tasks. This integrative approach consistently transcends the performance limits of single-modality baselines (**Supplementary Figure 21** and **Supplementary Table 13**), highlighting Eva’s potential to enhance current histopathology models.

## Discussion

In this work, we introduce Eva (**E**ncoder of **v**isual **a**tlas), a foundation model for multiplexed tissue imaging data based on a novel two-stage masked autoencoder framework. Through large-scale self-supervised training on over 1.1 million multiplexed image patches (comprising 41.6 million single-channel patches), Eva learns to encode tissue biological contexts into compressed embeddings. Eva adopts a flexible and efficient architecture that incorporates semantic embeddings of protein/gene descriptions and can be readily extended to arbitrary combinations of biomolecules. Its compute-efficient design enables simultaneous, rapid encoding of hundreds to thousands of channels (**Supplementary Figure 22**).

Visually, Eva reconstructs masked images with high fidelity and exhibits strong cross-modality prediction performance. Eva also shows robust inference capability on downstream tasks at multiple scales (**Figure 1B**), including cell-, neighborhood-, tissue-, and clinical-level tasks, where it outperforms existing foundation models.

Although Eva achieves strong performance on held-out validation datasets for both reconstruction and downstream prediction, a key limitation is the restricted availability of out-of-distribution datasets, particularly for emerging technologies. Our current model and benchmarks rely heavily on data generated using the CODEX/Phenocycler platform, with limited validations performed on other platforms such as IMC and MIBI. This may bias evaluation of the generalization ability of Eva across diverse assay types and experimental conditions. Second, the labels used in downstream predictions—such as image quality annotations, cell types, and microenvironment classes—are curated by human experts, which could introduce bias into the evaluations. To mitigate this, we incorporated datasets from a variety of tumor types and sources to provide more comprehensive results. Nonetheless, the development of a gold-standard dataset would be invaluable for enabling more objective comparisons.

Generation of MIF images from H&E highlights the versatility of Eva. We anticipate that incorporating such generative objectives natively during training could further enhance Eva’s cross-modality prediction capabilities. Last but not the least, it is visually evident from the attention maps (**Supplementary Figure 20**) that Eva captures signals that align with known cellular and tissue compartments. However, how to systematically extract the mechanistic driving factors of patient-level characteristics, in the form of concrete molecular or cellular insights, remains an open question. Future work will leverage the generative capabilities of Eva to design integrated tools to decode its embeddings and reveal insights from patient stratification tasks.

Eva represents a step towards a cohesive analytical framework for the information-dense, high-dimensional multiplexed tissue imaging data, in which diverse assays and analysis tasks can be incorporated into a shared representation space. At the study level, Eva should immediately facilitate routine analysis tasks and cross-dataset knowledge transfer (via zero/few-shot learning) at different scales: from individual cells to patients, expanding the scope and depth of biological insights from spatial data. Eva will allow researchers to interrogate cellular, microenvironmental, and patient-level phenotypes within a single embedding space. On a larger scale, Eva could accelerate efforts to build a comprehensive catalog of spatial variation across human diseases, analogous to existing atlases for dissociated single cells [53] and pan-cancer genomics [54] by providing a common backbone for cross-study integration and comparison. More broadly, Eva adds to a growing body of literature on foundation models for biology, extending this paradigm to the rich but challenging regimes of multiplexed imaging and spatial omics.

## Methods

### Validation datasets

**Stanford-GC** contains 1,429 regions in tissue microarray (TMA) format from patients with various cancers, mainly of the pancreas, stomach, and esophagus, plus giant cell tumors and rhabdomyosarcomas, collected at Stanford Hospital. PhenoCycler Fusion multiplexed imaging (0.5 μm/pixel) with 54 biomarkers was performed by Enable Medicine, capturing 4.42 million cells (6,010 ± 3,931 per region). A subset of 473 regions have corresponding H&E, and were included in the training data. The remaining MIF-only regions were not used for training. We used this dataset for:

- Cell type and cell composition prediction on sampled cells and patches. Cell type labels were derived from Leiden clustering of biomarker expression, manually annotated and refined.

**EM-Mixed** contains 451 tumor regions in TMA format from various types of cancer patients (e.g., lung, kidney, cervix, ovary, lymph node, skin), collected from an assortment of archival tissue biorepositories. Phenocycler multiplexed imaging (0.3775 μm/pixel) with 46 biomarkers was performed by Enable Medicine, capturing 3.41 million cells (7,603 ± 8,893 per region). A subset of 231 regions have corresponding H&E, and were included in the training data. The remaining MIF-only regions were not used for training. We used this dataset for:

- Cell type and cell composition prediction on sampled cells and patches. Cell type labels were derived from Leiden clustering of biomarker expression, manually annotated and refined.

**Stanford-GIST** [22] contains 550 regions in TMA format from gastrointestinal stromal tumor (GIST) patients, collected at Stanford Healthcare. Phenocycler multiplexed imaging (0.3775 μm/pixel) with 54 biomarkers was performed by Enable Medicine, capturing 2.07 million cells (3,768 ± 2,769 per region). All regions have corresponding H&E and were used as part of the training data. We used this dataset for:

- Cell type and cell composition prediction on sampled cells and patches. Cell type labels were derived from Leiden clustering of biomarker expression, manually annotated, then propagated using kNN with top biomarkers.

**Stanford-PC** [22] contains 473 regions in TMA format from pancreatic cancer patients, collected at Stanford Healthcare. Phenocycler multiplexed imaging (0.3775 μm/pixel) with 54 biomarkers was performed by Enable Medicine, capturing 2.67 million cells (5,640 ± 2,226 per region). All regions have corresponding H&E and were used as part of the training data. We used this dataset for:

- Cell type and cell composition prediction on sampled cells and patches. Cell type labels were derived from Leiden clustering of biomarker expression, manually annotated, then propagated using kNN with top biomarkers.

**MDACC-HCC** [56,57] contains 50 regions from hepatocellular carcinoma patients, collected at MD Anderson Cancer Center (MDACC). Phenocycler multiplexed imaging (0.3775 μm/pixel) with 51 biomarkers was performed by MDACC, capturing 1.61 million cells (32,246 ± 15,490 cells per region). A subset of 30 regions with corresponding H&E on adjacent sections were used as part of the training data. We used this dataset for:

- Cell type and cell composition prediction on sampled cells and patches. Cell type labels were derived from Leiden clustering of biomarker expression, manually annotated, re-clustered, and refined.
- Patch-level prediction of microenvironment (neighborhood) annotations on sampled patches. Annotations were derived from a previous work [56].
- Case-level zero-shot retrieval for response to immune checkpoint inhibitor prediction (MDACC-HCC Response) on 25 pre-treatment regions with corresponding labels.

**UKT-GEJ** [22] contains 240 regions in TMA format from gastroesophageal junction carcinoma patients, collected at University Hospital Tübingen. PhenoCycler Fusion multiplexed imaging (0.5 μm/pixel) with 54 biomarkers was performed by Enable Medicine, capturing 366 thousand cells (1,524 ± 793 cells per region). All regions have corresponding H&E and were used as part of the training data. We used this dataset for:

- Cell type and cell composition prediction on sampled cells and patches. Cell type labels were derived from Leiden clustering of biomarker expression, manually annotated, then propagated using kNN with top biomarkers.

**UPMC-HNC** [36] contains 379 regions in TMA format from head-and-neck cancer patients, collected at University of Pittsburgh Medical Center. Phenocycler multiplexed imaging (0.3775 μm/pixel) with 43 biomarkers was performed by Enable Medicine, capturing 2.37 million cells (6,248 ± 3,889 cells per region). 327 regions have corresponding H&E from adjacent sections, and were used as part of the training data. We used this dataset for:

- Cell type and cell composition prediction on sampled cells and patches. Cell type labels were derived from Leiden clustering of biomarker expression and manually annotated and refined.
- Patch-level prediction of microenvironment (neighborhood) annotations on sampled patches. Annotations were derived from a previous work [36].
- Case-level prediction of HPV infection status (UPMC-HNC HPV), prognosis/primary outcome (UPMC-HNC PO), and survival modeling on 308 regions with corresponding labels.

**Stanford-CRC** [36] contains 406 regions in TMA format from colorectal adenocarcinoma patients, collected at Stanford Healthcare. Phenocycler multiplexed imaging (0.3775 μm/pixel) with 41 biomarkers was performed by Enable Medicine, capturing 746 thousand cells (1,836 ± 817 cells per region). This dataset does not have corresponding H&E, and is not included in the training data. We used this dataset for:

- Cell type and cell composition prediction on sampled cells and patches. Cell type labels were derived from Leiden clustering of biomarker expression and manually annotated and refined.
- Case-level prediction of prognosis/primary outcome (Stanford-CRC PO) on 196 regions with corresponding labels.

**MDACC-MM** [58] contains 564 regions in TMA format from melanoma patients, collected at MDACC. Imaging Mass Cytometry (1 μm/pixel) with 26 biomarkers was performed by MDACC, capturing 1.36 million cells (2,446 ± 1,814 cells per region). This dataset does not have corresponding H&E, and is not included in the training data. We used this dataset for:

- Cell type and cell composition prediction on sampled cells and patches. Cell type labels were assigned based on marker expression, using intensity thresholding for phenotype determination, validated by visual inspection.
- Case-level prediction of response to immune checkpoint inhibitor on 239 pre-treatment regions with corresponding labels.

**RAHBT-DCIS** [59] contains 79 regions from ductal carcinoma *in situ* (breast cancer) patients, collected from the Washington University Resource Archival Human Breast Tissue (RAHBT) cohort. Multiplexed ion beam imaging by time of flight (0.488 μm/pixel) data with 38 biomarkers was reported by Risom et al [59], capturing 69.7 thousand cells (882 ± 313 cells per region). This dataset does not have corresponding H&E, and is not included in the training data. We used this dataset for:

- Cell type and cell composition prediction on sampled cells and patches. Cell type labels were determined via FlowSOM clustering on MIBI-TOF marker expression of segmented cells, with iterative hierarchical subclustering and lineage assignment based on canonical marker combinations.

**IUCPQ-LUAD** [60] contains 533 regions from lung adenocarcinoma patients (both the discovery and validation cohorts), provided by the IUCPQ Biobank of the Quebec Respiratory Health Research Network. Imaging Mass Cytometry (1 μm/pixel) data with 35 biomarkers was reported by Sorin et al [60], capturing 3.13 million cells (5,872 ± 1,812 cells per region). This dataset does not have corresponding H&E, and is not included in the training data. We used this dataset for:

- Cell type and cell composition prediction on sampled cells and patches. Cell type labels were assigned via supervised hierarchical gating using binary pixel masks of lineage markers applied in rank order to segmented cells.

**Stanford-HCC** [61] contains 212 regions from hepatocellular carcinoma patients, collected at Stanford Healthcare. Phenocycler multiplexed imaging (0.3775 μm/pixel) with 41 biomarkers was performed by Enable Medicine, capturing 2.16 million cells (10,206 ± 7,398 cells per region). A subset of 35 regions has corresponding H&E, and was used as part of the training data. Other MIF-only regions were not included in the training data. We used this dataset for:

- Cell type and cell composition prediction on sampled cells and patches. Cell type labels were derived from iterative Leiden clustering on scaled marker expression, starting with major lineage markers and refined by subclustering subtype-specific panels.

**UHB-CRC** [37] contains 140 regions from advanced-stage colorectal cancer patients, collected at the University Hospital Bern. Phenocycler multiplexed imaging (0.3775 μm/pixel) data with 56 biomarkers was reported by Schürch et al [37], capturing 279 thousand cells (1,996 ± 890 cells per region). This dataset does not have corresponding H&E, and is not included in the training data. We used this dataset for:

- Case-level prediction of presence of CLR according to Graham-Appelman criteria (UHB-CRC GA) and Klintrup-Mäkinen score (UHB-CRC KM) on 140 regions with corresponding labels.

**DFCI-HNC** [36] contains 58 regions from head-and-neck cancer patients, collected at Dana-Farber Cancer Institute. Phenocycler multiplexed imaging (0.3775 μm/pixel) with 43 biomarkers was performed by Enable Medicine, capturing 137 thousand cells (2,357 ± 909 cells per region). This dataset does not have corresponding H&E, and is not included in the training data. We used this dataset for:

- Case-level zero-shot retrieval for response to immune checkpoint inhibitor prediction (DFCI-HNC pTR) on 58 pre-treatment regions with corresponding labels.

**EM-PanCancerAtlas (EM-PCA)** is a collection of pan-cancer tissue samples, containing over 9,000 regions. This work used a subset of 2,501 regions from 9 cancer types (e.g. breast, liver, kidney, color, rectum, lung, stomach, ovaries, and lymph nodes), collected from various archival tissue biorepositories. PhenoCycler Fusion multiplexed imaging (0.5 μm/pixel) with 54 biomarkers was performed by Enable Medicine, capturing 25.7 million cells (10,290 ± 6,333 per region). All regions have corresponding H&E. This dataset was not used in training. We used this dataset for:

- Case-level prediction of tumor types (N=2378), distinguishing between nine common cancer types (e.g., breast cancer / lung cancer).
- Case-level prediction of tumor subtypes within lung cancer (N=299, e.g., adenocarcinoma / small cell carcinoma).
- Case-level survival modeling of a collection of late-stage colorectal adenocarcinoma patients (N=123) underwent standard treatment (EM-PCA-CRC).

**Quality datasets** were collected from the Enable Cloud Platform (Enable Medicine). We curated 4 types of annotations for 3 imaging quality predictions:

1. The Natural Image Quality Evaluator (NIQE) score: NIQE is a no-reference image quality assessment metric (Saad & Bovik, 2012) that evaluates the perceptual quality of an image by comparing its statistical features to those derived from natural images. We derived region-level, channel-agnostic binary labels for MIF images based on NIQE scores computed on the DAPI fluorescence channel: low quality was defined as a NIQE score above 15.
2. Biomarker channel grades: We curated human expert-assigned biomarker quality grades generated using an interactive visualization tool (Enable Cloud Platform Visualizer). Grades ranged from 0-3: 0–a totally failed or incorrect stain; 1–significant staining issues but signal is present; 2–good quality stain; 3–excellent quality stain with high signal-to-noise ratio. We derived region-level, channel-specific binary labels for MIF images, defining low quality channels as those receiving a grade of 0.
3. Notes and stories (biomarker reviews): We curated human annotations of biomarker quality generated using an interactive visualization and annotation tool (Enable Cloud Platform Visualizer). These annotations contain text justifications and exemplary in-context regions of interest (ROIs, i.e., bounding boxes, lassos). They included both positive and negative assessments. ROIs were mapped to 224×224 patch images and processed into patch-level, channel-specific binary labels indicating low or high quality stains. These data were exclusively used to evaluate Eva-based prediction models trained on the labels derived from “Biomarker channel grades” (zero-shot).
4. Note and stories (artifacts): We downloaded and processed annotations of imaging artifacts, which were generated using an interactive visualization and annotation tool (Enable Cloud Platform Visualizer). Annotations come with detailed text justifications and exemplary in-context ROIs. From these ROIs, patch-level, channel-agnostic binary labels were derived indicating whether an imaging artifact was present. Positive patches annotated in this manner were paired with negative patches randomly sampled from the validation dataset to create a balanced dataset for artifact detection training and evaluation.

### Training dataset

Eva was trained to reconstruct multiplexed images with partial inputs, where paired H&E images were also included as morphology guidance. The training dataset consists of 4159 regions spanning diverse tissue types, phenotypes (both cancerous and non-cancerous), institutions, and biomarker panels (**Supplementary Figure 2**), collectively capturing approximately 64 million cells. Subsets of the validation datasets described above—such as Stanford-GC, Stanford-GIST, Stanford-PC, EM-Mixed, UPMC-HNC, and UKT-GEJ—that included paired H&E formed major portions of the training data. Additional multiplexed images were included to further augment training diversity. In total, the training dataset contains around 1.1 million non-overlapping image patches (224×224), or around 41.6 million single-channel images.

### Data processing

Multiplexed images of tissue regions were first divided into non-overlapping patches with spatial dimensions of 224×224 pixels, a standard practice for computer vision datasets [62]. Each pixel represents a width ranging from 0.3775 to 1 μm. Each cropped patch is a multi-channel image with *C* channels, where each channel *c*_!_ corresponds to a specific biomarker. The paired H&E images are incorporated as additional channels, resulting in a total of *C* + 3 channels. Names of all biomarkers in the dataset are listed in **Supplementary Table 1**.

Input MIF images (representing protein expression signals) were clipped and normalized to 0-1 scales using lower and upper bounds derived from histogram thresholding of the full region. H&E images were first scaled to a 0-1 range by dividing pixel values by 255 (for unsigned 8-bit integer format), and then inverted by computing 1 − *I*, where *I* represents the scaled pixel values. This inversion generates black-background images that are more consistent with fluorescence images, improving training efficiency.

### Eva architecture

Eva, a novel masked autoencoder (MAE) architecture, is specifically designed for multiplexed tissue imaging data. It employs a hierarchical two-stage framework to effectively capture both inter-channel and spatial relationships within the MIF data. It comprises two main components: a lightweight Channel-level Encoder [20] and a Token-level Masked Autoencoder module, which work in sequence to process and reconstruct MIF and H&E data. The model processes input images of shape *X* ∈ ℝ*^B^*^×*C*×*H*×*W*^, where *B* represents the batch size, *C* denotes the number of channels, and *H*, *W* represent height and width of the input patch. In this section, we will describe the components of Eva and the masking strategies employed during model training. To maintain consistency in terminology, we refer to the cropped image extracted from the original imaging experiments, which has a spatial dimension of 224×224, as the “patch.” The tokenized image used for model processing, which has a spatial dimension of 8×8, is referred to as the “token” (**Supplementary Table 14**).

#### Channel-level Encoder

The Channel-level encoder acts as the initial stage of the two-stage transformer encoder, tasked with capturing relationships between various channels, including both MIF protein biomarkers and H&E channels. It begins by dividing the input patch images into non-overlapping multi-channel tokens and processing them with a channel-agnostic embedding module, where each token is processed independently using a shared convolutional kernel [17]. The channel-agnostic approach allows for arbitrary input channel combinations, making it particularly suitable for multiplexed image inputs.

In detail, the channel-level encoder divides input patches of shape *C* × *H* × *W* into non-overlapping *C* × *P* × *P* multi-channel tokens. For each token, we further divide it into a set of 1 × *P* × *P* single-channel tokens, each of which undergoes a dedicated 2D convolutional layer, resulting in a tokenized and embedded representation:

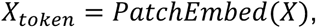

here *X_token_* ∈ ℝ*^B^*^×*N*×*C*×*P^2^*^ is the output of the initial patch embedding module, where *N* = (*W*/*P*) × (*H*/*P*) is the number of tokens cropped from the input patch. *P* is the spatial dimension of the token, which is set to 8 in Eva (**Supplementary Table 14**). The Channel-level Encoder ensures that each channel is processed independently while sharing the same convolutional weights, thereby maintaining computational efficiency as the number of channels scales.

After embedding, each single-channel token is further projected to a higher-dimensional space using a multi-layer projection network. Next, we integrate the biomarker information into the single-channel token embeddings using biomarker embeddings. Similar embeddings have previously been derived using different strategies: Evolutionary Scale Modeling (ESM) embedding of protein sequences adopted by VirTues [19], or plain positional encodings with learned representations adopted by KRONOS [16]. In our approach, we adopted language model-based biomarker embeddings generated by GenePT [21], which uses large language models (LLMs) to encode textual summaries of gene/protein functions. We find GenePT embeddings to be better at modeling realistic biological functions of biomarkers. For biomarkers that directly map to proteins, we extract their embeddings from a lookup table, and process them through a linear projection layer to match dimensions. For non-protein biomarkers including DAPI and 3 H&E channels, we randomly initialize a learnable embedding to each channel during training. The two embeddings are summed, concatenated with an aggregation token, and fed into a multi-head attention module, which integrates information within the multi-channel token along the channel axis *C*. Formally,

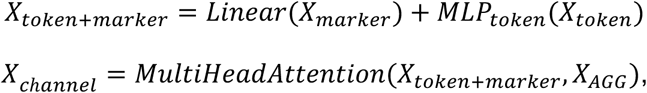

where *X_marker_* ∈ ℝ*^B^*^×*N*×*C*×*3072*^ contains all biomarker embeddings extracted from GenePT or randomly initialized, *X_AGG_* ∈ ℝ*^B^*^×*N*×*1*×*D*^*_channel_* is the additional aggregation token. *X_channel_* ∈ ℝ*^B^*^×*N*×(*C*+1)x*D_channel_*^ is the final output of the Channel-level Encoder, where the sequence of *C* + 1 embeddings correspond to *C* biomarker channels and an additional aggregation token, and *D_channel_* represents the embedding dimension. we extract corresponding output of aggregation token 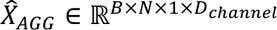 as a compressed representation of the input and fed it to the subsequent token-level MAE.

#### Token-level Masked Autoencoder

The Token-level MAE constitutes the second stage of the encoder and the decoder module, responsible for capturing spatial relationships between tokens. This model receives the channel-level aggregation token embeddings and performs a standard vision transformer-style (ViT) encoding [13] to capture and integrate information from different spatial locations. Concretely, the aggregation token embeddings are combined with learnable sinusoidal positional encodings to preserve spatial information [20], and further processed using an encoder with standard vision transformer blocks (along the token axis *N*), each containing multi-head self-attention mechanisms and feed-forward networks:

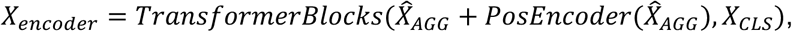

where *PosEncoder* generates the positional encoding for *X_channel_*, *X_CLS_* is an additional CLS token appended at the end of the sequence. Output of the token-level encoder is denoted as the token-sequence embedding *X_encoder_* ∈ ℝ*^B^*^×*N*(*N*+1)x1x*D_enc_*^, where *N* + 1 corresponds to the *N* tokens and an additional CLS token, and *D_enc_* represents the dimension of the token-sequence embedding. Notably, the channel axis is reduced to 1 as only the aggregation token embeddings are used.

Token-sequence embeddings will then be passed to the decoder module for image reconstruction. The decoder reconstructs both channel and spatial information from a compressed representation derived from the two-stage transformer. First, the token-sequence embedding is expanded along the channel axis to form *X_expand_* ∈ ℝ*^B^*^×(*N*+1)×*C*×*D_enc_*^. Then, the expanded embedding is combined with the biomarker embeddings used in the channel-level Encoder, which signifies information of different channels, and decoder positional embeddings:

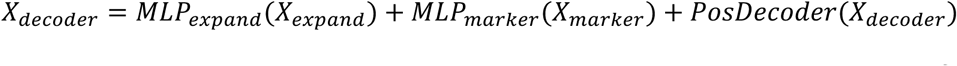

Subsequently, this decoder input is passed to the decoder blocks, which consist of several lightweight ViT blocks. Since the number of channels varies across batches, the decoder is only applied along the token dimension. Finally, the output is projected back to the original patch dimensions, allowing for pixel-level reconstruction of the input patch with both MIF and H&E channels:

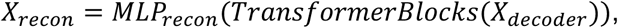

*^X^_recon_* ∈ ℝ*B*×*C*×*N*×^P^2^^ represents the reconstructed multi-channel image. We used mean square error (MSE) as the reconstruction loss function to minimize the pixel-level distance of input images and reconstructed images.

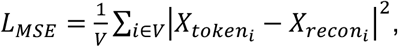

where *V* denotes the set of masked tokens, meaning we only compute the loss over the masked positions.

#### Embedding usage

The embeddings generated by Eva’s two-stage transformer encoder were used for downstream evaluations. For image patches, we computed the average of all token embeddings generated from the encoder of Token-level Masked Autoencoder within a patch to acquire its overall representation:

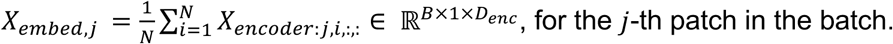

Region-level embeddings were derived by aggregating embeddings of patches from the region using a multiple instance learning (MIL) approach. MIL treats each object/region as a bag of instances/patches and learns to aggregate instance-level features into a bag-level representation through various pooling strategies. Concretely, we applied gated multi-head attention-based MIL mechanism [41]. For a bag of *N* patch embeddings denoted as *X_embed_* = {*X_embed_*_,1_, *X*_*embed*,2_ …, *X_embed_*_,*N*_}, the multi-head attention score and region-level aggregated embedding can be represented as:

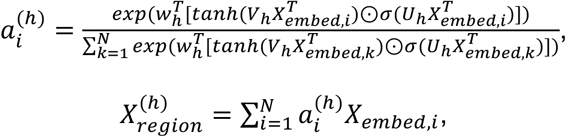

where *V* ∈ ℝ*^H^*^×*L*×*D*^and *U* ∈ ℝ*^H^*^×*L*×*D*^denote the transformation matrices that map the input embedding into the attention space with dimension *L*, where *H* is the number of heads in the attention block. *w* ∈ ℝ*^H^*^×*L*^ is the attention weight vector. The region-level embedding for each head is computed as the weighted sum of the input patch-level embeddings, where the attention score *a_i_* for the *i*-th patch in the region serves as the attention weight. The region-level embeddings from all attention heads are then concatenated to form the final region-level embedding.

#### Masking strategies

We implemented flexible masking mechanisms with multiple strategies to evaluate the robustness and different capabilities of Eva. Concretely:

- Random masking randomly masks single-channel tokens; i.e., tokens at different spatial locations will have different channels masked. This strategy is used to train Eva.
- Token masking randomly selects tokens and masks all channels of these selected tokens. This strategy focuses on evaluating Eva’s capabilities in capturing spatial relationships.
- Channel masking randomly selects channels and masks them from all tokens. This strategy focuses on evaluating Eva’s capabilities in capturing inter-biomarker dependencies.
- H&E masking masks out all H&E channels for all tokens. MIF masking masks out all MIF channels for all tokens. These two strategies focus on evaluating Eva’s cross-modality understanding.

All strategies generate a binary mask *Mask* ∈ ℝ*^N^*^×^*^C^* for each input image, with configurable masking ratios. Instead of extracting and concatenating visible tokens [12], we replaced masked contents with a specific token and further zeroed the attention weights in the channel-level encoder to explicitly make the masked contents invisible.

#### Training

Eva was implemented using PyTorch v2.6.0. The Vision Transformer (ViT) components were implemented based on PyTorch Image Models (Timm). Eva was trained solely with random masking, with a mask ratio of 0.75. In each training batch, we always included the DAPI channel and H&E channels. We also set a fixed maximum number of channels for each batch. For patches with more channels, we randomly sub-sampled the available channels. For patches with fewer channels, we zero-padded the channel dimension and used a channel mask to prevent the padded channels from participating in attention or loss computation. In particular, we set the maximum number of channels to 22 in training. The spatial dimension of input patch images in training was 224×224, and the spatial dimension of tokens was 8×8.

We used a fixed learning rate of 10^-4^, followed by a cosine annealing to 10^-5^. The whole training process lasted 20 epochs with a batch size of 16. The training framework and monitoring were implemented using PyTorch Lightning and Wandb. The backbone of the token-level encoder is a ViT-base model. The pretraining of Eva was conducted on 8×141 GB NVIDIA H200 GPUs, while all downstream experiments were carried out on a single 24 GB NVIDIA RTX 4090 GPU. Other detailed parameters can be found in **Supplementary Table 14**.

#### H&E to MIF Fine-tuning

To evaluate the model capability on the H&E to MIF prediction task, we fine-tuned Eva using an MIF masking objective, i.e., predicting MIF solely from H&E input. Specifically, the inputs consist of the H&E image and the marker names, and the model is tasked with predicting the intensity of the marker channel based on the marker name. Fine-tuning was performed on the same dataset used for Eva’s pretraining for 5 epochs with a fixed learning rate of 1×10⁻⁵. The fine-tuned model was used to assess prediction performance and to visualize the H&E to MIF prediction results.

#### Linear probing for downstream tasks

Linear probing is a widely used downstream evaluation protocol for foundation models [8,9,63], providing effective means to assess the expressiveness of embeddings. Specifically, a lightweight supervised classifier is trained on top of the foundation model embeddings to predict specific downstream task labels, during which weights of the foundation model backbone are fixed.

In our implementation, we adopted a unified linear probing setup across all backbone models to ensure fair comparison. We trained a single-layer projection that maps the feature space from ℝ*^D^*^model^ to ℝ*^D^*^class^ for classification tasks, or ℝ for regression tasks, without introducing any non-linear activation functions. The linear classifier was optimized using the AdamW optimizer [64] with a learning rate of 10^-4^ for 20 epochs, while keeping all backbone parameters fixed.

### Downstream tasks

Eva was evaluated on a wide range of downstream tasks, which can be broadly categorized into two classes: cell/tissue-level tasks, such as cell type prediction, cell composition prediction, and microenvironment classification; and clinical-level tasks, including tumor type classification, patient stratification, and survival analysis. For each task, we employed different evaluation strategies and metrics to assess the performance of different models.

#### Quality control of images

Image quality metrics including NIQE scores, biomarker grades, and imaging artifact presence, were annotated as described above. We used Eva embeddings of patch images to predict region quality labels via linear probing with a frozen backbone and multiple instance learning.

#### Cell type prediction

Cell type labels were annotated as described above. To predict cell types, we extracted 32×32 blocks centered at the centroids of segmented cells. The spatial dimension of 32 is picked to empirically match the approximate size of a single cell (10-30 μm). We placed the 32×32 single-cell blocks in the middle of 224×224 empty patches and further employed two masking strategies: cell masking and box masking. In cell masking, only pixels within the target cell boundary from cell segmentation are retained, with all pixels outside the boundary set to zeros. In box masking, all pixels within the 32×32 blocks are preserved.

We calculated Eva embeddings for the patches containing single-cell blocks and used them to predict cell type labels via linear probing. For each dataset, we sampled up to 1,000 cells from each cell type to maintain data balance. The datasets used for cell type prediction are described above. On average, each dataset contains 15 cell types (15 ± 5 per dataset), with around 14,000 cells (13912 ± 3585 per dataset) included for linear probe training. An 8:1:1 split was applied to construct the training, validation, and test sets.

In the zero-shot and few-shot generalization tests of cell type prediction, we first unified cell type labels between the source and the target datasets into a consistent list (**Supplementary Figure 14**). For zero-shot evaluation, the linear probing model was trained and validated on the source dataset and directly tested on the held-out test set of the target dataset. For few-shot evaluations, we incorporated varying amounts of cells from the target dataset for training and evaluated performance on the held-out test set of the target dataset.

#### Cell composition prediction

Cell type composition labels for 224×224 image patches were calculated by counting cells in the patches and normalizing the cell type counts, resulting in cell composition vectors 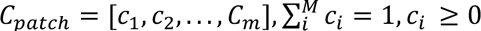, where *M* is the number of cell types, and *c_i_* is the proportion of the *i*-th cell type in the patch. We calculated Eva embeddings for patches and used them to predict cell composition through regression tasks via linear probing. The datasets used for cell composition prediction are described above. On average, each dataset contains 13,000 patches (13,311 ± 12,008 per dataset). We split the data into training, validation, and test sets using a ratio of 8:1:1.

#### Microenvironment classification

Microenvironment annotations (e.g., lymphocyte-rich tumor, myeloid cell-rich tumor, tumor-immune interface, tertiary lymphoid structure) were derived from previous analysis of UPMC-HNC [36] and MDACC-HCC [56]. Each cell was assigned a microenvironment label according to its local neighborhood (50∼100 μm), calculated from graph neural network embeddings.

We calculated Eva embeddings for 224×224 patches centered at cells with known microenvironment annotations. The embeddings were then used to predict the corresponding microenvironment class labels via linear probing.

#### Zero-shot image retrieval

We evaluated Eva’s embedding quality through zero-shot image retrieval. Embeddings of multiplexed image patches were generated by Eva. For a randomly selected query image patch, cosine similarity was computed between its embedding and support embeddings from the reference database:

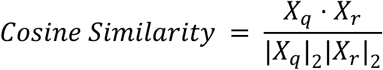

where *X*_q_, *X*_r_ ∈ ℝ*^Dmodel^* are the embeddings of query patch and reference patch, respectively. We excluded patches from the same region as the query patch to avoid potential bias and overfitting. For the retrieved patches, we evaluated whether they contained the same dominant cell type, derived from cell composition annotation, as the query patch.

#### Tumor type classification

Tumor type labels were acquired from metadata of the EM-PCA dataset, comprising nine major tumor types and additional subtypes for lung cancer (e.g., adenocarcinoma, squamous cell carcinoma, and large cell carcinoma). Each region was assigned a single tumor type and, when applicable, subtype label.

We calculated Eva embeddings for held-out regions from the EM-PCA dataset and used them to predict the region-level tumor type (and subtype) classes via linear probing. This task was evaluated through a random 5-fold cross-validation.

#### Patient stratification

Case-level labels of tumor characteristics (e.g., HPV positivity in UPMC-HNC), precision treatment (e.g., response to immunotherapy in MDACC-HCC), and prognosis (e.g., primary outcome of no-evidence-of-disease versus died-of-disease in Stanford-CRC) were acquired from respective validation datasets. Binary class labels for each task were curated and assigned to the relevant regions.

For each task, we calculated Eva embeddings for regions from the corresponding dataset and used them to predict the region-level patient stratification labels via linear probing. This task is evaluated through a random 5-fold cross-validation.

#### Survival analysis

Survival analysis is primarily used in clinical research to evaluate treatment effectiveness and identify key factors influencing prognosis. Survival outcomes, including event labels and time-to-event data, were acquired from the clinical metadata of two validation datasets: UPMC-HNC and EM-PCA (the colorectal cancer subset).

We calculated Eva embeddings for regions and used them to train a linear model using Cox proportional hazards survival loss function [50,51]:

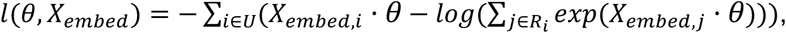

where *U* is the set of uncensored patients, and *R_i_* is the risk set at time *t_i_*, which includes all cases who have not yet encountered the event or been censored at that time point. *θ* is the parameters of the additional linear layer, and *X_embed_*_,*i*_ is the region-level Eva embedding for the *i*-th case. The inner product *X_embed_ θ* represents the predicted risk score for the case. This task is evaluated through a random 5-fold cross-validation.

#### Zero-shot case retrieval

Similarly, we evaluated Eva’s region-level embeddings through zero-shot case retrieval. For each query case/region, cosine similarity was computed between its embedding and support embeddings of other cases in the dataset. For the retrieved results, we evaluated whether they shared the same labels.

### Metrics

We applied various metrics for the reconstruction evaluations and downstream tasks introduced above:

- For reconstruction evaluation, we used mean squared error (MSE), Spearman’s rank correlations and Pearson correlation coefficient (PCC) to quantify the patch reconstruction quality of Eva. For random and token masking, the mask ratio was set to 0.75. The metrics were evaluated on masked tokens and channels.
- For cross-modality prediction, we used Spearman’s rank correlation, PCC, and structural similarity index measure (SSIM), calculated at both pixel and grid levels (8×8 average pooling). To ensure a fair comparison across predictions with varying output scales, all metrics were computed after applying quantile matching to align the prediction distributions with the ground truth.
- For multi-class classification tasks, including cell type, microenvironment, and tumor type classification, we calculated one-vs-rest multi-class receiver operating characteristic (ROC) for each class and reported the average area under the curve (AUC). We also evaluated and reported per-class metrics using F1-score for tumor type and cell type prediction tasks.
- For binary classification tasks, including quality control and patient stratification tasks, we used the area under the receiver operating characteristic curve (AUC) as the evaluation metric.
- For the cell composition prediction task, we employed PCC and MSE as evaluation metrics, which assess the relative and absolute discrepancy between predicted and ground truth cell compositions.
- For the patch-level zero-shot image retrieval task, we used Top-K accuracy and MSE as evaluation metrics. Top-K accuracy assesses whether the correct result is included among the top k retrieved results, where the dominant cell type of a patch is used as the label. Top-K MSE is computed as the lowest MSE between cell compositions of the retrieved patches and the query patch. We reported Top-k accuracy for k=1, 3, 5, and MSE for the patch-level zero-shot image retrieval task.
- For the region-level zero-shot case retrieval task, we similarly employed Top-1 accuracy as the evaluation metric, where the patient stratification task label for the region is used.
- For survival analysis tasks, we employed standard evaluation metrics, including the concordance index (C-index) and comparison of Kaplan-Meier (KM) curves with p-values calculated from log-rank tests. C-index quantifies the discriminative ability of a survival model by measuring the proportion of comparable pairs in which the predicted risk rankings agree with the observed survival times. KM curves were drawn for high-risk and low-risk groups, defined by bi-secting test cases into two equal-sized groups based on their predicted risk scores. Log-rank tests were subsequently performed to compare survival distributions between the high-risk and low-risk groups, providing p-values to assess whether statistically significant differences exist [47].

All metrics were implemented using Python packages scikit-learn, PyTorch, and lifelines.

### Related models

To contextualize the performance of Eva, we utilized related models for embedding extraction and/or benchmarking, including the latest multiplexed imaging foundation models VirTues [19] and KRONOS [16], dedicated models for H&E to MIF translation including ROSIE [22] and GigaTIME [23], as well as pathology foundation models (PFMs) UNI [8] and Prov-GigaPath [52].

#### VirTues

VirTues [19] is a multiplexed imaging foundation model trained on IMC data. It employs a masked autoencoder (MAE) based model for both biomarker and token-level reconstruction. We employed the open-sourced weights of VirTues, and extracted summary tokens *X_virTues_* ∈ ℝ^512^ of MIF image patches, which is the averaged embeddings of all input image tokens. Biomarkers were represented as protein sequences retrieved from UniProt [66] and encoded using ESM [1]. For markers trained by VirTues, we retained their original protein sequences to maintain consistency and standardized the nomenclature for identical markers named differently in VirTues and Eva. For markers not trained by VirTues, we obtained their protein sequences, extracted their ESM embeddings, and integrated them into the model. All data preprocessing, normalization, evaluation and downstream embedding usage strategies of VirTues followed the original manuscript and the official implementation. The released weights of VirTues can be found at https://huggingface.co/bunnelab/virtues.

#### KRONOS

KRONOS [16] is a multiplexed imaging foundation model trained using the DINOv2 [11] framework. To evaluate the model’s performance, we utilized the open-source model weights of KRONOS. The biomarker embedding of KRONOS was trained using a fixed positional embedding, which is sensitive to the input order and type of biomarkers. Consequently, during evaluation, only biomarkers pre-trained by KRONOS were passed to it. To avoid omissions caused by inconsistencies in capitalization or differences in nomenclature, we manually aligned the names of biomarkers in the two models. We extracted the classification (CLS) token embeddings *X_KRONOS_* ∈ ℝ^384^ of MIF image patches for downstream tasks, as specified by KRONOS’s manuscript. All data preprocessing, normalization, and evaluation strategies followed the original manuscript and the official implementation.The released weights of KRONOS can be found at https://huggingface.co/MahmoodLab/KRONOS.

#### ROSIE

ROSIE [22] is based on ConvNext [68] and used to predict MIF virtual staining from H&E. The model was trained with 50 markers. We followed the official ROSIE evaluation protocol as released in their implementation. Specifically, the H&E images are processed with a sliding-window inference scheme, where each window is extracted with the official patch size 128×128, resized to 224×224 with ImageNet normalization, and the resulting marker-wise predictions are fused back onto the same size of input image using the Gaussian weighting strategy. As described in ROSIE, the raw output values without postprocessing were used for quantitative analysis or stitching into large regions after quantile matching. The released weights of ROSIE can be found at https://huggingface.co/ericwu09/ROSIE.

#### GigaTIME

GigaTIME [23], built upon the UNet++ architecture [67], is an AI model designed to translate H&E images into virtual multiplex MIF images, trained with 21 markers. Its original output is optimized for binary prediction; we adapted the post-sigmoid outputs to evaluate the model’s pixel-level performance. For patch-level predictions, the H&E patches were resized and processed by GigaTIME’s inference pipeline and stitched back when computing metrics or visualization. Specifically, evaluation metrics were computed following quantile matching, which aligned the predicted intensity rankings with the ground truth. To ensure compatibility with Eva and accurately assess marker performance, we standardized the nomenclature by mapping the GigaTIME markers CD3, Actin-D, and CK to CD3e, aSMA, and PanCK, respectively. The released weights of GigaTIME can be found at https://huggingface.co/prov-gigatime/GigaTIME.

#### UNI

UNI [8] is an H&E foundation model trained on 200 million histopathology images (350,000 slides) based on the DINOv2 framework. To establish a reference for case-level predictions using H&E inputs, we used the open-source model weights of UNI to extract the classification (CLS) token embeddings *X_UNI_* ∈ ℝ^1024^ of H&E image patches for applicable case-level tasks, as specified by UNI’s implementation. The released weights of UNI can be found at https://huggingface.co/MahmoodLab/UNI.

#### Prov-GigaPath

Prov-GigaPath [52] is a whole-slide PFM trained on 1.3 billion histopathology image tiles (171,189 whole slides) based on the DINOv2 framework for its tile encoder. To establish a reference for case-level predictions using H&E inputs, we used the open-source model weights of Prov-GigaPath to extract the classification (CLS) token embeddings *X_prov-Gigapath_* ∈ ℝ^1536^ of H&E image patches for applicable case-level downstream tasks, as specified by Prov-GigaPath’s implementation. The released weights of Prov-GigaPath can be found at https://huggingface.co/prov-gigapath/prov-gigapath.

## Data and code availability

The code and model weights of Eva are available at: https://github.com/YAndrewL/Eva and https://huggingface.co/yandrewl/Eva, respectively.

Public data used in this study can be accessed at:

- UPMC-HNC, Stanford-CRC, DFCI-HNC: https://zenodo.org/records/13179600 and https://app.enablemedicine.com/portal/atlas-library/studies/92394a9f-6b48-4897-87de-999614952d94?sid=1168
- MDACC-HCC: https://zenodo.org/records/15392699
- RAHBT-DCIS: https://data.mendeley.com/datasets/d87vg86zd8
- IUCPQ-LUAD: https://zenodo.org/records/7383627
- Stanford-HCC: https://springernature.figshare.com/articles/dataset/Spatial_Analysis_Reveals_Target able_Macrophage-Mediated_Mechanisms_of_Immune_Evasion_in_Hepatocellular_Carcinoma_Minimal_Residual_Disease/26539345
- UHB-CRC: https://data.mendeley.com/datasets/mpjzbtfgfr/1 and https://doi.org/10.7937/tcia.2020.fqn0-0326

## Acknowledgement

This work was supported by the University Research Committee of the University of Hong Kong, and the Research Grants Council of Hong Kong, China under the Early Career Scheme (27202625). We gratefully acknowledge Tianqi AI laboratory for providing the GPU resources that were essential for the computational aspects of this research.

We extend our sincere gratitude to the developers of the VirTues, KRONOS, GigaTIME, and ROSIE models. Their commitment to the development and public release of their models have profoundly benefited the research community. These open-source models established the rigorous comparative standards that were instrumental in guiding the evaluation and development of this work.

We extend our sincere gratitude to the software engineers, scientists, technicians, and support staff at Enable Medicine who collectively built the data platform that made this work possible.

## Author Contributions

Y.L. and R.S. developed the computational models and performed data analysis. M.B., A.K., E.W., P.C., I.L., M.R., E.B., H.B.D., M.M., and A.T.M. collected, prepared, processed, and annotated the data used in the study. R.J., W.D., and L.Q. provided guidance and assistance with model training and evaluation. R.L., G.C., P.S., R.D., C.M.S., and J.Z. facilitated data generation and offered advice for the project. A.E.T. and Z.W. conceptualized the study, designed the methodology, and supervised the project. Y.L., A.E.T., and Z.W. wrote the manuscript. All authors reviewed and edited the manuscript and approved its final version.

## References

1. Lin Z, Akin H, Rao R, Hie B, Zhu Z, Lu W, et al. Evolutionary-scale prediction of atomic-level protein structure with a language model. Science. 2023;379: 1123–1130. doi:10.1126/science.ade2574

2. Madani A, Krause B, Greene ER, Subramanian S, Mohr BP, Holton JM, et al. Large language models generate functional protein sequences across diverse families. Nat Biotechnol. 2023;41: 1099–1106. doi:10.1038/s41587-022-01618-2

3. Nguyen E, Poli M, Durrant MG, Kang B, Katrekar D, Li DB, et al. Sequence modeling and design from molecular to genome scale with Evo. Science. 2024;386: eado9336. doi:10.1126/science.ado9336

4. Avsec Ž, Latysheva N, Cheng J, Novati G, Taylor KR, Ward T, et al. Advancing regulatory variant effect prediction with AlphaGenome. Nature. 2026;649: 1206–1218. doi:10.1038/s41586-025-10014-0

5. Chen J, Hu Z, Sun S, Tan Q, Wang Y, Yu Q, et al. Interpretable RNA foundation model from unannotated data for highly accurate RNA structure and function predictions. arXiv [q-bio.QM]. 2022. doi:10.1101/2022.08.06.503062

6. Tejada-Lapuerta A, Schaar AC, Gutgesell R, Palla G, Halle L, Minaeva M, et al. Nicheformer: a foundation model for single-cell and spatial omics. Nat Methods. 2025; 1–14. doi:10.1038/s41592-025-02814-z

7. Filiot A, Ghermi R, Olivier A, Jacob P, Fidon L, Camara A, et al. Scaling self-Supervised Learning for histopathology with Masked Image Modeling. medRxiv. 2023. p. 2023.07.21.23292757. doi:10.1101/2023.07.21.23292757

8. Chen RJ, Ding T, Lu MY, Williamson DFK, Jaume G, Song AH, et al. Towards a general-purpose foundation model for computational pathology. Nat Med. 2024;30: 850–862. doi:10.1038/s41591-024-02857-3

9. Xiang J, Wang X, Zhang X, Xi Y, Eweje F, Chen Y, et al. A vision-language foundation model for precision oncology. Nature. 2025;638: 769–778. doi:10.1038/s41586-024-08378-w

10. Ma C, Tan W, He R, Yan B. Pretraining a foundation model for generalizable fluorescence microscopy-based image restoration. Nat Methods. 2024;21: 1558–1567. doi:10.1038/s41592-024-02244-3

11. Oquab M, Darcet T, Moutakanni T, Vo H, Szafraniec M, Khalidov V, et al. DINOv2: Learning robust visual features without supervision. arXiv [cs.CV]. 2023. Available: http://arxiv.org/abs/2304.07193

12. He K, Chen X, Xie S, Li Y, Dollár P, Girshick R. Masked Autoencoders Are Scalable Vision Learners. arXiv [cs.CV]. 2021. Available: http://arxiv.org/abs/2111.06377

13. Dosovitskiy A, Beyer L, Kolesnikov A, Weissenborn D, Zhai X, Unterthiner T, et al. An image is worth 16×16 words: Transformers for image recognition at scale. arXiv [cs.CV]. 2020. Available: http://arxiv.org/abs/2010.11929

14. Filiot A, Jacob P, Mac Kain A, Saillard C. Phikon-v2, A large and public feature extractor for biomarker prediction. arXiv [eess.IV]. 2024. Available: http://arxiv.org/abs/2409.09173

15. Lu MY, Chen B, Williamson DFK, Chen RJ, Liang I, Ding T, et al. Towards a visual-language foundation model for computational pathology. arXiv [cs.CV]. 2023. Available: http://arxiv.org/abs/2307.12914

16. Shaban M, Chang Y, Qiu H, Yeo YY, Song AH, Jaume G, et al. A Foundation Model for Spatial Proteomics. arXiv [cs.CV]. 2025. Available: http://arxiv.org/abs/2506.03373

17. Bao Y, Sivanandan S, Karaletsos T. Channel Vision Transformers: An Image Is Worth 1 x 16 x 16 Words. arXiv [cs.CV]. 2023. Available: http://arxiv.org/abs/2309.16108

18. Pham C, Caicedo JC, Plummer BA. ChA-MAEViT: Unifying channel-aware Masked Autoencoders and Multi-channel vision transformers for improved cross-channel learning. arXiv [cs.CV]. 2025. Available: http://arxiv.org/abs/2503.19331

19. Wenckstern J, Jain E, Vasilev K, Pariset M, Wicki A, Gut G, et al. AI-powered virtual tissues from spatial proteomics for clinical diagnostics and biomedical discovery. arXiv [q-bio.QM]. 2025. doi:10.48550/arXiv.2501.06039

20. Vaswani A, Shazeer N, Parmar N, Uszkoreit J, Jones L, Gomez AN, et al. Attention is all you need. arXiv [cs.CL]. 2017. Available: http://arxiv.org/abs/1706.03762

21. Chen Y, Zou J. Simple and effective embedding model for single-cell biology built from ChatGPT. Nat Biomed Eng. 2025;9: 483–493. doi:10.1038/s41551-024-01284-6

22. Wu E, Bieniosek M, Wu Z, Thakkar N, Charville GW, Makky A, et al. ROSIE: AI generation of multiplex immunofluorescence staining from histopathology images. bioRxivorg. 2024. doi:10.1101/2024.11.10.622859

23. Valanarasu JMJ, Xu H, Usuyama N, Kim C, Wong C, Argaw P, et al. Multimodal AI generates virtual population for tumor microenvironment modeling. Cell. 2026;189: 386–400.e19. doi:10.1016/j.cell.2025.11.016

24. Li Z, Li Y, Xiang J, Wang X, Yang S, Zhang X, et al. AI-enabled virtual spatial proteomics from histopathology for interpretable biomarker discovery in lung cancer. Nat Med. 2026;32: 231–244. doi:10.1038/s41591-025-04060-4

25. Wang Z, Bovik AC, Sheikh HR, Simoncelli EP. Image quality assessment: from error visibility to structural similarity. IEEE Trans Image Process. 2004;13: 600–612. doi:10.1109/tip.2003.819861

26. Rumberger JL, Greenwald NF, Ranek JS, Boonrat P, Walker C, Franzen J, et al. Automated classification of cellular expression in multiplexed imaging data with Nimbus. Nat Methods. 2025;22: 2161–2170. doi:10.1038/s41592-025-02826-9

27. Liu CC, Greenwald NF, Kong A, McCaffrey EF, Leow KX, Mrdjen D, et al. Robust phenotyping of highly multiplexed tissue imaging data using pixel-level clustering. Nat Commun. 2023;14: 4618. doi:10.1038/s41467-023-40068-5

28. Zhang W, Li I, Reticker-Flynn NE, Good Z, Chang S, Samusik N, et al. Identification of cell types in multiplexed in situ images by combining protein expression and spatial information using CELESTA. Nat Methods. 2022;19: 759–769. doi:10.1038/s41592-022-01498-z

29. Amitay Y, Bussi Y, Feinstein B, Bagon S, Milo I, Keren L. CellSighter: a neural network to classify cells in highly multiplexed images. Nat Commun. 2023;14: 4302. doi:10.1038/s41467-023-40066-7

30. Gerlinger M, Rowan AJ, Horswell S, Math M, Larkin J, Endesfelder D, et al. Intratumor heterogeneity and branched evolution revealed by multiregion sequencing. N Engl J Med. 2012;366: 883–892. doi:10.1056/NEJMoa1113205

31. Binnewies M, Roberts EW, Kersten K, Chan V, Fearon DF, Merad M, et al. Understanding the tumor immune microenvironment (TIME) for effective therapy. Nat Med. 2018;24: 541–550. doi:10.1038/s41591-018-0014-x

32. Lu MY, Williamson DFK, Chen TY, Chen RJ, Barbieri M, Mahmood F. Data-efficient and weakly supervised computational pathology on whole-slide images. Nat Biomed Eng. 2021;5: 555–570. doi:10.1038/s41551-020-00682-w

33. Campanella G, Hanna MG, Geneslaw L, Miraflor A, Werneck Krauss Silva V, Busam KJ, et al. Clinical-grade computational pathology using weakly supervised deep learning on whole slide images. Nat Med. 2019;25: 1301–1309. doi:10.1038/s41591-019-0508-1

34. Kleshchevnikov V, Shmatko A, Dann E, Aivazidis A, King HW, Li T, et al. Cell2location maps fine-grained cell types in spatial transcriptomics. Nat Biotechnol. 2022;40: 661–671. doi:10.1038/s41587-021-01139-4

35. Elosua-Bayes M, Nieto P, Mereu E, Gut I, Heyn H. SPOTlight: seeded NMF regression to deconvolute spatial transcriptomics spots with single-cell transcriptomes. Nucleic Acids Res. 2021;49: e50. doi:10.1093/nar/gkab043

36. Wu Z, Trevino AE, Wu E, Swanson K, Kim HJ, D’Angio HB, et al. Graph deep learning for the characterization of tumour microenvironments from spatial protein profiles in tissue specimens. Nat Biomed Eng. 2022;6: 1435–1448. doi:10.1038/s41551-022-00951-w

37. Schürch CM, Bhate SS, Barlow GL, Phillips DJ, Noti L, Zlobec I, et al. Coordinated cellular neighborhoods orchestrate antitumoral immunity at the colorectal cancer invasive front. Cell. 2020;182: 1341–1359.e19. doi:10.1016/j.cell.2020.07.005

38. Ding T, Wagner SJ, Song AH, Chen RJ, Lu MY, Zhang A, et al. A multimodal whole-slide foundation model for pathology. Nat Med. 2025;31: 3749–3761. doi:10.1038/s41591-025-03982-3

39. Xu K, Qin M, Sun F, Wang Y, Chen Y-K, Ren F. Learning in the frequency domain. arXiv [cs.CV]. 2020. Available: http://arxiv.org/abs/2002.12416

40. Yao J, Zhu X, Jonnagaddala J, Hawkins N, Huang J. Whole slide images based cancer survival prediction using attention guided deep multiple instance learning networks. Med Image Anal. 2020;65: 101789. doi:10.1016/j.media.2020.101789

41. Ilse M, Tomczak JM, Welling M. Attention-based deep multiple instance learning. arXiv [cs.LG]. 2018. Available: http://arxiv.org/abs/1802.04712

42. Coudray N, Ocampo PS, Sakellaropoulos T, Narula N, Snuderl M, Fenyö D, et al. Classification and mutation prediction from non-small cell lung cancer histopathology images using deep learning. Nat Med. 2018;24: 1559–1567. doi:10.1038/s41591-018-0177-5

43. Lu M, Pan Y, Liu F, Shen D. SMILE: Sparse-attention based multiple instance contrastive learning for glioma sub-type classification using pathological images. Atzori M, Burlutskiy N, Ciompi F, Li Z, Minhas F, Müller H, et al., editors. 2021;156: 159–169. Available: https://proceedings.mlr.press/v156/lu21a.html

44. Karasaki T, Moore DA, Veeriah S, Naceur-Lombardelli C, Toncheva A, Magno N, et al. Evolutionary characterization of lung adenocarcinoma morphology in TRACERx. Nat Med. 2023;29: 833–845. doi:10.1038/s41591-023-02230-w

45. Martínez-Jiménez F, Movasati A, Brunner SR, Nguyen L, Priestley P, Cuppen E, et al. Pan-cancer whole-genome comparison of primary and metastatic solid tumours. Nature. 2023;618: 333–341. doi:10.1038/s41586-023-06054-z

46. Cox DR. Regression models and life-tables. J R Stat Soc Series B Stat Methodol. 1972;34: 187–202. doi:10.1111/j.2517-6161.1972.tb00899.x

47. Wang P, Li Y, Reddy CK. Machine learning for survival analysis. ACM Comput Surv. 2019;51: 1–36. doi:10.1145/3214306

48. Kaplan EL, Meier P. Nonparametric Estimation from Incomplete Observations. J Am Stat Assoc. 1958;53: 457. doi:10.2307/2281868

49. Harrell FE Jr, Lee KL, Mark DB. Multivariable prognostic models: issues in developing models, evaluating assumptions and adequacy, and measuring and reducing errors. Stat Med. 1996;15: 361–387. doi:10.1002/(SICI)1097-0258(19960229)15:4<361::AID-SIM168>3.0.CO;2-4

50. Song AH, Chen RJ, Jaume G, Vaidya AJ, Baras AS, Mahmood F. Multimodal Prototyping for cancer survival prediction. arXiv [cs.CV]. 2024. Available: http://arxiv.org/abs/2407.00224

51. Zadeh SG, Schmid M. Bias in cross-entropy-based training of deep survival networks. IEEE Trans Pattern Anal Mach Intell. 2021;43: 3126–3137. doi:10.1109/TPAMI.2020.2979450

52. Xu H, Usuyama N, Bagga J, Zhang S, Rao R, Naumann T, et al. A whole-slide foundation model for digital pathology from real-world data. Nature. 2024;630: 181–188. doi:10.1038/s41586-024-07441-w

53. Heimberg G, Kuo T, DePianto DJ, Salem O, Heigl T, Diamant N, et al. A cell atlas foundation model for scalable search of similar human cells. Nature. 2025;638: 1085–1094. doi:10.1038/s41586-024-08411-y

54. Cancer Genome Atlas Research Network, Weinstein JN, Collisson EA, Mills GB, Shaw KRM, Ozenberger BA, et al. The Cancer Genome Atlas Pan-Cancer analysis project. Nat Genet. 2013;45: 1113–1120. doi:10.1038/ng.2764

55. Janesick A, Shelansky R, Gottscho AD, Wagner F, Williams SR, Rouault M, et al. High resolution mapping of the tumor microenvironment using integrated single-cell, spatial and in situ analysis. Nat Commun. 2023;14: 8353. doi:10.1038/s41467-023-43458-x

56. Wu Z, Boen J, Jindal S, Basu S, Bieniosek M, He S, et al. Spatial multi-omics and deep learning reveal fingerprints of immunotherapy response and resistance in hepatocellular carcinoma. bioRxivorg. 2025. doi:10.1101/2025.06.11.656869

57. Kaseb AO, Hasanov E, Cao HST, Xiao L, Vauthey J-N, Lee SS, et al. Perioperative nivolumab monotherapy versus nivolumab plus ipilimumab in resectable hepatocellular carcinoma: a randomised, open-label, phase 2 trial. Lancet Gastroenterol Hepatol. 2022;7: 208–218. doi:10.1016/S2468-1253(21)00427-1

58. Bentebibel S-E, Johnson D, Pazdrak B, McGrail D, Lecagoonporn S, Haymaker C, et al. 782 Intratumoral sotigalimab with pembrolizumab activates antigen-presenting cells and induces local and distant anti-tumor responses in first-line metastatic melanoma: results of a phase I/II study. Regular and Young Investigator Award Abstracts. BMJ Publishing Group Ltd; 2022. pp. A814–A814. doi:10.1136/jitc-2022-sitc2022.0782

59. Risom T, Glass DR, Averbukh I, Liu CC, Baranski A, Kagel A, et al. Transition to invasive breast cancer is associated with progressive changes in the structure and composition of tumor stroma. Cell. 2022;185: 299–310.e18. doi:10.1016/j.cell.2021.12.023

60. Sorin M, Rezanejad M, Karimi E, Fiset B, Desharnais L, Perus LJM, et al. Single-cell spatial landscapes of the lung tumour immune microenvironment. Nature. 2023;614: 548–554. doi:10.1038/s41586-022-05672-3

61. Lemaitre L, Adeniji N, Suresh A, Reguram R, Zhang J, Park J, et al. Spatial analysis reveals targetable macrophage-mediated mechanisms of immune evasion in hepatocellular carcinoma minimal residual disease. Nat Cancer. 2024;5: 1534–1556. doi:10.1038/s43018-024-00828-8

62. Deng J, Dong W, Socher R, Li L-J, Li K, Fei-Fei L. ImageNet: A large-scale hierarchical image database. 2009 IEEE Conference on Computer Vision and Pattern Recognition. IEEE; 2009. pp. 248–255. doi:10.1109/cvpr.2009.5206848

63. Vorontsov E, Bozkurt A, Casson A, Shaikovski G, Zelechowski M, Severson K, et al. A foundation model for clinical-grade computational pathology and rare cancers detection. Nat Med. 2024;30: 2924–2935. doi:10.1038/s41591-024-03141-0

64. Loshchilov I, Hutter F. Decoupled weight decay regularization. arXiv [cs.LG]. 2017. Available: http://arxiv.org/abs/1711.05101

65. Davidson-Pilon C. lifelines: survival analysis in Python. J Open Source Softw. 2019;4: 1317. doi:10.21105/joss.01317

66. Ahmad S, Jose da Costa Gonzales L, Bowler-Barnett EH, Rice DL, Kim M, Wijerathne S, et al. The UniProt website API: facilitating programmatic access to protein knowledge. Nucleic Acids Res. 2025;53: W547–W553. doi:10.1093/nar/gkaf394

67. Zhou Z, Siddiquee MMR, Tajbakhsh N, Liang J. UNet++: A Nested U-Net Architecture for Medical Image Segmentation. arXiv [cs.CV]. 2018. doi:10.48550/arXiv.1807.10165

68. Liu Z, Mao H, Wu C-Y, Feichtenhofer C, Darrell T, Xie S. A ConvNet for the 2020s. 2022 IEEE/CVF Conference on Computer Vision and Pattern Recognition (CVPR). IEEE; 2022. doi:10.1109/cvpr52688.2022.01167

