## Supplementary Figures and Tables for "Modeling patient tissues at molecular resolution with Eva"

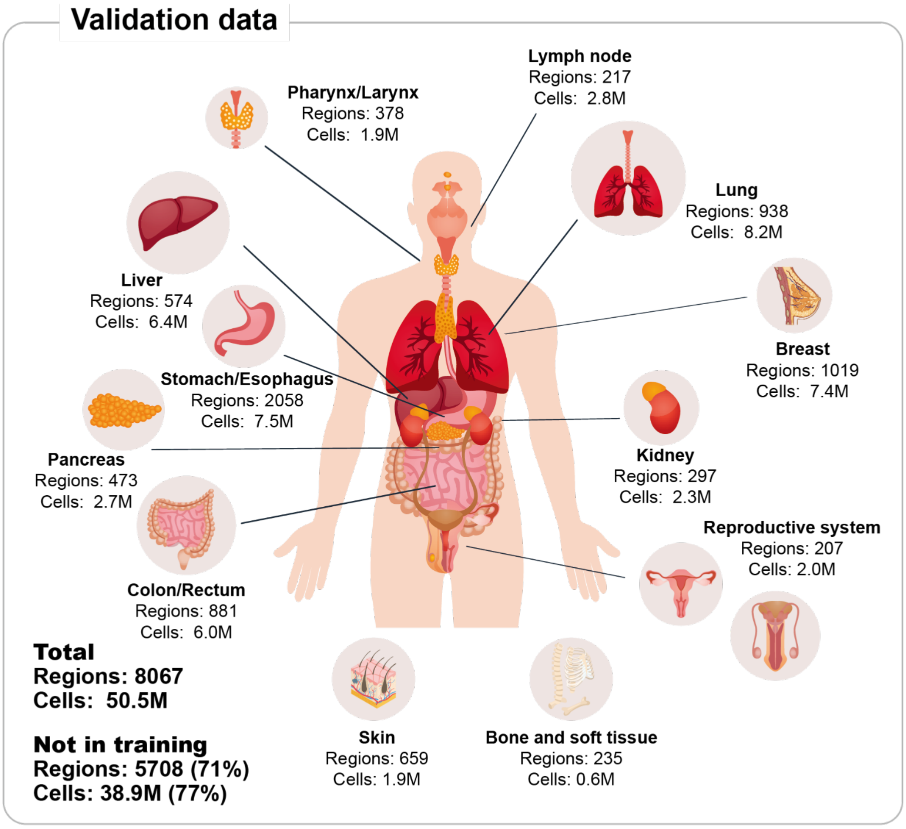


**Supplementary Figure 1: Validation dataset used for evaluating Eva.** 71% of all validation regions were unseen during training, and none of the cell, neighborhood, and patient labels were used during training.

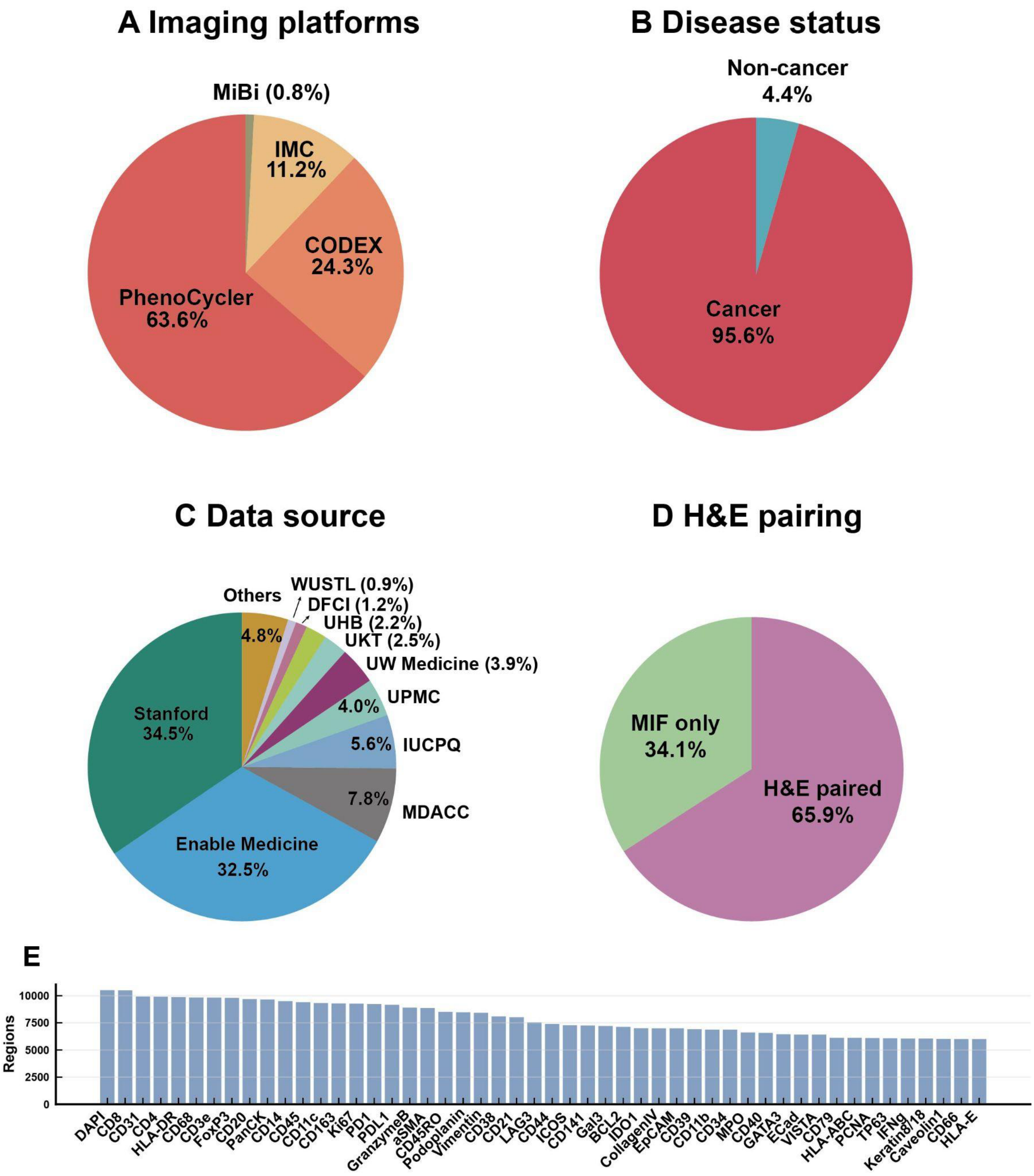


**Supplementary Figure 2. Breakdown of Eva's training and validation data by imaging platforms, data source, and more. A**, Proportion of imaging platforms. **B**, Proportion of disease status, including cancerous and non-cancerous samples. **C**, Proportion of data sources. **D**, Proportion of images with and without paired H&E. **E**, The 50 most frequent immunofluorescence markers, where the y-axis indicates the number of regions in which each marker is used.

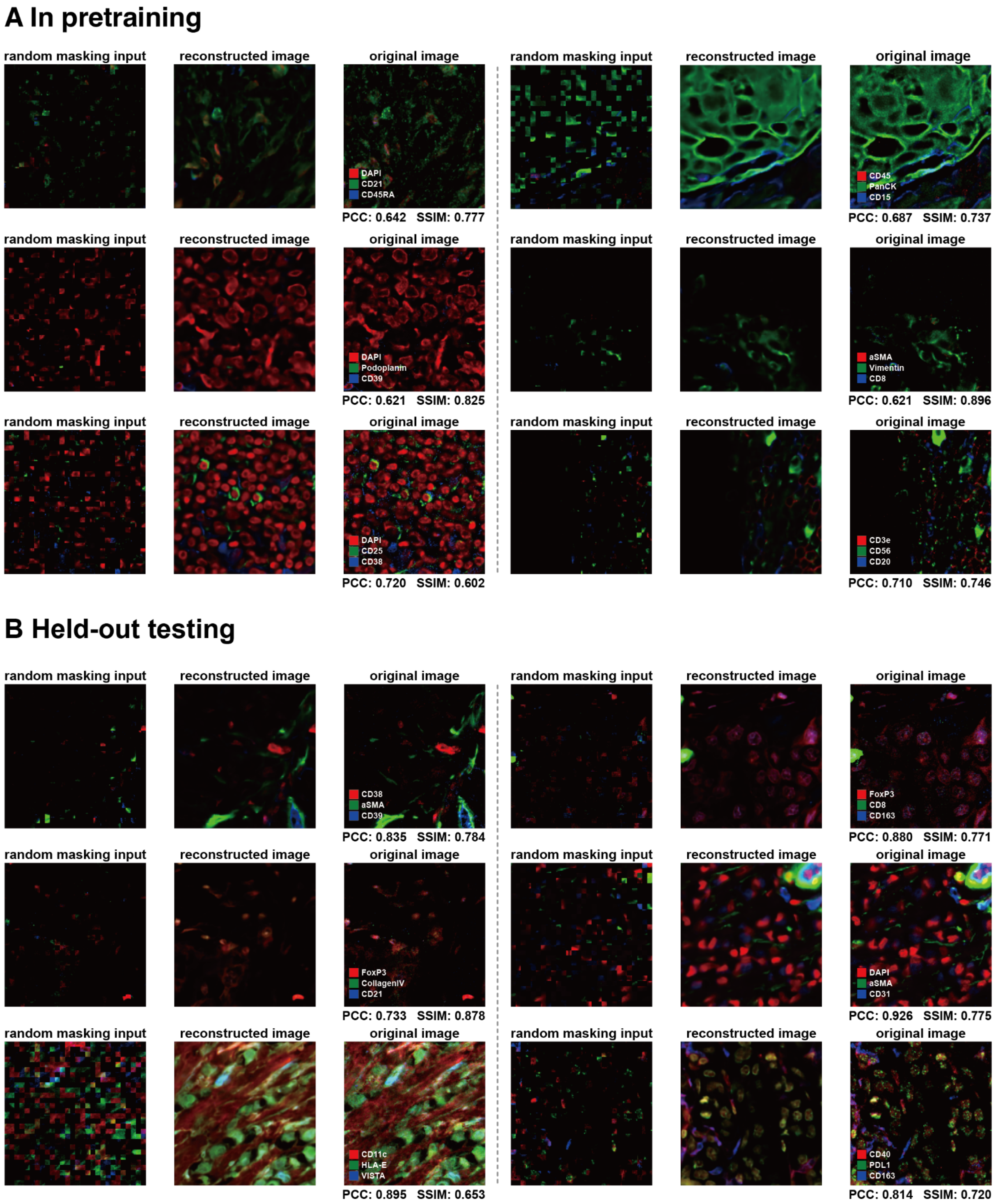


**Supplementary Figure 3. Visualization of reconstruction under random masking (masking ratio = 0.75). A**, Samples from the pretraining dataset. **B**, Samples from the held-out validation dataset. Quantitative evaluation was performed using Pearson correlation coefficient (PCC) and structural similarity index measure (SSIM).

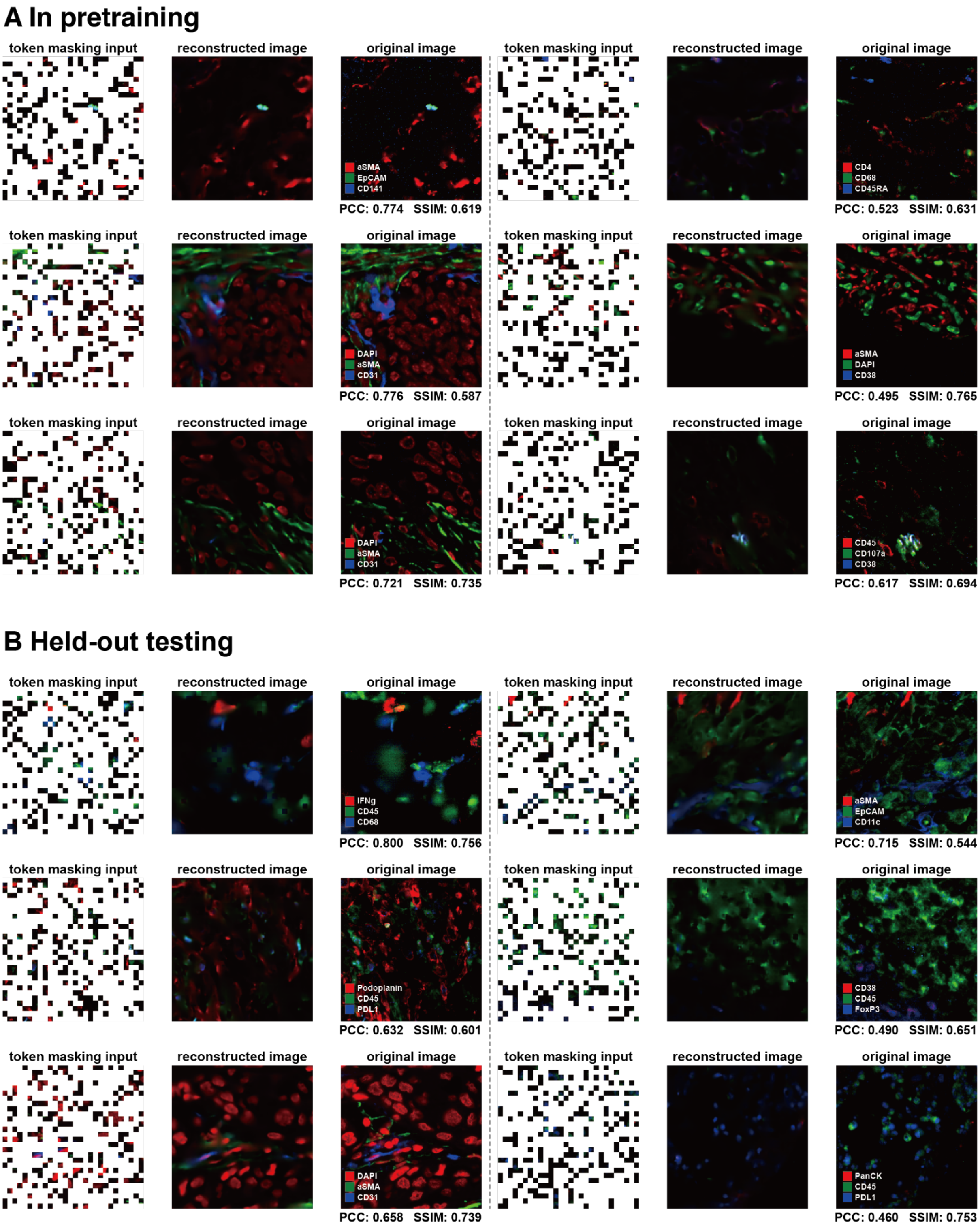


**Supplementary Figure 4. Reconstruction visualization with token masking (masking ratio = 0.75). A**, Example patches from the training dataset. **B**, Example patches from the held-out validation dataset. Quantitative evaluations were performed using PCC and SSIM.

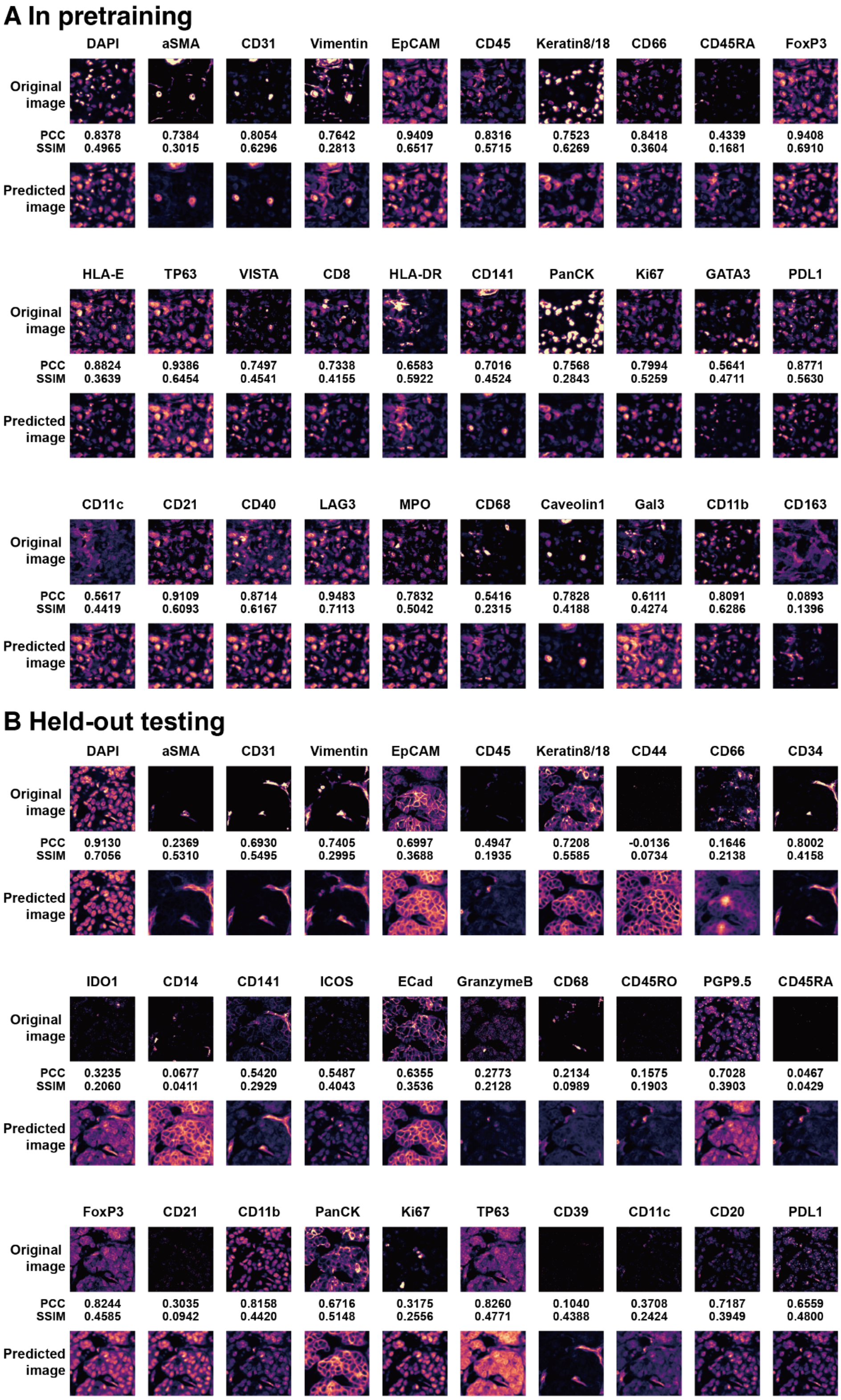


**Supplementary Figure 5. Visualization of reconstruction under channel masking. A**, An example patch from the training dataset. **B**, An example patch from the held-out validation dataset. For each marker, the corresponding channel is masked and then reconstructed by Eva. Quantitative evaluations were performed using PCC and SSIM.

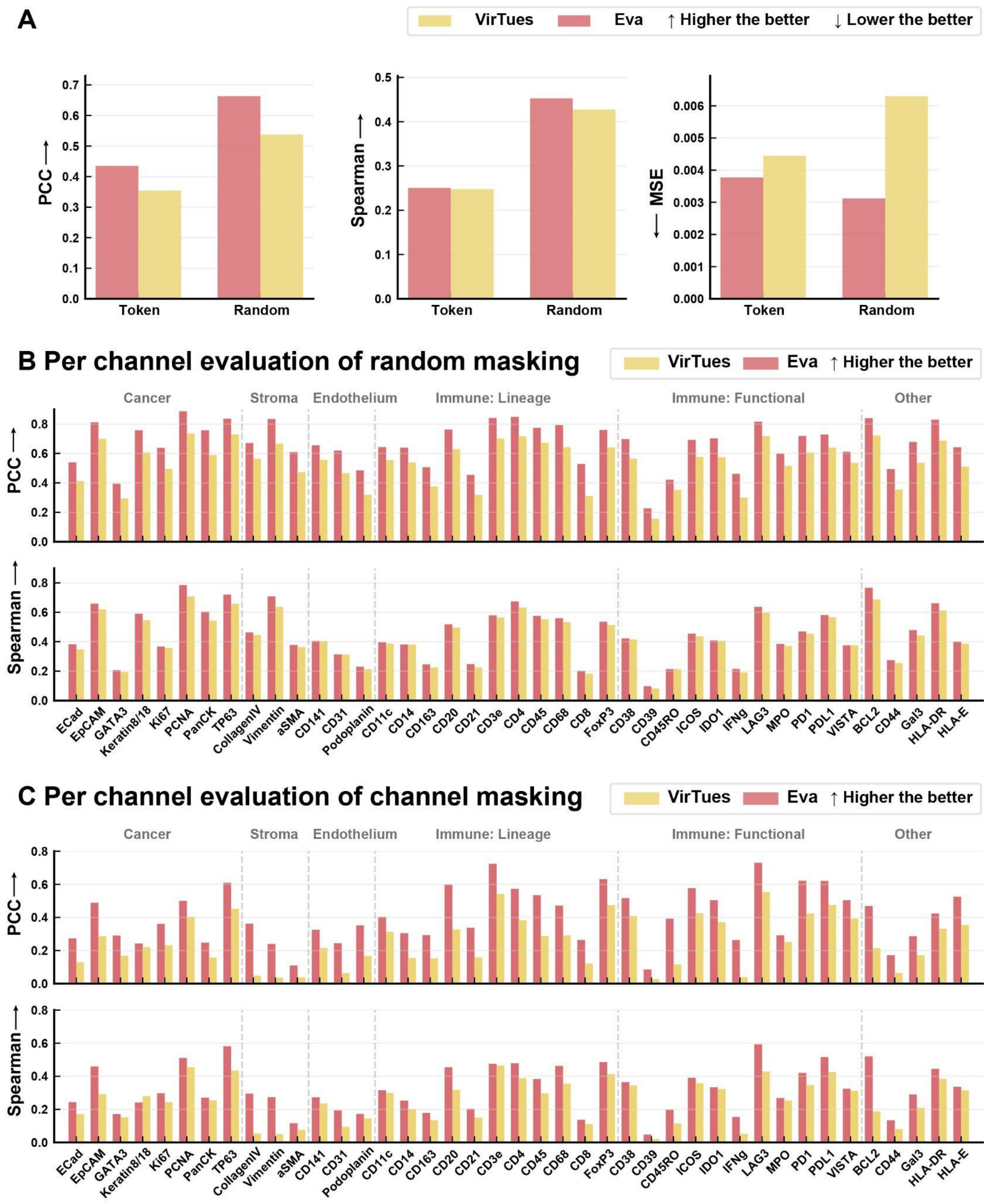


**Supplementary Figure 6. Reconstruction performances of VirTues and Eva on held-out data. A**, Comparison of reconstruction metrics—PCC, SSIM, and mean squared error (MSE)—under token masking and random masking. The masking ratio used for evaluation was 0.75. **B-C**, Comparison of per-channel reconstruction metrics—PCC and SSIM—under random masking and channel masking. The markers are grouped by their primary biological indications in immunofluorescence. A total of 10,000 patches were randomly sampled from the held-out validation dataset to calculate these metrics. For random and token masking, the mask ratio was set to 0.75. Overall metrics were computed on masked tokens and channels.

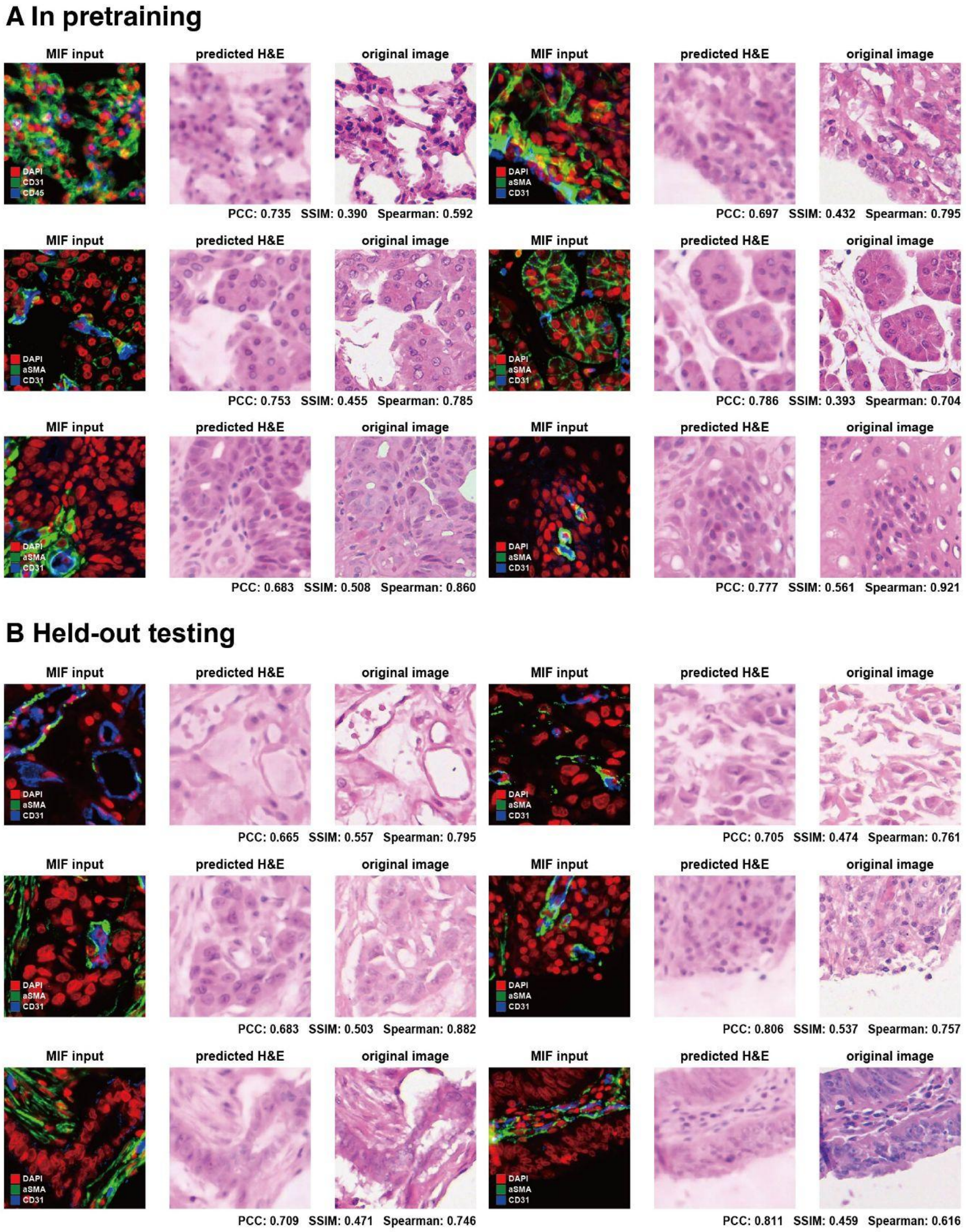


**Supplementary Figure 7. Cross-modality reconstruction from MIF to H&E. A**, Example patches from the training dataset. **B**, Example patches from the held-out validation dataset. Quantitative evaluations were performed using PCC, SSIM, and Spearman's rank correlations.

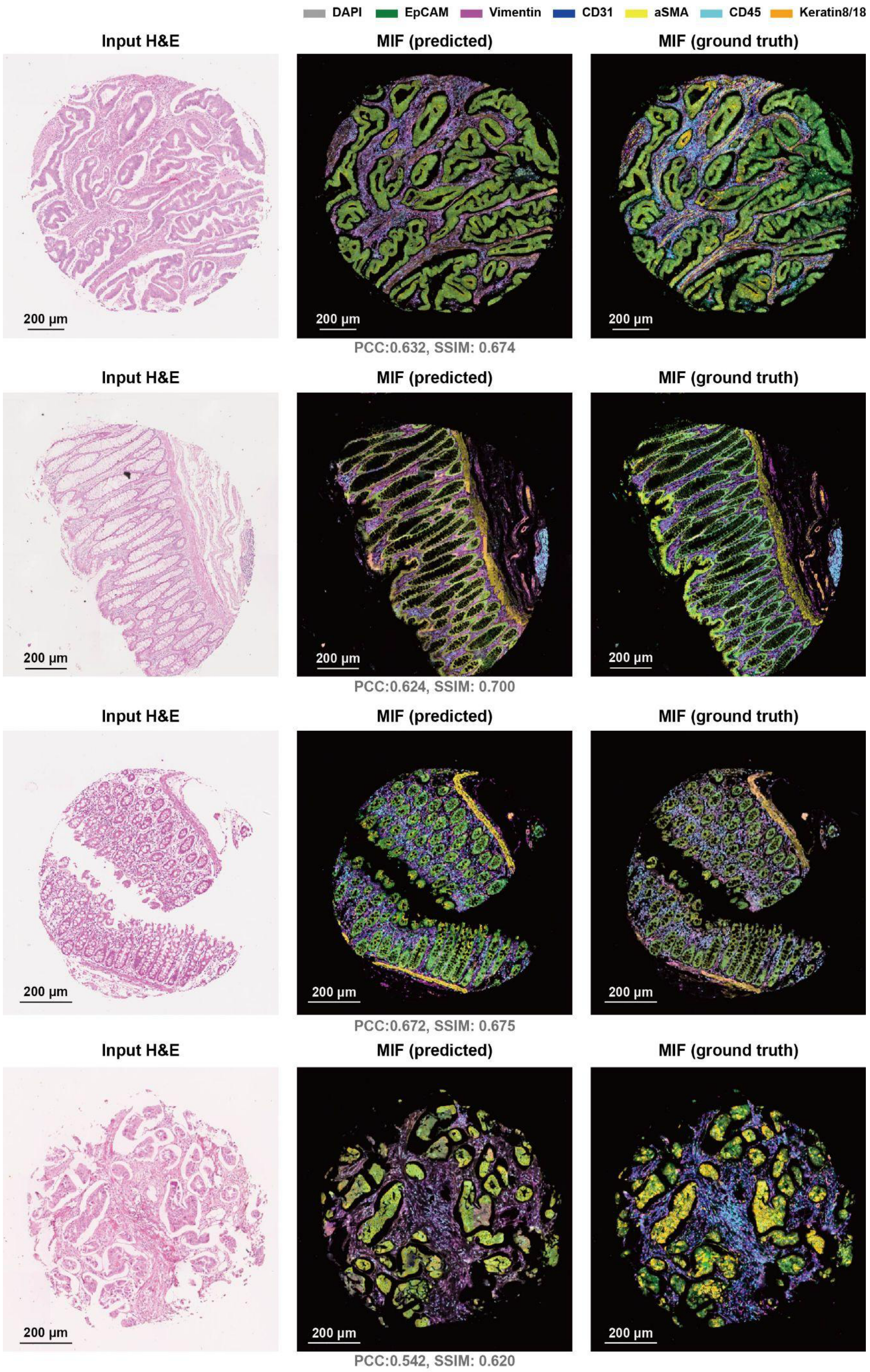


**Supplementary Figure 8. Virtual protein staining using H&E with a fine-tuned Eva model.** Samples were selected from the held-out validation dataset. Quantitative evaluations were performed using PCC and SSIM.

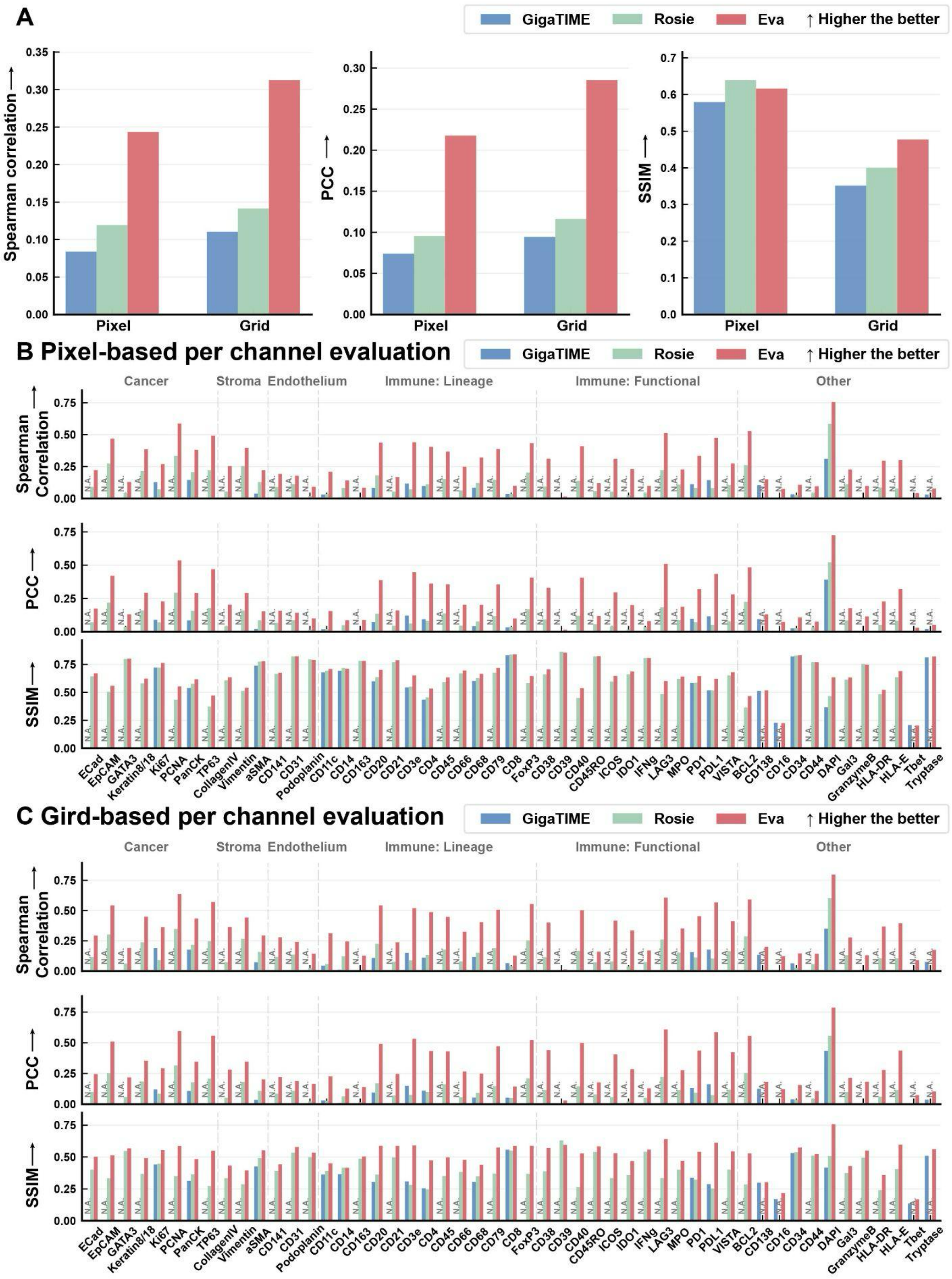


**Supplementary Figure 9. Virtual staining performances of Eva, GigaTIME, and ROSIE on held-out data. A,** Comparison of averaged PCC, Spearman's rank correlations, and SSIM for H&E-based virtual staining by Eva, GigaTIME, and ROSIE. **B-C,** Comparison of per-channel metrics—PCC, Spearman's rank correlations, and SSIM—at pixel- and grid-level (8×8 average pooling). The markers are grouped by their primary biological indications in immunofluorescence. A total of 10,000 patches were randomly sampled from the held-out validation dataset to calculate these metrics. Markers not available from GigaTIME or ROSIE are labeled as N.A.

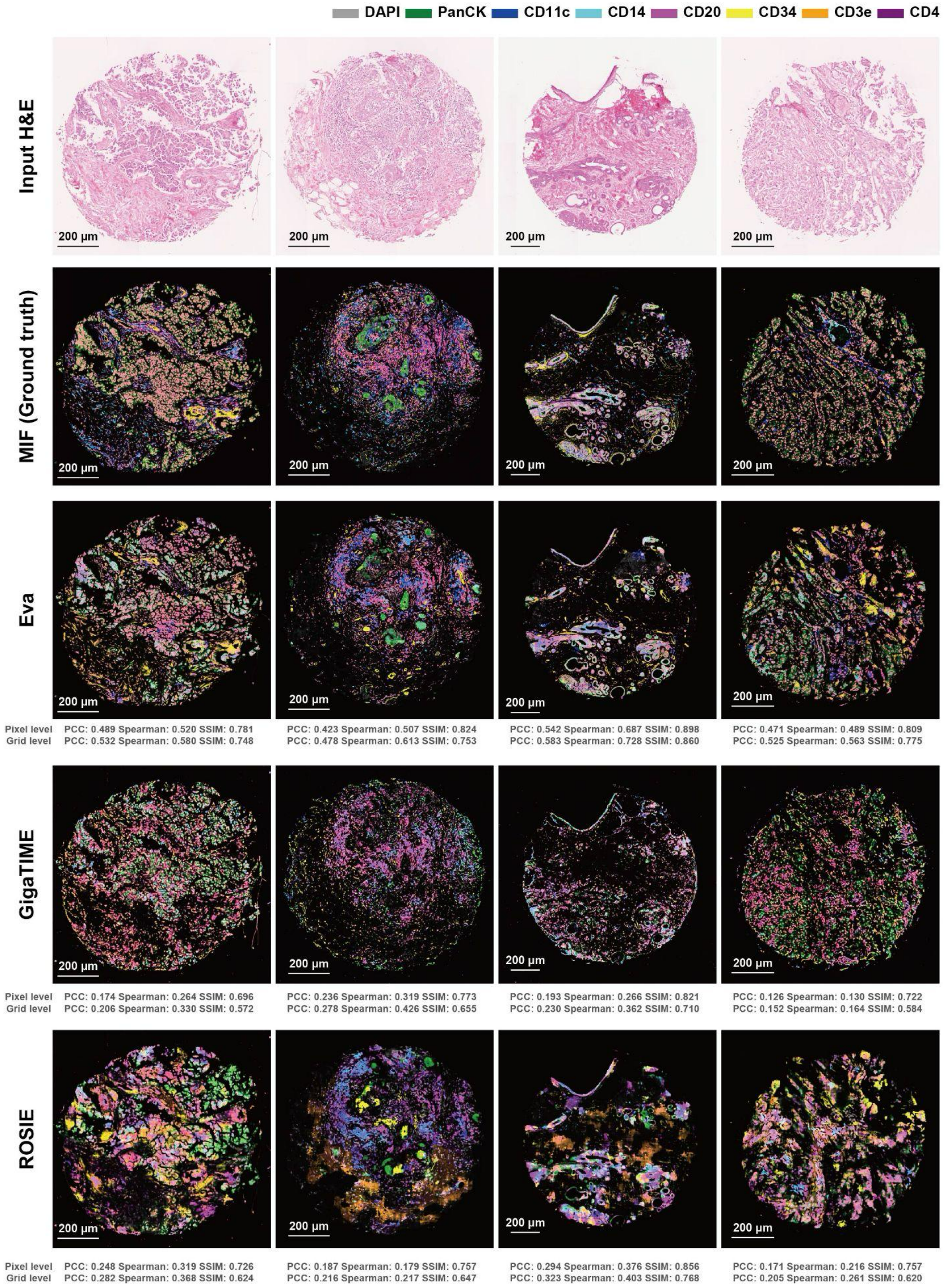


**Supplementary Figure 10. Visualization and comparison of virtual staining performance by Eva, GigaTIME, and ROSIE on the held-out validation dataset.** Virtual staining results on 4 TMA cores. 8 key immunofluorescence markers are shown in different colors. Visualizations display pixel intensities after quantile matching with the ground truth. Computations for metrics including pixel- and grid-level PCC, Spearman's rank correlation, and SSIM are detailed in the **Methods**. The TMA cores were randomly sampled from the held-out validation dataset.

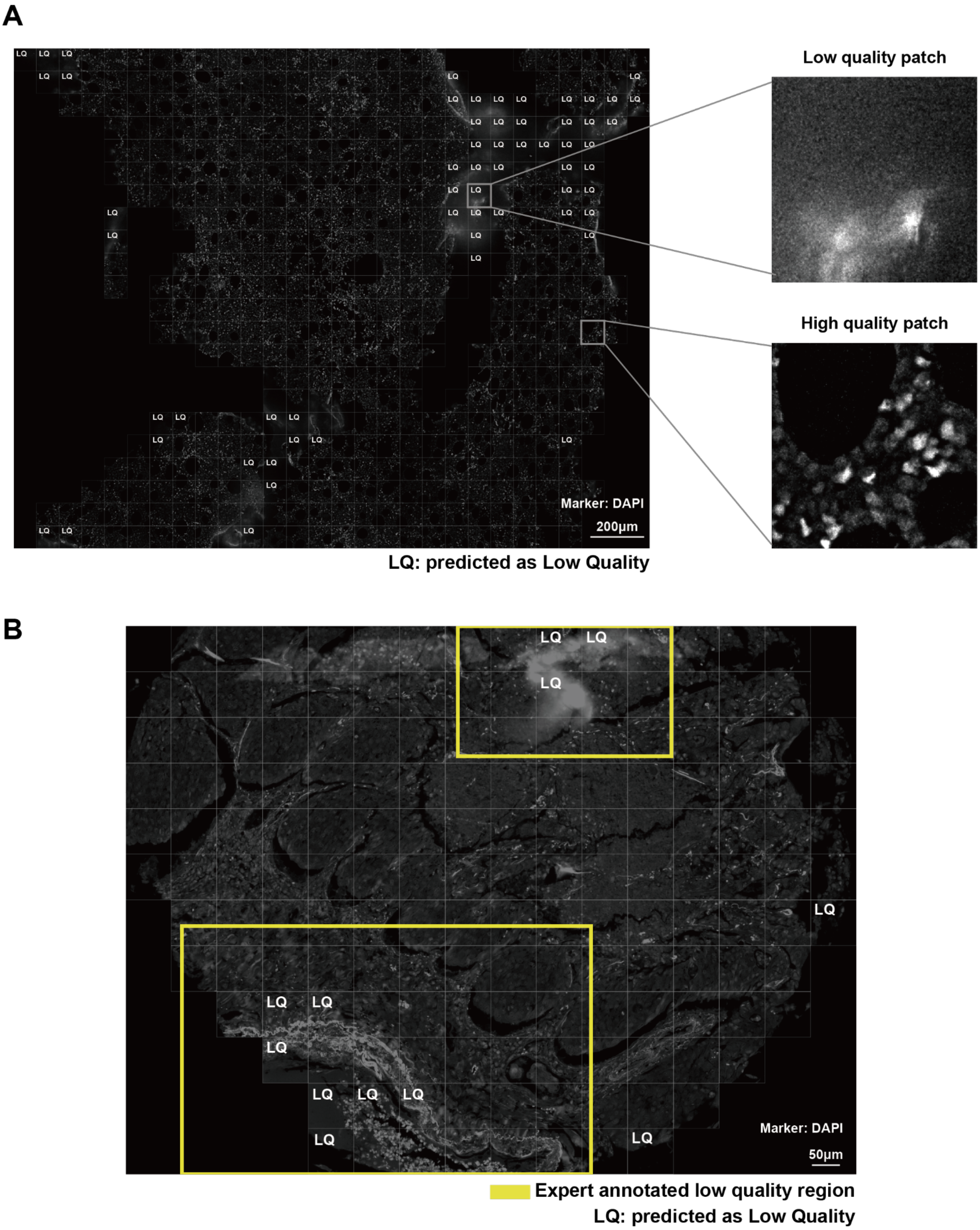


**Supplementary Figure 11. Quality control predictions of image patch NIQE scores by Eva. A**, Predicted patch quality, where patches classified as low quality are labeled as LQ. Zoomed-in views of two representative patches are shown. **B**, Comparison between the predicted patch quality map and expert-annotated low quality regions of interest.

### A Patch quality evaluation with expert annotated artifacts

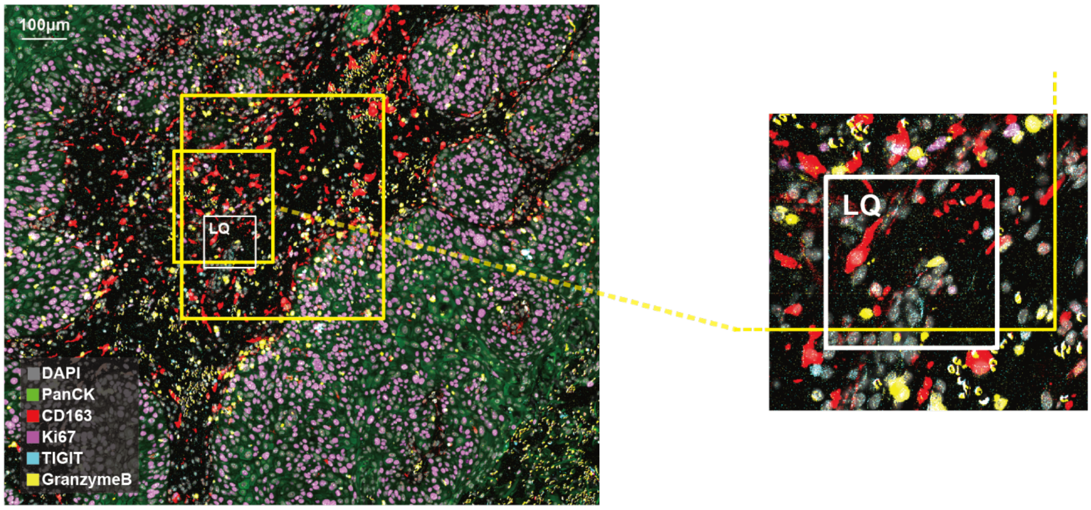


**Expert annotation:** **TIGIT and GranzymeB received scores of 1 (low quality) due to low signal-to-noise ratio and background subtraction artifacts, while Ki67 was scored 2 with relatively higher signal quality.**

### B Patch quality evaluation with expert annotated low quality ROI

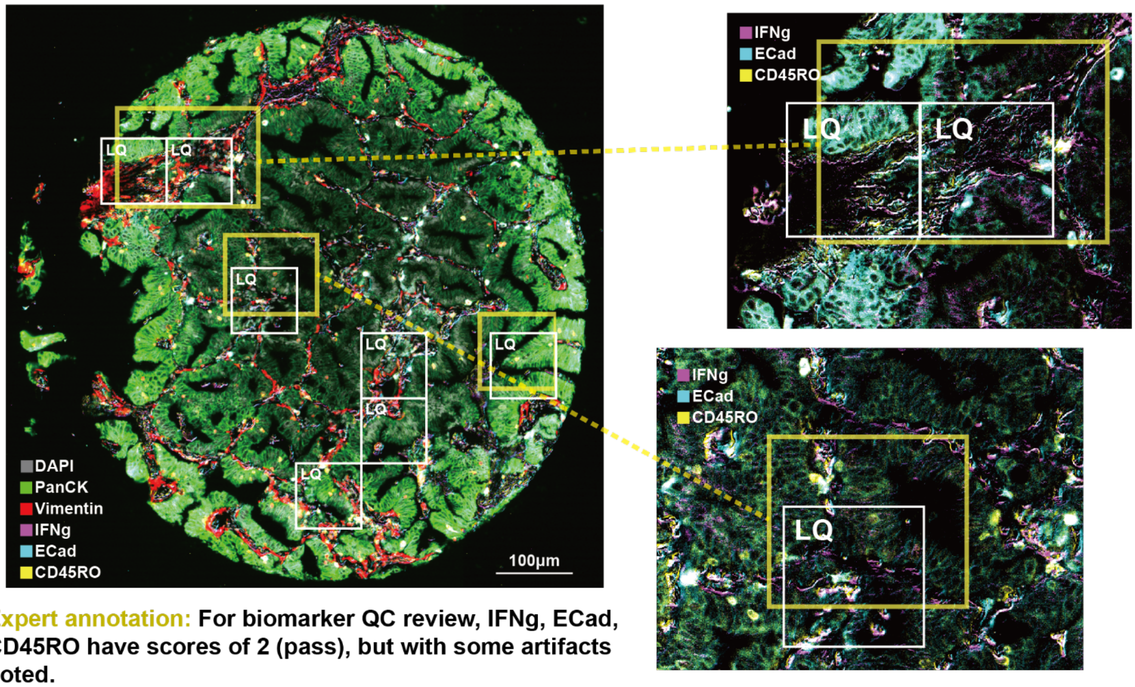


**Expert annotation:** **For biomarker QC review, IFNg, ECad, CD45RO have scores of 2 (pass), but with some artifacts noted.**

**Supplementary Figure 12: Artifact detection with Eva. A** and **B**, Artifacts detected by Eva matching expert annotations. Yellow boxes were expert-drawn regions of interest with detailed annotations. LQ: low quality (artifact detected).

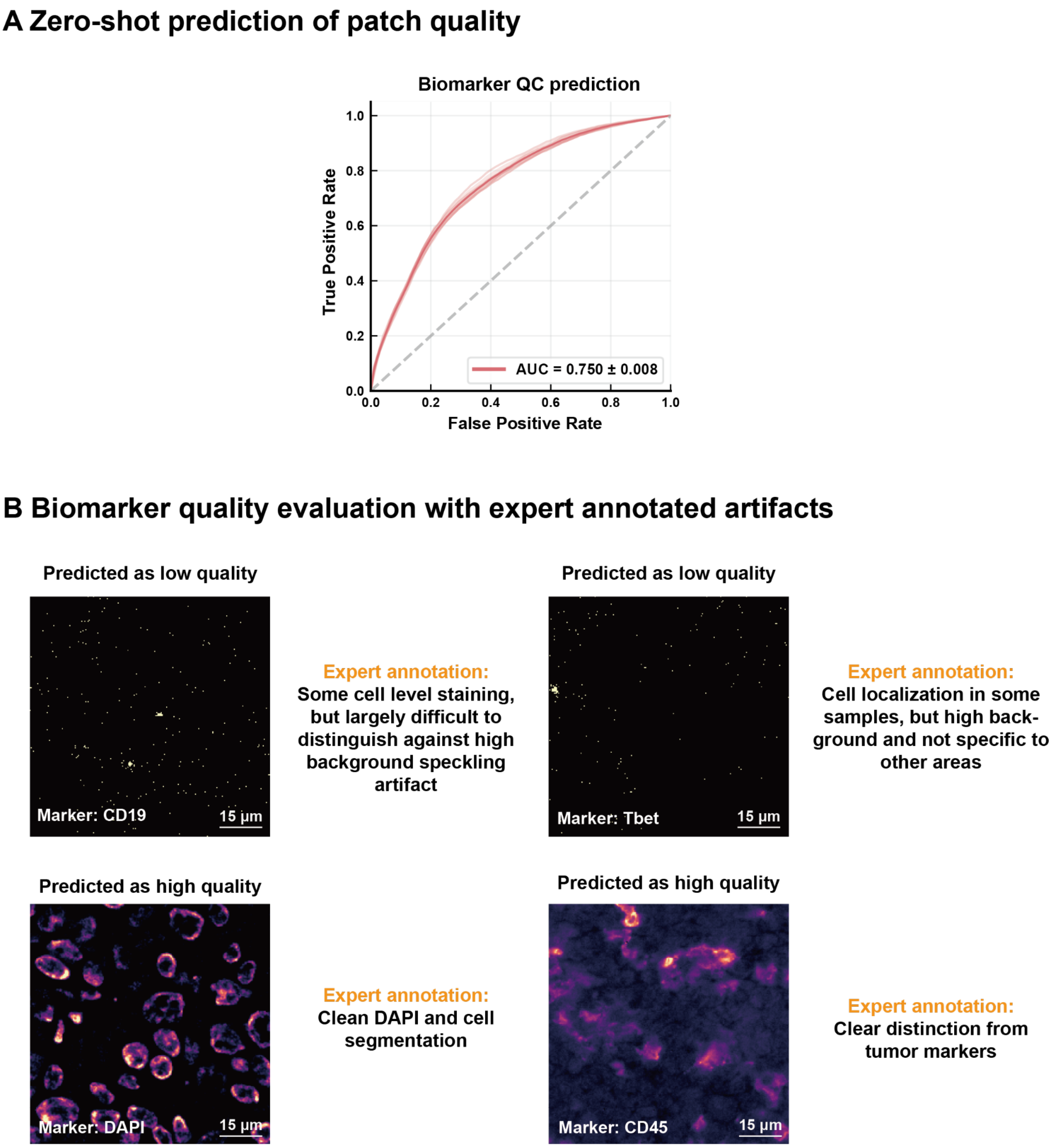


**Supplementary Figure 13: Biomarker quality assessment with Eva. A**, Performance metrics (AUCs) of zero-shot biomarker quality predictions generated by Eva. The models were trained with linear probes using the biomarker QC tasks and data shown in **Figure 3F**. Evaluations were performed on a held-out collection of expert annotations of biomarker qualities. **B**, Visualizations of high-quality and low-quality examples with expert annotations.

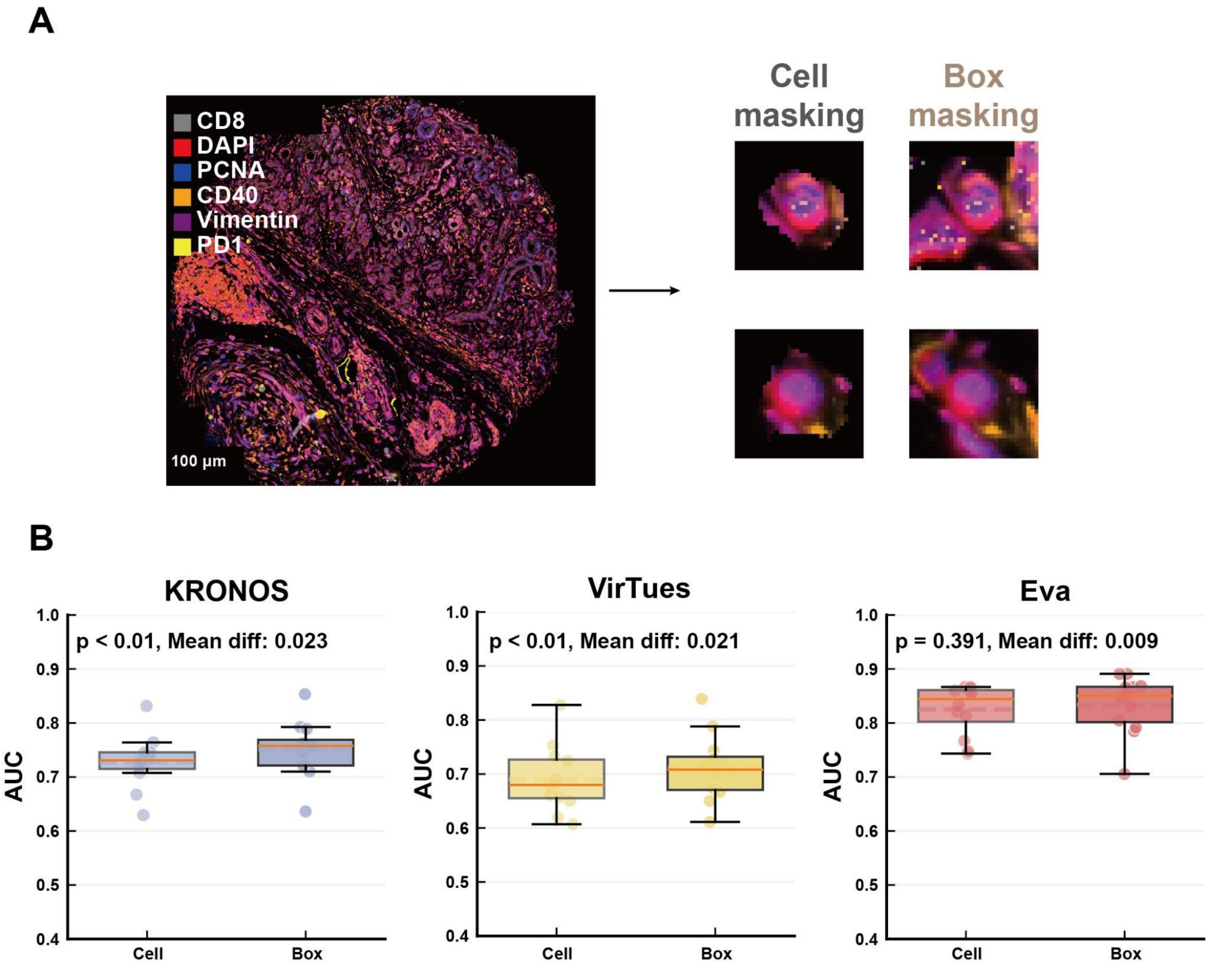


**Supplementary Figure 14. Model performance for cell type prediction using cell masking and box masking. A**, Schematic illustration of the two masking approaches. In cell masking, only pixels within the cell segmentation boundaries are preserved, whereas in box masking, all pixels within a 32 × 32 box centered at the centroid of the cell segmentation are retained. **B**, Boxplot comparison of cell masking and box masking for KRONOS, VirTues, and Eva. Statistical differences between cell masking and box masking performances were assessed using the Wilcoxon signed-rank test on paired AUC values across multiple datasets.

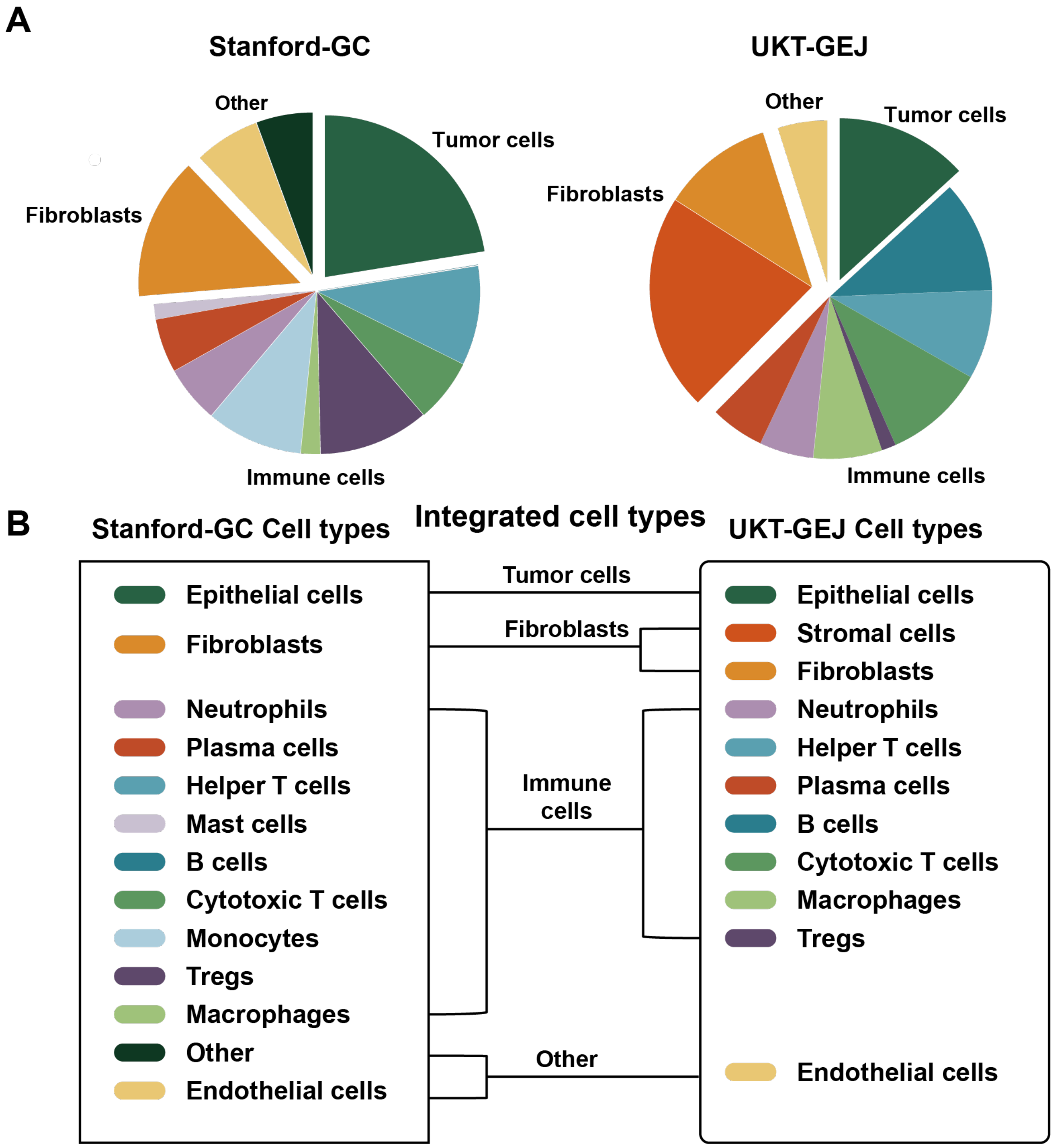


**Supplementary Figure 15. Cell type distribution and unified cell mapping of the Stanford-GC and UKT-GEJ datasets for the zero-shot and few-shot cell label transfer.** **A**, Cell type distributions for both datasets. **B**, Cell type mapping for both datasets, in which unified cell types (tumor cells, fibroblasts, immune cells, and other) are defined for zero-shot and few-shot evaluations.

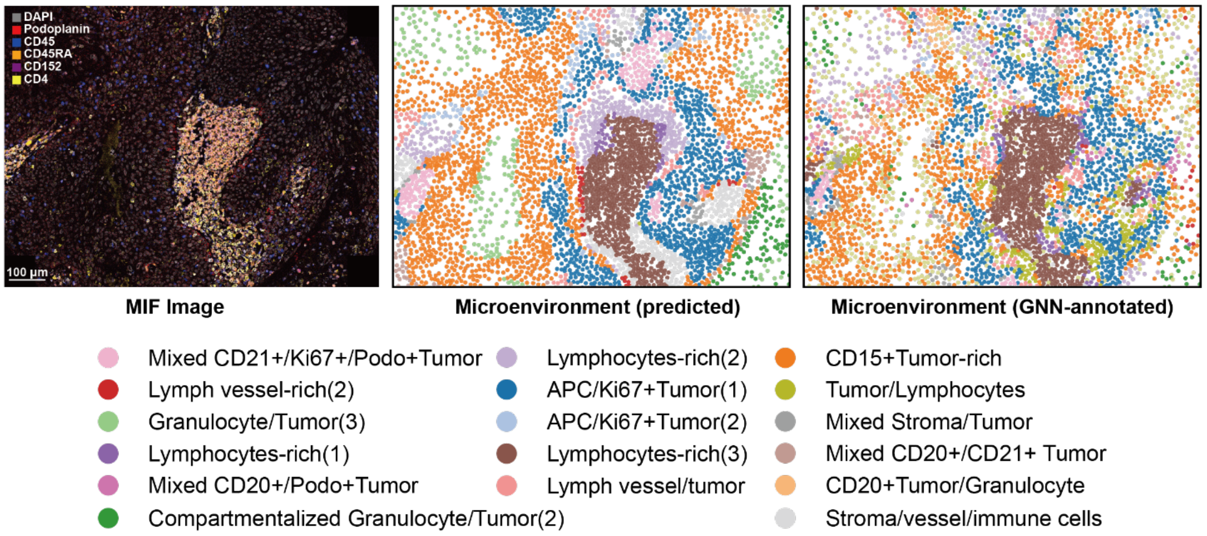


**Supplementary Figure 16. Visualization of microenvironment prediction with Eva.** Microenvironment type prediction is performed on patches centered on the labeled microenvironment centroid. The original MIF image, the predicted microenvironment, and the GNN-annotated microenvironment are shown. GNN: Graph neural networks, proposed by previous work [36].

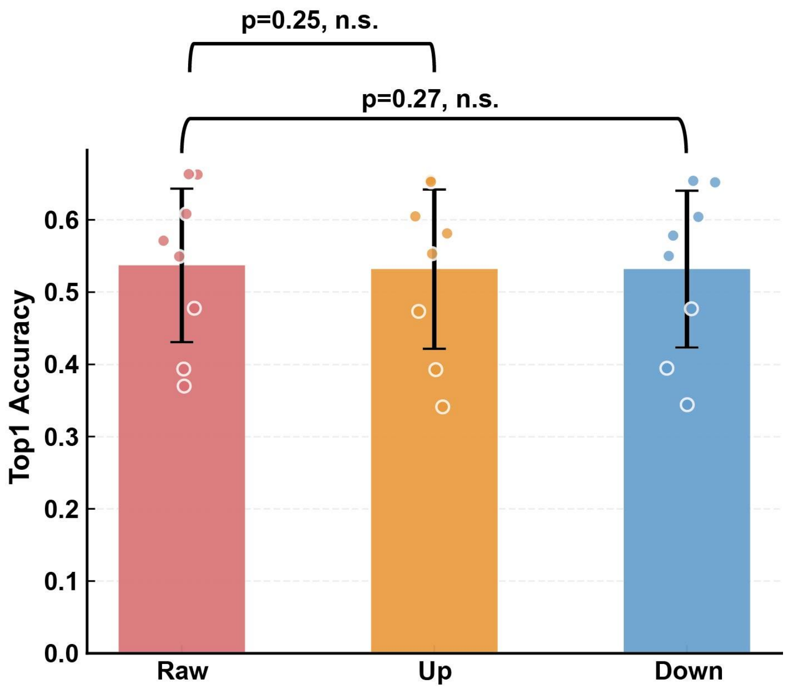


**Supplementary Figure 17. Model robustness to image resolution variations.** Eva's performances were quantified using Top-1 accuracy in the patch-level retrieval task, averaged across multiple datasets. Upsampling was performed by mapping each patch onto a denser grid and then resizing it back to its original dimensions. Downsampling was performed by projecting each patch onto a coarser grid and then expanding it back to its original dimensions. The scaling factor is 2 for upsampling and 0.5 for downsampling. Statistical differences between the metrics were assessed using the Wilcoxon signed-rank test.

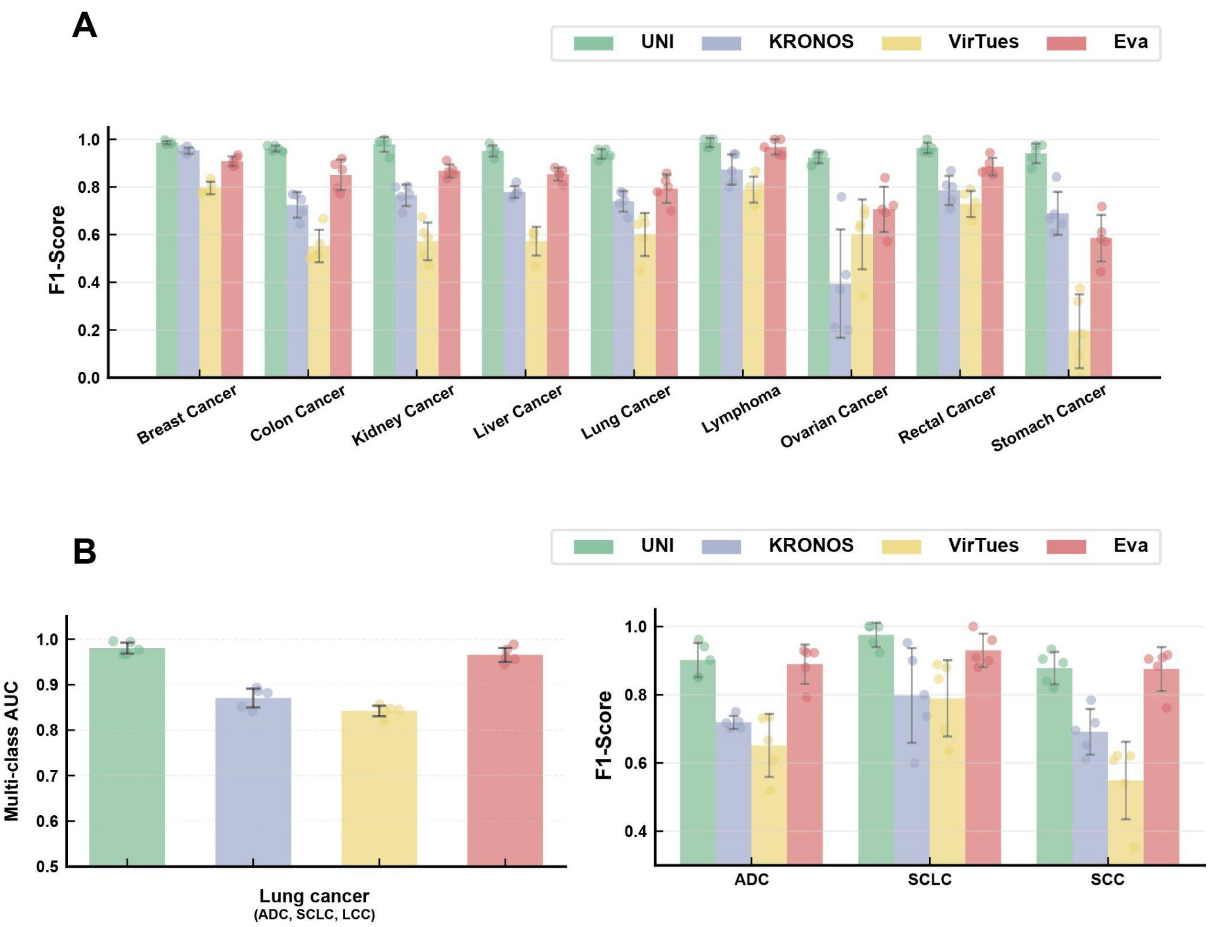


**Supplementary Figure 18. Comparison of foundation models on tumor typing and subtyping tasks. A,** Model performance on tumor type predictions. **B,** Model performance on lung cancer subtype predictions. ADC: Adenocarcinoma, SCLC: Small Cell Lung Cancer, SCC: Squamous Cell Carcinoma. Model performances reported by multi-class AUC and F1-scores were evaluated using 5-fold cross-validation.

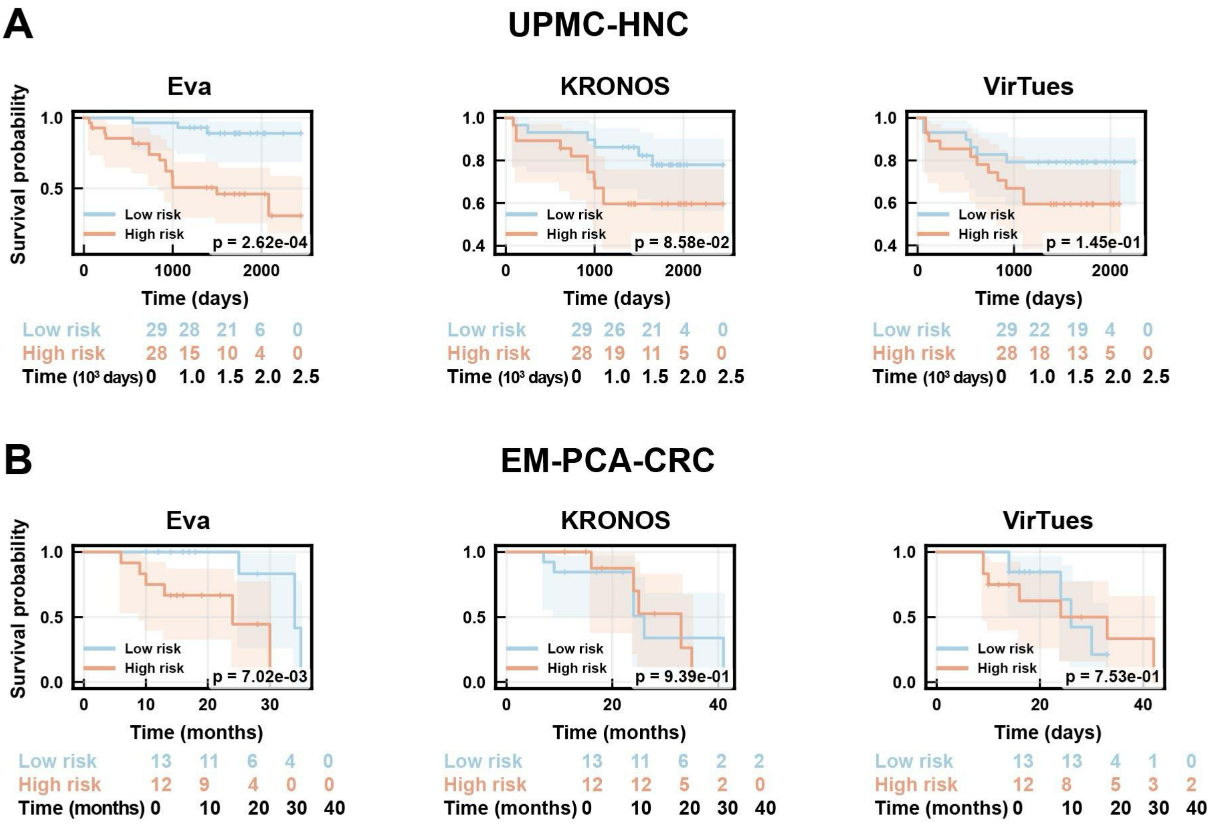


**Supplementary Figure 19. Survival analysis comparison of multiplexed imaging foundation models. A-B**, Kaplan–Meier (KM) curves of patients from the UPMC-HNC and EM-PCA-CRC datasets, bisected into low-risk and high-risk groups based on risks generated by different foundation models. The p-values were calculated using log-rank tests comparing the survival outcomes between the two groups.

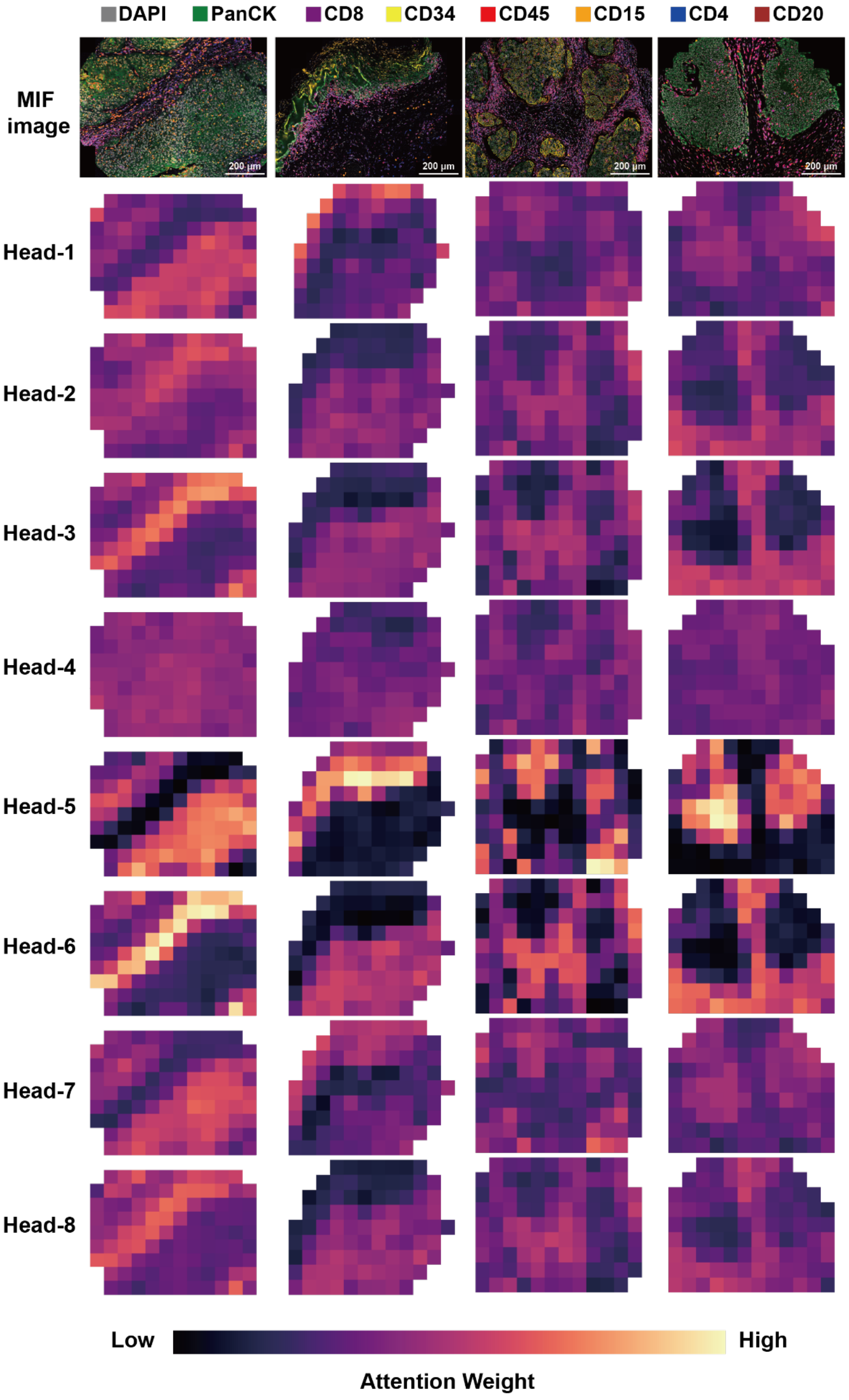


**Supplementary Figure 20: Attention weights for patient stratification extracted from multiple instance learning (MIL).** MIL employed 8 heads, and the computation of attention weights is described in **Methods**.

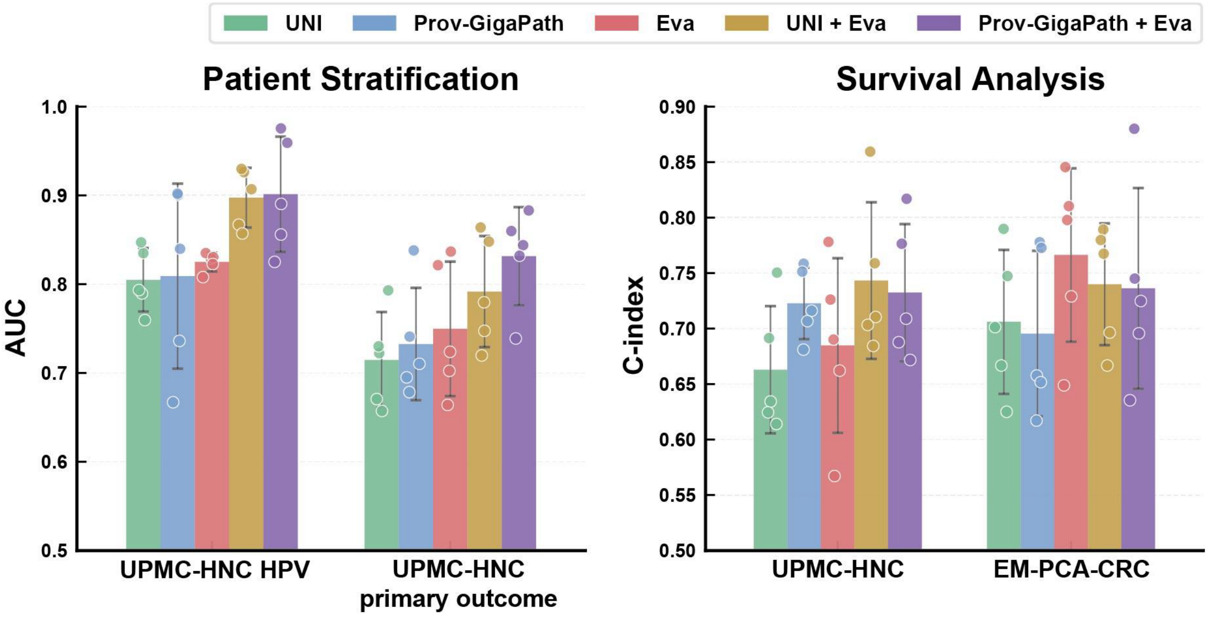


**Supplementary Figure 21. Patient stratification and survival analysis with pathology foundation models (PFMs) and their combinations with Eva. A,** Comparison of patient stratification performances on H&E available dataset (i.e., UPMC-HNC HPV and UPMC-HNC primary outcome) using standard PFMs: UNI and Prov-GigaPath, as well as the MIF-based model Eva. Combined use of embeddings from both modalities (i.e., UNI + Eva, Prov-GigaPath + Eva) yielded the best performance. **B,** Comparison of concordance index for survival analysis on H&E available datasets (i.e., UPMC-HNC and EM-PCA-CRC). Similarly, combined use of both modalities improved upon PFM-only approaches. All tasks were evaluated by 5-fold cross-validation. The combined approaches utilized a late fusion strategy, in which the embeddings extracted by the two distinct models on H&E and MIF data, respectively, were concatenated and evaluated using the same linear probing strategy.

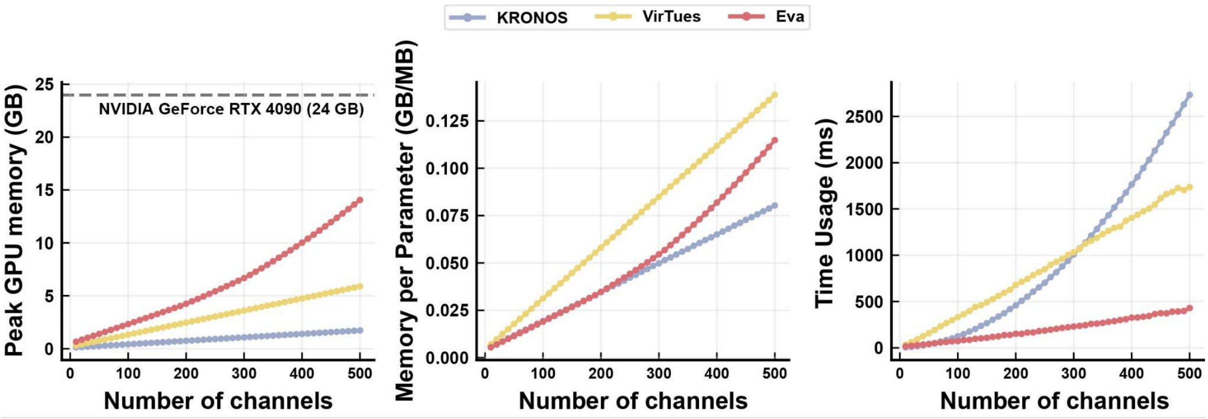


**Supplementary Figure 22: Memory and time usage vary with the number of channels.** Left: Peak GPU memory usage. The dashed line indicates the memory constraint of a consumer-grade 24GB NVIDIA GeForce RTX 4090 GPU. Middle: Memory usage per parameter. Parameter sizes for each model: KRONOS: 21.48 MB, VirTues: 42.45 MB, Eva: 122.44 MB; Right: Time usage, averaged over 50 inference runs. Experiments were conducted on a single 24 GB NVIDIA GeForce RTX 4090 GPU.

### Supplementary tables:

**Supplementary Table1: Biomarker and GenePT gene usage in Eva training.** Data are sorted in alphabetical order.

| Biomarker | Gene | Biomarker | Gene |
|---|---|---|---|
| Arg-1 | ARG1 | aSMA | ACTA2 |
| ATM | ATM | bCatenin | CTNNB1 |
| BCL2 | BCL2 | BCL6 | BCL6 |
| c-Myc | MYC | cPARP-cCasp3 | PARP1 |
| CA9 | CA9 | Caveolin1 | CAV1 |
| CD1c | CD1C | CD2 | CD2 |
| CD3e | CD3E | CD4 | CD4 |
| CD5 | CD5 | CD7 | CD7 |
| CD8 | CD8A | CD11b | ITGAM |
| CD11c | ITGAX | CD14 | CD14 |
| CD15 | FUT4 | CD16 | FCGR3A |
| CD19 | CD19 | CD20 | MS4A1 |
| CD21 | CR2 | CD25 | IL2RA |
| CD27 | CD27 | CD30 | TNFRSF8 |
| CD31 | PECAM1 | CD33 | CD33 |
| CD34 | CD34 | CD36 | CD36 |
| CD38 | CD38 | CD39 | ENTPD1 |
| CD40 | CD40 | CD44 | CD44 |
| CD45 | PTPRC | CD45RA | PTPRC |
| CD45RO | PTPRC | CD47 | CD47 |
| CD49 | ITGA4 | CD56 | NCAM1 |
| CD57 | B3GAT1 | CD62L | SELL |
| CD66 | CEACAM6 | CD66b | CEACAM8 |
| CD68 | CD68 | CD69 | CD69 |
| CD71 | TFRC | CD73 | NT5E |
| CD74 | CD74 | CD79 | CD79A |
| CD86 | CD86 | CD90 | THY1 |
| CD94 | KLRD1 | CD103 | ITGAE |
| CD107a | LAMP1 | CD117 | KIT |
| CD123 | IL3RA | CD127 | IL7R |
| CD134 | TNFRSF4 | CD137 | TNFRSF9 |
| CD138 | SDC1 | CD140b | PDGFRB |
| CD141 | THBD | CD146 | MCAM |
| CD152 | CTLA4 | CD162 | SELPLG |
| CD163 | CD163 | CD183 | CXCR3 |
| CD194 | CCR4 | CD196 | CCR6 |

| | | | |
|---|---|---|---|
| CD197 | CCR7 | CD206 | MRC1 |
| CD207 | CD207 | CD208 | CD207 |
| CD209 | CD209 | CD227 | MUC1 |
| CD271 | NGFR | CD276 | CD276 |
| CDX2 | CDX2 | ChromograninA | CHGA |
| Clusterin | CLU | CollagenI | COL1A1 |
| CollagenIV | COL4A1 | CX3CR1 | CX3CR1 |
| CXCL12 | CXCL12 | CXCL13 | CXCL13 |
| CXCR1 | CXCR1 | CXCR2 | CXCR2 |
| CXCR5 | CXCR5 | DAPI | DAPI |
| ECad | CDH1 | EGFR | EGFR |
| EpCAM | EPCAM | ERa | ESR1 |
| F4/80 | ADGRE1 | FAP | FAP |
| Fibronectin | FN1 | FoxP3 | FOXP3 |
| Gal3 | LGALS3 | GATA3 | GATA3 |
| Geminin | GMNN | GFAP | GFAP |
| GLUT1 | SLC2A1 | GranzymeB | GZMB |
| H3K27me3 | H3F3A | HER2 | ERBB2 |
| HIF1a | HIF1A | HistoneH3 | H3F3A |
| HistoneH3p | H3F3A | HLA-ABC | HLA-A |
| HLA-DR | HLA-DRA | HLA-E | HLA-E |
| ICOS | ICOS | IDO1 | IDO1 |
| IFNg | IFNG | INOS | NOS2 |
| IRF4 | IRF4 | Keratin5 | KRT5 |
| Keratin7 | KRT7 | Keratin8 | KRT8 |
| Keratin8/18 | KRT8 | Keratin10 | KRT10 |
| Keratin14 | KRT14 | Keratin17 | KRT17 |
| Keratin19 | KRT19 | Ki67 | MKI67 |
| LAG3 | LAG3 | LOX-1 | OLR1 |
| Ly6G | LY6G6C | LYVE1 | LYVE1 |
| MCT | SLC16A1 | MMP9 | MMP9 |
| MMP12 | MMP12 | MPO | MPO |
| mTOR | MTOR | Na/K ATPase | ATP1A1 |
| Nestin | NES | NKG2D | KLRK1 |
| NPM1 | NPM1 | Olig2 | OLIG2 |
| OX40 | TNFRSF4 | p-ERK1/2 | MAPK3 |
| p16 | CDKN2A | p53 | TP53 |
| PanCK | KRT | PARP1 | PARP1 |
| pATM | ATM | Pax5 | PAX5 |
| PCNA | PCNA | PD1 | PDCD1 |
| PDL1 | CD274 | PDL2 | PDCD1LG2 |

| | | | |
|---|---|---|---|
| Perforin | PRF1 | Perilipin | PLIN1 |
| Periostin | POSTN | Perlecan | HSPG2 |
| PGP9.5 | UCHL1 | PGR | PGR |
| PNAD | PNAD | Podoplanin | PDPN |
| pS6 | RPS6 | pSTAT3 | STAT3 |
| RAD51 | RAD51 | RORgammaT | RORC |
| RPS6 | RPS6 | S100A4 | S100A4 |
| S100A8/9 | S100A8 | Siglec8 | SIGLEC8 |
| SNAI2 | SNAI2 | SOX2 | SOX2 |
| SOX9 | SOX9 | SOX10 | SOX10 |
| Synaptophysin | SYP | Tbet | TBX21 |
| TCF1 | TCF7 | TCRb | TRB |
| TCRgammadelta | TRGC1 | Tenascin-C | TNC |
| TFAM | TFAM | TIGIT | TIGIT |
| TIM3 | HAVCR2 | TMEM16A | ANO1 |
| TOX | TOX | TP63 | TP63 |
| Tryptase | TPSB2 | TTF1 | NKX2-1 |
| Twist | TWIST1 | Vimentin | VIM |
| VISTA | VSIR | XCR1 | XCR1 |
| yH2AX | H2AX | | |

**Supplementary Table 2: Performance comparison of AUC of cell type prediction tasks.** Masking indicates different types of masking after cell segmentation.

| Dataset | Masking | KRONOS | VirTues | Eva |
|---|---|---|---|---|
| Stanford-GC | cell | 0.7640 | 0.7526 | **0.8591** |
| | box | 0.7926 | 0.7882 | **0.8911** |
| EM-Mixed | cell | 0.7313 | 0.6786 | **0.8211** |
| | box | 0.7458 | 0.7040 | **0.8317** |
| Stanford-GIST | cell | 0.6673 | 0.7329 | **0.7433** |
| | box | 0.7099 | 0.7436 | **0.7847** |
| Stanford-PC | cell | 0.7305 | 0.6610 | **0.8598** |
| | box | 0.7615 | 0.7012 | **0.8664** |
| MDACC-HCC | cell | 0.7335 | 0.6899 | **0.8667** |
| | box | 0.7571 | 0.7124 | **0.8657** |
| UKT-GEJ | cell | 0.7457 | 0.6808 | **0.8643** |
| | box | 0.7622 | 0.7207 | **0.8685** |
| UPMC-HNC | cell | 0.7077 | 0.6197 | **0.8661** |
| | box | 0.7581 | 0.6717 | **0.8910** |
| Stanford-HCC | cell | **0.8314** | 0.8277 | 0.7666 |
| | box | **0.8534** | 0.8390 | 0.8049 |
| Stanford-CRC | cell | 0.7174 | 0.6502 | **0.8553** |
| | box | 0.7209 | 0.6666 | **0.8469** |
| MDACC-MM | cell | 0.6294 | 0.6069 | **0.7474** |
| | box | 0.6362 | 0.6112 | **0.7056** |
| IUCPQ-LUAD | cell | 0.7276 | 0.6568 | **0.8329** |
| | box | 0.7216 | 0.6499 | **0.7915** |
| RAHBT-DCIS | cell | 0.7456 | 0.7245 | **0.8143** |
| | box | 0.7889 | 0.7281 | **0.8539** |

**Supplementary Table 3: Performance comparison of AUC of microenvironment prediction tasks.**

| Dataset | KRONOS | VirTues | Eva |
|---|---|---|---|
| UPMC-HNC | 0.8295 | 0.8421 | **0.8842** |
| MDACC-HCC | 0.8594 | 0.8300 | **0.8915** |

**Supplementary Table 4: Performance comparison of PCC of cell composition task, higher the better.**

| Dataset | KRONOS | VirTues | Eva |
|---|---|---|---|
| UPMC-HNC | 0.6287 | 0.6768 | **0.7807** |
| Stanford-GC | 0.6351 | 0.6445 | **0.7420** |
| MDACC-HCC | 0.6615 | 0.6564 | **0.7531** |
| Stanford-PC | 0.5855 | 0.6308 | **0.7549** |
| Stanford-GIST | 0.5495 | 0.5780 | **0.6625** |
| UKT-GEJ | 0.6138 | 0.6490 | **0.7450** |
| EM-Mixed | 0.5881 | 0.5988 | **0.7113** |
| Stanford-HCC | 0.5161 | 0.5438 | **0.6187** |
| Stanford-CRC | 0.4798 | 0.5510 | **0.6035** |
| MDACC-MM | 0.1616 | 0.1960 | **0.2674** |
| IUCPQ-LUAD | 0.3876 | 0.3761 | **0.4349** |
| RAHBT-DCIS | 0.2093 | 0.2702 | **0.2996** |

**Supplementary Table 5: Performance comparison of MSE of cell composition task, lower the better.**

| Dataset | KRONOS | VirTues | Eva |
|---|---|---|---|
| UPMC-HNC | 0.0048 | 0.0045 | **0.0031** |
| Stanford-GC | 0.0090 | 0.0096 | **0.0063** |
| MDACC-HCC | 0.0063 | 0.0071 | **0.0054** |
| Stanford-PC | 0.0089 | 0.0081 | **0.0055** |
| Stanford-GIST | 0.0164 | 0.0152 | **0.0133** |
| UKT-GEJ | 0.0099 | 0.0095 | **0.0063** |
| EM-Mixed | 0.0062 | 0.0065 | **0.0038** |
| Stanford-HCC | 0.0295 | 0.0281 | **0.0209** |
| Stanford-CRC | 0.0171 | 0.0138 | **0.0113** |
| MDACC-MM | 0.0030 | **0.0025** | **0.0025** |
| IUCPQ-LUAD | 0.0029 | 0.0027 | **0.0018** |
| RAHBT-DCIS | 0.0098 | 0.0083 | **0.0068** |

**Supplementary Table 6: Performance comparison of Top-K accuracy of retrieval based on MIF patches.** Top-K accuracy is computed based on the dominant cell type of the patch, higher the better.

| Dataset | Top-K | KRONOS | VirTues | Eva |
|---|---|---|---|---|
| Stanford-GIST | 1 | 0.4336 | 0.4051 | **0.4776** |
| | 3 | 0.6419 | 0.6275 | **0.6872** |
| | 5 | 0.7222 | 0.7173 | **0.7621** |
| Stanford-PC | 1 | 0.5078 | 0.5019 | **0.6082** |
| | 3 | 0.7305 | 0.7228 | **0.8067** |
| | 5 | 0.8082 | 0.8026 | **0.8702** |
| UPMC-HNC | 1 | 0.2725 | 0.2637 | **0.3935** |
| | 3 | 0.4885 | 0.4814 | **0.6269** |
| | 5 | 0.5976 | 0.5921 | **0.7263** |
| Stanford-HCC | 1 | 0.2957 | 0.2441 | **0.3699** |
| | 3 | 0.4789 | 0.3939 | **0.5721** |
| | 5 | 0.5595 | 0.4761 | **0.6541** |
| MDACC-HCC | 1 | 0.5210 | 0.3777 | **0.5493** |
| | 3 | 0.6730 | 0.5402 | **0.7274** |
| | 5 | 0.7392 | 0.6147 | **0.8005** |
| EM-Mixed | 1 | 0.5568 | 0.5339 | **0.6627** |
| | 3 | 0.7193 | 0.6904 | **0.8113** |
| | 5 | 0.7842 | 0.7484 | **0.8613** |
| Stanford-GC | 1 | 0.5543 | 0.5336 | **0.6632** |
| | 3 | 0.7507 | 0.7225 | **0.8255** |
| | 5 | 0.8153 | 0.7921 | **0.8817** |
| UKT-GEJ | 1 | 0.5011 | 0.4960 | **0.5711** |
| | 3 | 0.7260 | 0.7019 | **0.7809** |
| | 5 | 0.7993 | 0.7789 | **0.8521** |
| IUCPQ-LUAD | 1 | 0.6880 | 0.6446 | **0.7546** |
| | 3 | 0.8301 | 0.7887 | **0.8825** |
| | 5 | 0.8694 | 0.8387 | **0.9131** |
| RAHBT-DCIS | 1 | 0.3747 | 0.4101 | **0.4791** |
| | 3 | 0.5484 | 0.6078 | **0.7242** |
| | 5 | 0.6429 | 0.7117 | **0.8132** |

**Supplementary Table 7: Performance comparison of Top-K MSE of retrieval based on MIF patches.** MSE is computed as the nearest embedding distance in queried patches, lower the better.

| Dataset | Top-K | KRONOS | VirTues | Eva |
|---|---|---|---|---|
| Stanford-GIST | 1 | 0.0269 | 0.0274 | **0.0234** |
| | 3 | 0.0161 | 0.0164 | **0.0137** |
| | 5 | 0.0127 | 0.0129 | **0.0108** |
| Stanford-PC | 1 | 0.0150 | 0.0147 | **0.0103** |
| | 3 | 0.0083 | 0.0083 | **0.0059** |
| | 5 | 0.0065 | 0.0065 | **0.0046** |
| UPMC-HNC | 1 | 0.0103 | 0.0102 | **0.0072** |
| | 3 | 0.0067 | 0.0065 | **0.0046** |
| | 5 | 0.0052 | 0.0054 | **0.0038** |
| Stanford-HCC | 1 | 0.0824 | 0.0934 | **0.0670** |
| | 3 | 0.0530 | 0.0655 | **0.0431** |
| | 5 | 0.0428 | 0.0535 | **0.0347** |
| MDACC-HCC | 1 | 0.0147 | 0.0210 | **0.0130** |
| | 3 | 0.0092 | 0.0134 | **0.0078** |
| | 5 | 0.0075 | 0.0110 | **0.0062** |
| EM-Mixed | 1 | 0.0141 | 0.0135 | **0.0088** |
| | 3 | 0.0082 | 0.0081 | **0.0051** |
| | 5 | 0.0067 | 0.0067 | **0.0042** |
| Stanford-GC | 1 | 0.0171 | 0.0181 | **0.0112** |
| | 3 | 0.0091 | 0.0101 | **0.0057** |
| | 5 | 0.0067 | 0.0078 | **0.0042** |
| UKT-GEJ | 1 | 0.0174 | 0.0171 | **0.0133** |
| | 3 | 0.0096 | 0.0097 | **0.0075** |
| | 5 | 0.0075 | 0.0078 | **0.0060** |
| IUCPQ-LUAD | 1 | 0.0041 | 0.0049 | **0.0025** |
| | 3 | 0.0022 | 0.0028 | **0.0015** |
| | 5 | 0.0018 | 0.0022 | **0.0013** |
| RAHBT-DCIS | 1 | 0.0156 | 0.0154 | **0.0122** |
| | 3 | 0.0098 | 0.0080 | **0.0062** |
| | 5 | 0.0074 | 0.0059 | **0.0046** |

**Supplementary Table 8: Performance of F1-score of tumor type classification task.**

| Tumor type | UNI | KRONOS | VirTues | Eva |
|---|---|---|---|---|
| Breast Cancer | **0.9863±0.0071** | 0.9523±0.0129 | 0.7958±0.0262 | 0.9081±0.0202 |
| Colon Cancer | **0.9610±0.0122** | 0.7248±0.0538 | 0.5521±0.0682 | 0.8508±0.0642 |
| Kidney Cancer | **0.9783±0.0312** | 0.7651±0.0453 | 0.5716±0.0797 | 0.8673±0.0281 |
| Liver Cancer | **0.9516±0.0235** | 0.7789±0.0251 | 0.5724±0.0602 | 0.8535±0.0270 |
| Lung Cancer | **0.9388±0.0205** | 0.7397±0.0434 | 0.6004±0.0906 | 0.7926±0.0594 |
| Lymphoma | **0.9862±0.0189** | 0.8730±0.0635 | 0.7891±0.0547 | 0.9677±0.0323 |
| Ovarian Cancer | **0.9228±0.0230** | 0.3944±0.2271 | 0.6013±0.1467 | 0.7061±0.0951 |
| Rectal Cancer | **0.9636±0.0223** | 0.7858±0.0614 | 0.7289±0.0551 | 0.8852±0.0371 |
| Stomach Cancer | **0.9407±0.0410** | 0.6895±0.0902 | 0.1944±0.1551 | 0.5851±0.0979 |

**Supplementary Table 9: Comparison of F1-score of lung cancer subtype classification task.**

| Lung cancer subtype | UNI | KRONOS | VirTues | Eva |
|---|---|---|---|---|
| ADC | **0.9015±0.0504** | 0.7188±0.0194 | 0.6513±0.0911 | 0.8895±0.0576 |
| SCLC | **0.9751±0.0356** | 0.7978±0.1389 | 0.7894±0.1157 | 0.9298±0.0491 |
| SCC | **0.8780±0.0475** | 0.6916±0.0667 | 0.5487±0.1146 | 0.8752±0.0645 |

ADC: Adenocarcinoma; SCLC: Small Cell Lung Carcinoma; SCC: Squamous Cell Carcinoma

**Supplementary Table 10: Comparison of C-index of survival analysis.**

| Dataset | KRONOS | VirTues | Eva |
|---|---|---|---|
| UPMC-HNC | 0.6350±0.0526 | 0.6089±0.0655 | **0.6847±0.0788** |
| EM-PCA-CRC | 0.6444±0.1478 | 0.6508±0.0852 | **0.7663±0.0781** |

**Supplementary Table 11: Performance comparison of AUC of patient classification tasks.**

| Dataset | KRONOS | VirTues | Eva |
|---|---|---|---|
| UPMC-HNC HPV | 0.7371±0.0374 | 0.7583±0.0897 | **0.8250±0.0105** |
| UPMC-HNC PO | 0.6540±0.1365 | 0.7169±0.0844 | **0.7497±0.0758** |
| MDACC-MM response | 0.7154±0.0283 | 0.6780±0.0545 | **0.7774±0.0619** |
| UHB-CRC GA | 0.7311±0.0309 | 0.6890±0.0886 | **0.7884±0.1200** |
| UHB-CRC KM | 0.6135±0.1420 | 0.5887±0.2114 | **0.7413±0.0781** |
| Stanford-CRC PO | 0.4692±0.0728 | 0.4985±0.0134 | **0.6091±0.1079** |

**Supplementary Table 12: Performance comparison of Top-1 accuracy of zero-shot case-level retrieval tasks.**

| Dataset | Random | KRONOS | VirTues | Eva |
|---|---|---|---|---|
| UPMC-HNC HPV | 0.3926 | 0.6239 | 0.6358 | **0.6606** |
| UPMC-HNC PO | 0.4175 | 0.5719 | 0.5994 | **0.6636** |
| MDACC-MM response | 0.5062 | 0.6111 | 0.6326 | **0.7742** |
| UHB-CRC GA | 0.3337 | 0.6570 | 0.6280 | **0.7246** |
| UHB-CRC KM | 0.3688 | 0.6763 | 0.6908 | **0.7101** |
| DFCI-HNC pTR | 0.3732 | 0.4474 | **0.5702** | 0.5356 |
| MDACC-HCC response | 0.4569 | 0.5294 | 0.4706 | **0.7059** |

**Supplementary Table 13: Performance comparison of Eva, PFMs and late fusion ensembles on downstream tasks.**

| Task type | Dataset | Eva | UNI | Prov-GigaPath | Eva+UNI | Eva+ Prov-GigaPath |
|---|---|---|---|---|---|---|
| Case-level (AUC) | UPMC-HNC HPV | 0.8250±0.0105 | 0.8049±0.0358 | 0.8090±0.1042 | 0.8974±0.0336 | 0.9012±0.0649 |
| | UPMC-HNC PO | 0.7497±0.0758 | 0.7146±0.0540 | 0.7325±0.0633 | 0.7917±0.0625 | 0.8315±0.0552 |
| Survival (C-index) | UPMC-HNC | 0.6847±0.0788 | 0.6629±0.0573 | 0.7227±0.0321 | 0.7432±0.0705 | 0.7323±0.0619 |
| | EM-PCA-CRC | 0.7663±0.0781 | 0.7060±0.0649 | 0.6954±0.0745 | 0.7399±0.0548 | 0.7362±0.0904 |

**Supplementary Table 14: Dataset and model parameters used in Eva pretraining.**

| Parameter | Value |
|---|---|
| Batch size | 16 |
| Patch size | 224 |
| Token size | 8 |
| Marker embedding | GenePT |
| Embedding dimension | 3072 |
| Masking strategy | random |
| Masking ratio | 0.75 |
| Epochs | 20 |
| Warmup epochs | 5 |
| Start learning rate | $10^{-4}$ |
| Warmup end learning rate | $10^{-4}$ |
| Final learning rate | $10^{-5}$ |
| Learning rate schedule | Cosine |
| CE dimension | 512 |
| CE MLP ratio | 4 |
| CE heads | 4 |
| CE layers | 2 |
| CE output dimension | 512 |
| TE dimension | 768 |
| TE MLP ratio | 4 |
| TE heads | 12 |
| TE layers | 12 |
| TE output dimension | 512 |
| DE dimension | 512 |
| DE MLP ratio | 4 |
| DE heads | 16 |
| DE layers | 8 |

CE: Channel-level Encoder; TE: Token-level Encoder; DE:Decoder
